# An epigenetic memory at the *CYP1A* gene in cancer-resistant, pollution-adapted killifish

**DOI:** 10.1101/2024.08.14.607951

**Authors:** Samantha Carrothers, Rafael Trevisan, Nishad Jayasundara, Nicole Pelletier, Emma Weeks, Joel N. Meyer, Richard Di Giulio, Caren Weinhouse

## Abstract

Human exposure to polycyclic aromatic hydrocarbons (PAH) is a significant and growing public health problem. Frequent, high dose exposures are likely to increase due to a warming climate and increased frequency of large-scale wildfires. Here, we characterize an epigenetic memory at the *cytochrome P450 1A* (*CYP1A*) gene in a population of wild *Fundulus heteroclitus* that has adapted to chronic, extreme PAH pollution. In wild-type fish, *CYP1A* is highly induced by PAH. In PAH-tolerant fish, *CYP1A* induction is blunted. Since CYP1A metabolically activates PAH, this memory protects these fish from PAH-mediated cancer. However, PAH-tolerant fish reared in clean water recover *CYP1A* inducibility, indicating that blunted induction is a non-genetic memory of prior exposure. To explore this possibility, we bred depurated wild fish from PAH-sensitive and -tolerant populations, manually fertilized exposure-naïve embryos, and challenged them with PAH. We observed epigenetic control of the reversible memory of generational PAH stress in F_1_ PAH-tolerant embryos. Specifically, we observed a bivalent domain in the *CYP1A* promoter enhancer comprising both activating and repressive histone post-translational modifications. Activating modifications, relative to repressive ones, showed greater increases in response to PAH in sensitive embryos, relative to tolerant, consistent with greater gene activation. Also, PAH-tolerant adult fish showed persistent induction of *CYP1A* long after exposure cessation, which is consistent with defective *CYP1A* shutoff and recovery to baseline. Since *CYP1A* expression is inversely correlated with cancer risk, these results indicate that PAH-tolerant fish have epigenetic protection against PAH-induced cancer in early life that degrades in response to continuous gene activation.

**Significance:** Epigenetic memory, or the inheritance across cell division within an organism or across generations, of environmental exposure response is a compelling phenomenon with limited understanding of mechanism. Here, we characterized an epigenetic memory at the *CYP1A* gene in pollution-adapted *Fundulus heteroclitus*. We found that the *CYP1A* promoter enhancer contains a bivalent domain, comprising both active and repressive histone modifications, that shows reduced function correlating with reduced gene induction by its pollutant activator. In early life, this memory protects fish against pollution-induced cancer. However, this reduced function carries a cost; adult fish show defective transcriptional recovery of *CYP1A*, which increases cancer risk later in life. These results provide an initial mechanism for a model epigenetic memory and highlight potential costs.

## Introduction

Human exposure to polycyclic aromatic hydrocarbons (PAHs) is a significant and growing public health problem(1). PAHs are byproducts of organic combustion that are present in high levels in cigarette smoke, coal-fired power plant emissions, vehicular exhaust, and wildfire smoke(1–5). Wildfire smoke is highly mutagenic; this mutagenicity is strongly correlated with PAH content in both particulate and gas phases, due to the highly uncontrolled, inefficient combustion of biomass that occurs during wildfires(6). Large-scale wildfires lead to high-dose PAH exposures to human populations(7). These exposure events are increasing as the climate warms; the United Nations Environment Programme predicts a global increase in extreme fires of 14% by 2030, 30% by 2050, and 50% by the end of the century(7).

Sustained or extreme exposure to certain PAHs causes lung cancer in humans(2, 4–6, 8–10) and environmental PAH exposures are responsible for most lung cancer cases worldwide(11, 12). A recent study in The Lancet Global Health labeled lung cancer the leading contributor to preventable death in high income countries(11, 12). Tobacco use is the largest environmental risk factor for lung cancer(13–16), followed closely by air pollution(16, 17). In addition, lung cancer is a common disease with poor outcomes. Lung cancer is the most common cancer (nearly 2.5 million cases per year, as of 2020)(18–20) and the leading cause of cancer deaths globally (>1.8 million deaths in 2020)(18–20). In the United States, lung cancers rank second in new cancer case types and first in cancer cases in both sexes(21); treatment outcomes are poor, as indicated by a five-year survival rate of 23% as of 2018(21). Population-level solutions for reducing lung cancer burden involve effective smoking cessation campaigns and policies to ensure clean air and prevent wildfires. However, policy approaches are ideally complemented by personalized exposure mitigation and treatment strategies since individuals show variation in carcinogenic responses to PAHs.

Genotoxic PAHs cause cancer by increasing the frequency of DNA mutations(1). These PAHs do not cause mutations in their parent forms; parent PAHs are pro-carcinogens that are transformed into mutagenic metabolites when bioactivated by metabolic enzymes(1). In mammals, the primary enzyme that bioactivates PAHs is cytochrome P4501A1 (CYP1A1)(22–25). CYP1A1 is a monooxygenase that adds a reactive hydroxyl group or epoxide to PAHs(22–25). When this hydroxyl group reacts further with a glutathione or glucuronide molecule, the resulting conjugated compound is rendered transportable out of the cell for excretion from the body(22, 23). However, reactive PAH metabolites can also form DNA adducts which, if unrepaired, can lead to DNA mutations(22–25). High mutation burdens increase the probability of cancer driver mutations that can transform healthy cells into cancerous ones(22, 23). As a result, CYP1A1 activity is both required for PAH clearance and responsible for PAHs’ carcinogenicity(22, 23). *CYP1A1* gene expression is substantially induced in the presence of PAHs through ligand-dependent activation of the aryl hydrocarbon receptor (AHR), speeding xenobiotic clearance (22, 23, 26, 27). Therefore, as the dose of PAHs increases, so does the DNA mutation burden and cancer risk, due in part to proportionally greater *CYP1A1* expression(28–30). Although the various types of PAHs are metabolized to reactive intermediates that vary in mutagenic potency(28–30), the cancer risk of a given PAH, and, by extension, a given mixture of PAHs found in real-world exposures, largely depends on how strongly it induces *CYP1A1* expression(31, 32).

*CYP1A1* induction predicts PAH-induced cancer risk. PAH exposure and cancer are strongly associated, but PAH-related cancer risk is highly variable among individuals. For example, tobacco smoking is an established risk factor for lung cancer, but there is considerable unexplained variation in risk among smokers(33, 34). Genetic polymorphisms in metabolic genes may explain some of this variation(35–39). Individuals that carry one of four well-characterized mutations in *CYP1A1* are at higher risk for lung cancer(35–39) and, in some ethnic groups, this risk is compounded in carriers that smoke(38, 40–42). These genetic variants likely confer increased cancer risk by increasing formation of mutagenic metabolites, either through increased enzymatic activity or through increasing inducibility of the *CYP1A1* gene in the presence of PAH(43–47). *CYP1A1*’s role in lung cancer development is substantiated further by studies in cancer tissue. *CYP1A1* expression is higher in human lung cancer tissue than in non-cancerous lung tissue(48), is associated with a history of tobacco smoking(49, 50), and has been proposed as a diagnostic marker for lung cancer(22). These findings suggest that genetic or environmental factors that cause sustained increases in *CYP1A1* expression are prevalent and tractable risk factors for lung cancer. Since global lung cancer rates are likely to rise with rising wildfire smoke pollution, we hypothesize that preventing these sustained increases in *CYP1A1* expression will mitigate increases in lung cancer rates.

To better understand how to prevent sustained increases in *CYP1A1* expression in human populations, we focus on a wild fish population that exhibits naturally depressed induction of this metabolic gene(51–55). Specifically, we leverage a natural experiment in a population of wild mummichog (*Fundulus heteroclitus*) in the mainstem and tributaries of the Elizabeth River, Virginia(56). This population of fish has shown rapid, evolutionary adaptation to extreme exposures to PAHs derived from creosote(56, 57). The adaptive phenotype includes resistance to acute toxicity, developmental abnormalities, and liver cancer that occur in wild-type fish exposed to similar doses of PAHs(56, 58). Notably, cancer resistance in these fish results from blunted induction by PAHs of the gene encoding the fish’s single *CYP1A* isoform, *cytochrome P450 1A* (*CYP1A*)(58). In PAH-tolerant fish, PAH exposure still triggers upregulation of *CYP1A*, but to a substantially lesser degree than in wild-type, PAH-sensitive fish(51–55).

Because resistance to *CYP1A* induction occurs in the same population that resists PAH-induced teratogenesis, both phenotypes were thought to result from adaptive downregulation of AHR signaling in PAH-tolerant fish(51, 52, 54, 55). However, several lines of evidence support the conclusion that these two phenotypes are distinct. First, the resistance to *CYP1A* induction by PAH in pollution-tolerant mummichog is epigenetic (i.e., heritable yet non-genetic)(52, 54, 55). The blunted *CYP1A* response persists in first-generation embryos and larvae of wild-caught, PAH-tolerant fish bred in clean water in the laboratory(52). However, gene inducibility partially recovers in first-generation adults and second-generation embryos(52, 55); third-generation embryos show complete recovery to wild-type levels(52). In contrast, resistance to PAH-induced developmental abnormalities is inherited unchanged across generations of PAH-tolerant fish(56, 57). Second, PAH developmental toxicity partially requires AHR signaling but does not require downstream *CYP1A* expression(59). When AHR is induced and CYP1A is inhibited simultaneously in PAH-tolerant embryos, embryos showed increased levels of developmental abnormalities, likely due to decreased PAH clearance in the absence of active CYP1A(59). Finally, blunted *CYP1A* induction by PAH is not linked to increased mummichog larval survival or greater reproductive fitness, indicating that blunted *CYP1A* inducibility is not an evolutionarily adaptive trait in PAH-tolerant fish(52, 60).

Therefore, we hypothesized that the transient resistance of *CYP1A* to PAH induction reflects an epigenetic memory of past PAH exposure that formed independently of the genetic adaptation to PAH-related teratogenesis. This phenotype is not a result of maternal loading of PAHs(60) nor is it due to maternal effects(60). In this study, we show evidence of epigenetic control of *CYP1A* response to PAH in pollution adapted *Fundulus heteroclitus*. Specifically, we show that the promoter-enhancer region of *CYP1A* contains a bivalent domain, which is characterized by the presence of post-translational, covalent histone modifications that are associated with both gene activation and gene repression(61). This bivalent chromatin state is present in both embryos and adult fish and its response to PAH challenge is consistent with observed *CYP1A* gene expression. In addition, PAH-tolerant adult fish with a history of PAH exposure show persistent induction of *CYP1A*, in the absence of concurrent exposure. These findings suggest the presence of two distinct, but related epigenetic memories at *CYP1A*. Since *CYP1A*(*1*) induction is responsible for PAHs’ cancer-causing effects^3^, people that show blunted *CYP1A* induction in response to PAH are likely to be protected against cancer. In contrast, people that show higher baseline *CYP1A*(*1*) expression or enhanced *CYP1A*(*1*) induction secondary to sustained PAH exposure are likely to be at increased risk for cancer. Our results are a critical first step in the development of preventive or therapeutic strategies for suppressing persistent induction of *CYP1A1* and reducing cancer risk in human populations with sustained, high dose PAH exposures.

## Results

### The *CYP1A* promoter-enhancer contains an environmentally responsive bivalent domain

To explore the epigenetic memory at *CYP1A*, we performed experiments in wild-caught depurated adults and laboratory-bred F_1_ embryos from both PAH-tolerant and PAH-sensitive populations of *F. heteroclitus* (**Fig. 1A**). We did not observe AhR motif sequence differences between PAH-tolerant and PAH-sensitive adult mummichog (**Fig. 1B, Supporting text, Supp. Table 1, Supp. Fig. 1**), which ruled out the most likely genetic explanation for reduced *CYP1A* inducibility in PAH-tolerant fish. Therefore, we explored potential epigenetic explanations. An epigenetic trait is a mitotically (and possibly meiotically) heritable change in gene expression or gene responsiveness that is not explained by genetic alterations(62–65). Epigenetic traits involve inheritance of signaling molecules (e.g., coding or non-coding RNA, transcription factors)(62, 63) or of structural chromatin features(66–69) or both. Before evaluating the *CYP1A* memory as an epigenetic trait, we sought to confirm prior reports of gene expression patterns at *CYP1A*. Using RT-qPCR, we quantified *CYP1A* expression in 14-day-old embryos (*N*=3 pools of 10 embryos each) derived from depurated, wild-caught PAH-sensitive and PAH-tolerant adult mummichog (**Fig. 1A**). To quantify effect sizes in gene expression comparisons, we computed Hedge’s g, which expresses effect sizes in units of standard deviation (see Methods) and is best for comparisons between groups with large variation in biological outcomes(70), including gene expression patterns in these genetically diverse, wild fish (**Fig. 1C-E**). Using this approach, we observed ∼4.5-fold higher basal *CYP1A* expression in PAH-sensitive embryos, as compared to PAH-tolerant embryos (Hedge’s g: 1.68, 95% CI: −0.07, 3.32) (**Supp. Fig. 2, Supp. Tables 2-3**). (Hedge’s g equal to or greater than 0.8 standard deviations represents a large effect, 0.4-0.6 is a moderate effect, and 0.2 is a small effect(70) Hedge’s g point estimates are generally interpreted on this scale alone, but we add 95% confidence intervals here for completeness.) To test whether embryos from the two populations differed in their response to PAH challenge, we exposed additional embryo pools to 5% Elizabeth River sediment extract, which contains the PAH mixture present in the original contaminated site(56) (**Fig. 1A**). We confirmed prior reports of strong *CYP1A* induction in PAH-sensitive fish (>400-fold; Hedge’s g: −4.57, 95% CI: −7.45, −1.27) (**Fig. 1C, Supp. Fig. 2, Supp. Tables 2-3**) and substantially muted *CYP1A* induction in PAH-tolerant fish (∼100-fold; Hedge’s g: −8.83, 95% CI: −14.85, −2.90) (**Fig. 1C, Supp. Tables 2-3**). The Hedge’s g metric was larger for the PAH-tolerant embryos, as compared to the PAH-sensitive, despite a much larger mean fold-change in expression in PAH-sensitive embryos, because the standard deviation in *CYP1A* induction was much larger in the sensitive group than in the tolerant (**Fig. 1C, Supp. Fig. 2, Supp. Tables 2-3**). This result indicates that either epigenetic memory formation at *CYP1A* or genetic adaptation to PAH or a combination of both effects leads to reduced inter-individual variation in PAH response, which suggests a loss of plasticity. The Hedge’s g metrics for these comparisons were both negative, since we computed differences in raw cycles-to-threshold (C(t)) values (see Methods), and C(t) is inversely correlated with gene expression. To formally quantify the difference in PAH response between populations, we computed the difference-in-difference (D-i-D), or difference in Hedge’s g values; the D-i-D for [PAH-tolerant (treatment-control) – PAH-sensitive (treatment-control)] was −4.26 (95% CI: −7.03, −1.45) (**Fig. 1C, Supp. Tables 2-3**), indicating an extremely large difference in *CYP1A* induction by PAH between the groups.

**Figure 1.**
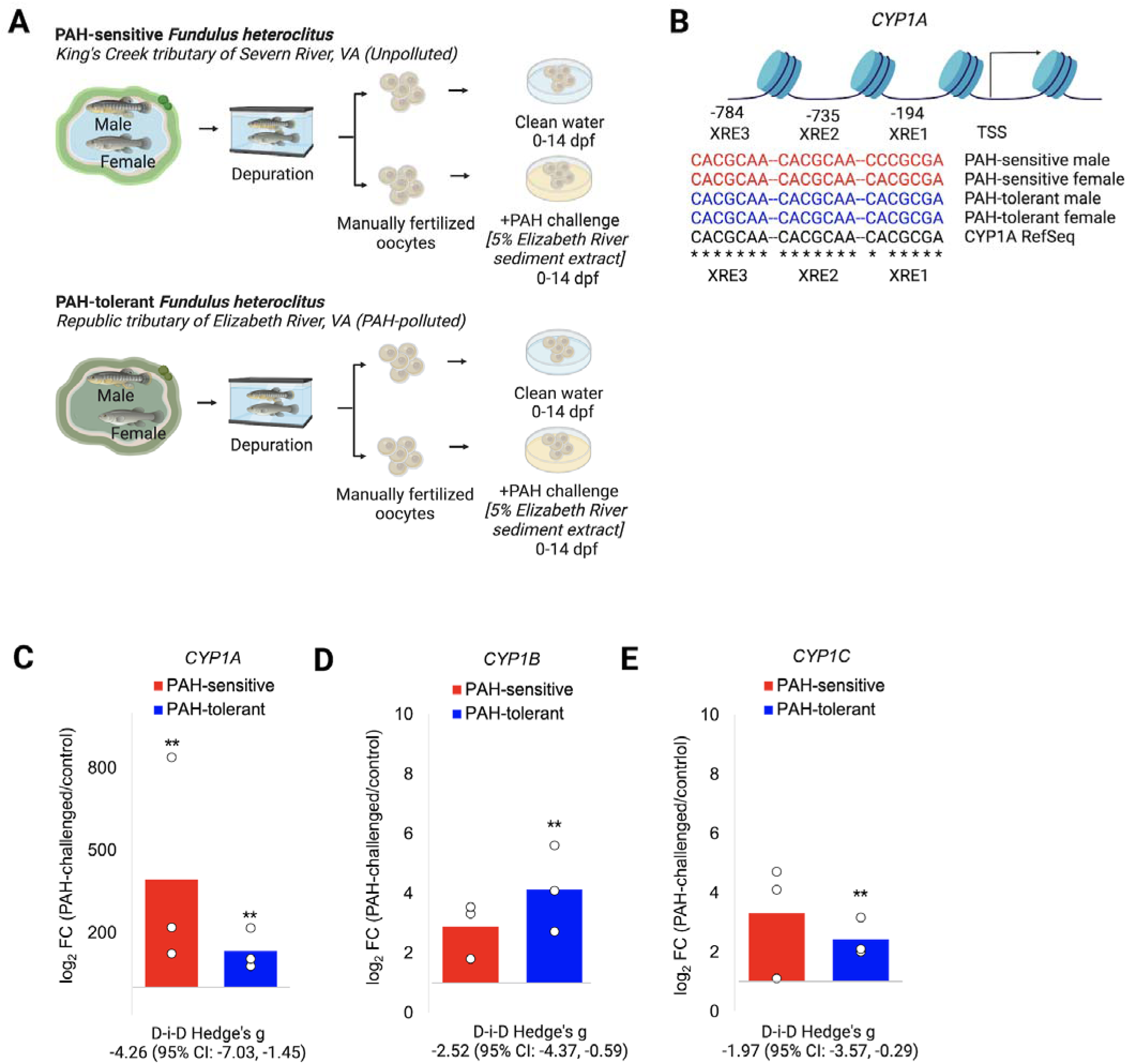
The epigenetic memory at *CYP1A* in PAH-tolerant *Fundulus heteroclitus*. **(A)** Experimental design schematic for depuration of wild-caught fish, manual fertilization of embryos, and exposure to PAH. **(B)** Xenobiotic response elements (XREs) in the *CYP1A* proximal promoter-enhancer are conserved across four fish (one male and one female each from PAH-tolerant and PAH-sensitive populations). **(C-E)** Relative fold-change values for embryonic gene expression for three AhR target genes (*CYP1A*, *CYP1B*, and *CYP1C*). Figures show C(t) values from PAH-challenged embryo samples normalized to the average control C(t). \*\**P*-value <0.01 from independent samples t-tests comparing C(t) values from PAH-challenged embryos to control embryos within each group. Hedge’s g values represent effect sizes for difference-in-difference (D-i-D) tests [PAH-tolerant (treatment-control)] – [PAH-sensitive (treatment-control)]. Hedge’s g = 0.2 is a small effect, Hedge’s g =0.4-0.6 is a moderate effect, Hedge’s g = or > 0.8 is a large effect. 95% confidence intervals that do not cross zero reflect p<0.05. Dpf – days post-fertilization, PAH – polycyclic aromatic hydrocarbons, D-i-D – Difference-in-difference, XRE – xenobiotic response element, TSS – transcription start site.

Next, we tested for the presence of similar responses at the related metabolic AhR gene targets, *CYP1B* and *CYP1C*. Prior studies showed similarly heritable but reversible resistance of these genes to induction by PAH in tolerant fish(55). Both *CYP1B* (PAH-sensitive: Hedge’s g: −1.62, 95% CI: −3.24, 0.10; PAH-tolerant: Hedge’s g: −4.15, 95% CI: −7.16, −1.10) and *CYP1C* (PAH-sensitive: Hedge’s g: −1.41, 95% CI: −2.95, 0.23; PAH-tolerant: Hedge’s g: −3.39, 95% CI: −5.95, −0.77) were induced by PAH in embryos, albeit to a lesser degree than *CYP1A* (**Fig. 1D-E, Supp Tables 2-3**). Like *CYP1A*, both genes showed larger Hedge’s g effect sizes in PAH-tolerant embryos, which suggests similarly reduced inter-individual differences in PAH response as a consequence of adaptation (*CYP1B* D-i-D −2.52, 95% CI: −4.37, −0.59; *CYP1C* D-i-D −1.97, 95% CI: −3.57, −0.29) **(Supp Tables 2-3**). *CYP1C*, but not *CYP1B*, displayed greater average fold-change response to PAH in sensitive embryos than in tolerant ones (**Fig. 1D-E, Supp. Tables 2-3)**.

We hypothesized that these gene expression patterns resulted from epigenetic inheritance of exposure-induced changes in structural chromatin features. Structural chromatin features, including methylation of cytosine-guanine dinucleotides in DNA and post-translational modifications to histone proteins (65), regulate gene expression(65) and are both environmentally responsive(71) and inherited through cell division(67–69). One earlier study reported a complete absence of DNA methylation in the *CYP1A* promoter-enhancer in both PAH-tolerant and -sensitive fish, with or without PAH exposure(72). Therefore, using chromatin immunoprecipitation (ChIP), we explored the potential roles of three histone post-translational modifications (PTMs): trimethylation of lysine 4 on histone H3 (H3K4me3), which is associated with active promoters(73); H3K4me1, which marks active enhancers(74); and H3K27me3, which is associated with gene repression at both promoters and enhancers(75–77). We anticipated that the *CYP1A* promoter-enhancer in wild-type, PAH-sensitive fish would show low levels of activating histone PTMs at baseline that would increase in response to PAH challenge, and no repressive histone PTMs. In PAH-tolerant fish, we hypothesized that levels of activating histone PTMs would increase with PAH challenge, but to a lesser degree than in sensitive fish. We also anticipated that repressive histone PTMs would be newly present at the promoter to counteract strong gene induction. However, we observed a bivalent chromatin state, characterized by activating and repressive histone modifications(61, 78), encompassing all XREs and the TSS in both PAH-sensitive and PAH-tolerant embryos (**Fig. 2A-C, Supp. Fig. 3, Supp. Tables 4-8**). However, tolerant embryos showed lower levels of H3K4me3 and higher levels of both H3K4me1 and H3K27me3 at baseline (**Supp. Figs. 3-5, Supp. Tables 4-8**). These differences in baseline histone PTMs may be due to non-genetic inheritance of parental PAH response or to unidentified genetic changes disrupting sequence specificity for histone-containing nucleosomes(79) or to a combination of both effects. In addition, embryos from each population displayed markedly different responses to PAH challenge (**Fig. 2A-C, Supp. Fig. 3, Supp. Tables 4-8**). Consistent with strong gene induction, PAH-sensitive embryos showed modest increases in average enrichment of both activating modifications at the TSS, and in H3K4me1 at XREs, combined with a decrease in repressive H3K27me3 across the promoter-enhancer (**Fig. 2A-C, Supp. Fig. 3, Supp. Tables 4-8**). However, despite a marked increase in *CYP1A* expression in these embryos, most of these differences in mean histone PTM levels yielded small to moderate effect sizes in mixed models comparing exposed to control pools. This outcome likely is attributable to high inter-individual variation, including in H3K4me3 at the TSS (Hedge’s g: 0.08) and at all four loci for both H3K4me1 (TSS Hedge’s g: 0.34; XRE1 Hedge’s g: 0.44; XRE2 Hedge’s g: 0.39; XRE3 Hedge’s g: 0.06) and H3K27me3 (TSS Hedge’s g: −0.22; XRE1 Hedge’s g: −0.22; XRE2 Hedge’s g: 0.04; XRE3 Hedge’s g: −0.19) (**Fig. 2A-C, Supp. Fig. 3, Supp. Tables 4-8**). In contrast, PAH-tolerant embryos showed a muted increase in average enrichment of H3K4me3 at the TSS and dramatic decreases in both H3K4me1 and H3K27me3 (**Fig. 2A-C, Supp. Fig. 3, Supp. Tables 4-8**). Following the same pattern as *CYP* expression data, effect sizes expressed in standard deviations were larger in tolerant embryos than in sensitive embryos for both H3K4me1 (TSS Hedge’s g: −0.50; XRE1 Hedge’s g: −0.63; XRE2 Hedge’s g: −0.25; XRE3 Hedge’s g: −0.56) and H3K27me3 (TSS Hedge’s g: −1.17; XRE1 Hedge’s g: −1.20; XRE2 Hedge’s g: −1.36; XRE3 Hedge’s g: −1.07), and accordingly smaller for H3K4me3 at the TSS (Hedge’s g: 0.08). We confirmed the differences in effects between populations by computing D-i-D metrics for [PAH-tolerant (treatment-control) – PAH-sensitive (treatment-control)] (**Fig. 2A-C, Supp. Tables 4-8**). The pattern of activating modifications that we observed in tolerant embryos was consistent with decreased promoter and enhancer activity, which matched our observed blunted *CYP1A* induction. Decreased regulatory element activity may be due to reduced AHR binding or altered enhancer response to AHR binding. However, we did not have an immediate explanation for the significant decrease in H3K27me3 in tolerant embryos challenged with PAH. We reasoned that, because these modifications work together to produce a chromatin state at bivalent loci, the relative proportion of repressive to activating modifications would be more informative than enrichment of individual modifications. Therefore, we calculated the ratio of repressive to activating modifications (H3K27me3: H3K4me1 and H3K27me3: H3K4me3) (**Fig. 2D-E, Supp. Fig. 4, Supp. Tables 9-10**). Neither ratio differed by population at baseline (**Supp. Fig. 4, Supp. Tables 9-10**). In response to PAH challenge, the H3K27me3: H3K4me1 ratio showed a substantial decrease in sensitive embryos (i.e., large increases in the relative enrichment of activating modifications, as compared to repressive ones), as compared to a slight decrease in tolerant embryos (**Fig. 2D, Supp. Fig. 4, Supp. Table 9**). These findings agree with the blunted gene induction seen in these embryos. The H3K27me3: H3K4me3 ratio showed similar decreases in response to PAH; sensitive embryos showed greater decreases in this ratio at the TSS and its nearest XRE, XRE1, as compared to tolerant ones (**Fig. 2E-F, Supp. Fig. 4, Supp. Table 10**). Together, these data indicate a generationally heritable epigenetic memory of ancestral PAH exposure in a PAH-tolerant population. This memory manifests in blunted responses to PAH challenge at *CYP1A*, both in regulation of a promoter bivalent domain and in transcriptional output, in exposure-naïve embryos.

**Figure 2.**
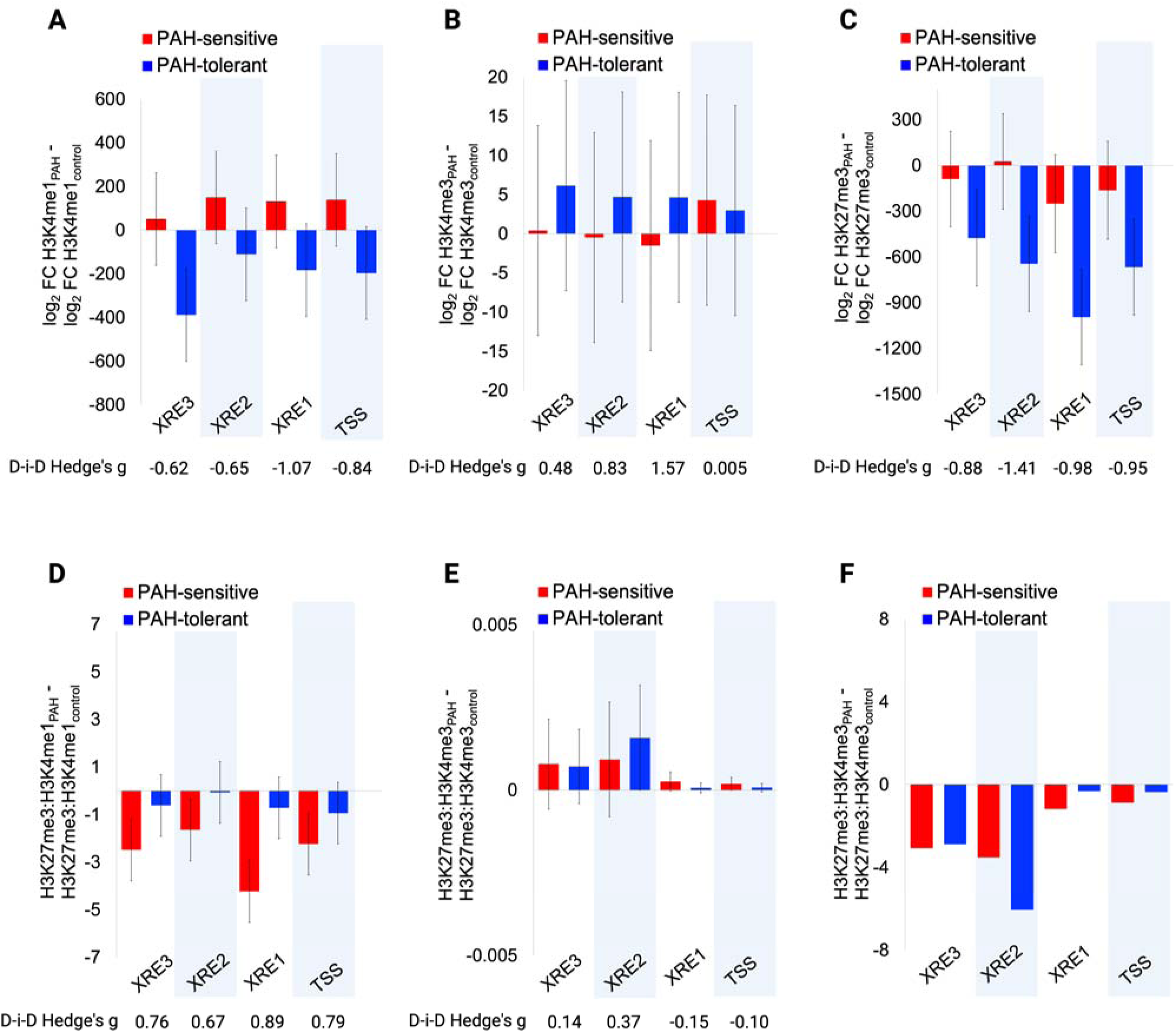
The *F. heteroclitus CYP1A* promoter-enhancer contains an environmentally responsive bivalent domain. **(A-C)** Mean and SEM for differences in histone PTM (H3K4me1, H3K4me3, H3K27me3) enrichment at the *CYP1A* proximal promoter-enhancer in embryos challenged with PAH (relative to control) from PAH-tolerant and PAH-sensitive populations. **(D-F)** Mean and SEM for differences in ratios of repressive/activating histone PTMs (**(D)** H3K27me3/H3K4me1; (E) H3K27me3/H3K4me3) in embryos challenged with PAH (relative to control) from PAH-tolerant and PAH-sensitive populations. The ratio differences shown in (**E**) are inverse transformed, due to skewed data distribution. For ease of interpretation, differences in back-transformed means are shown in **(F)**. Hedge’s g values represent effect sizes for difference-in-difference (D-i-D) tests [PAH-tolerant (treatment-control)] – [PAH-sensitive (treatment-control)]. Hedge’s g = 0.2 is a small effect, Hedge’s g =0.4-0.6 is a moderate effect, Hedge’s g = or > 0.8 is a large effect. 95% confidence intervals shown in Supplemental Tables 5-8. XRE – xenobiotic response element, TSS – transcription start site.

### Continuous PAH exposure triggers a secondary epigenetic memory at *CYP1A* in adult fish

Most prior evidence of the *CYP1A* epigenetic memory was generated through studies of developmental toxicity of PAH and therefore focused on testing effects in exposure-naïve embryos. However, if memory formation is distinct from PAH developmental toxicity, then we would expect to see memory effects in adult animals, too.

To test this hypothesis, we measured *CYP1A* mRNA levels in liver tissue from depurated, wild-caught adult fish from both sensitive and tolerant populations (parents of the embryos used in the prior set of experiments) (*N*=5 males and *N*=5 females per population). PAH-sensitive adults in this experiment were PAH-naïve; PAH-tolerant adults had no ongoing or residual PAH exposure but did have a long history of sustained, extreme exposure in the wild. We focused on liver tissue, since fish do not have human-equivalent lungs; fish are primarily exposed to PAHs via ingestion of contaminated water(56) and humans are primarily exposed via inhalation of contaminated air(1). In addition, th*e CYP1A* memory was previously characterized in adult mummichog liver tissue(52, 53), since liver is the organ with the highest concentration of CYP1A protein(26). We observed higher *CYP1A* expression in PAH-tolerant adults, as compared to PAH-sensitive adults, even in the absence of PAH challenge (Hedge’s g: −1.43, 95% CI: −2.37, −0.45) (**Fig. 3A, Supp. Fig. 5, Supp. Tables 11-12**). Notably, this effect was strongest in male fish (Hedge’s g: −2.38, 95% CI: −3.94, −0.74), as compared to female fish (Hedge’s g: −0.56, 95% CI: −1.69, 0.6) (D-i-D for [(PAH-tolerant males - PAH-sensitive males) – (PAH-tolerant females - PAH-sensitive females)] Hedge’s g: −1.82, 95% CI: −3.0, −0.59) (**Fig. 3A, Supp. Fig. 5, Supp. Tables 11-12**). This result indicates a larger difference between males from the two populations than between females.

**Figure 3.**
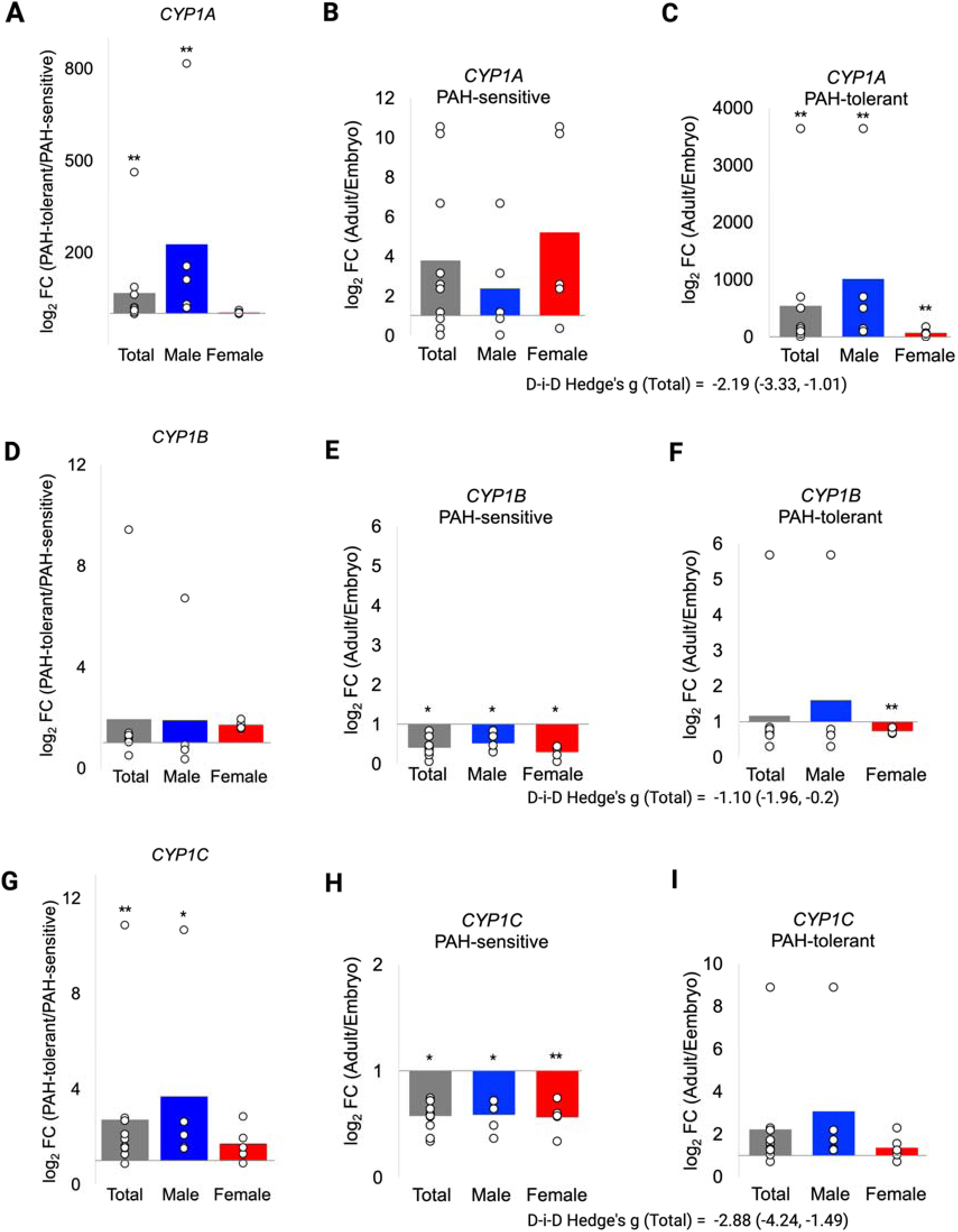
AhR target genes remain induced in adult PAH-tolerant *F. heteroclitus* in the absence of PAH. **(A, D, G)** Relative fold-change in gene expression of the AhR target genes *CYP1A* (a), *CYP1B* (d), and *CYP1C* (g) in adult fish from a PAH-tolerant vs. PAH-sensitive population. **(B-C, E-F, H-I)** Relative fold-change in *CYP1A* (b-c), *CYP1B* (e-f), and *CYP1C* (h-i) expression in adult fish relative to embryos from the same population (PAH-sensitive or PAH-tolerant). All figures show C(t) values from adult liver tissue normalized to the average C(t) for each comparison group (PAH-sensitive adults in (A, D, G) and embryos in (B-C, E-F, H-I)), separated by sex. \*\**P*-value <0.01 and * P-value <0.05 for independent samples t-tests of C(t) values. Hedge’s g values represent effect sizes for difference-in-difference (D-i-D) tests [PAH-tolerant (adult-embryo)] – [PAH-sensitive (adult-embryo)]. the label “Total” indicates combined male and female data. Hedge’s g = 0.2 is a small effect, Hedge’s g =0.4-0.6 is a moderate effect, Hedge’s g = or > 0.8 is a large effect. 95% confidence intervals that do not cross zero reflect p<0.05.

Since male fish from both populations are known to exhibit higher magnitude induction of *CYP1A* in response to PAH than female fish(52, 53, 80), and, in our experiment, tolerant female fish did not show residual activity after exposure cessation, this result strongly suggests incomplete shutoff of the gene after exposure cessation in male fish. It is possible that this effect was partially due to higher gene expression in males in both populations. To rule this out, we tested whether *CYP1A* expression was sex-specific within populations, and if so, whether this sex difference was the same in both populations. We observed a moderate increase in baseline *CYP1A* expression in sensitive female fish over male fish (Hedge’s g: 0.60, 95% CI: −0.57, 1.74) (**Supp. Fig. 5, Supp. Tables 11-12**), but a very substantial increase in residual *CYP1A* expression in tolerant male fish vs. females (Hedge’s g: −1.42, 95% CI: −2.70, −0.08) (**Supp. Fig. 5, Supp. Tables 11-12**). We also tested whether our observation of differences in adult expression was partially due to expected changes with normal development. To evaluate this, we compared *CYP* gene expression in each group of adult fish to gene expression in the group of population-matched embryos. *CYP1A* expression increases up to 10-fold in PAH-sensitive adult fish, as compared to embryos (**Fig. 3B, Supp. Figs. 1 and 5, Supp. Table 13**). In contrast, *CYP1A* expression increased ∼1000-fold in male and ∼70-fold in female PAH-tolerant adults, as compared to embryos (**Fig. 3C, Supp. Figs. 1 and 5, Supp. Table 13**). Taken together, our findings support the interpretation that *CYP1A* expression in PAH-tolerant male adults represents incomplete recovery from prior PAH exposure. These results suggest the formation of a secondary epigenetic memory within individual, PAH-tolerant fish that forms with sustained PAH challenge.

To further characterize this secondary epigenetic memory in adult PAH-tolerant fish, we tested two additional hypotheses. Some prior data on epigenetic memory suggests that the strength of initial gene induction predicts epigenetic memory formation(81). Therefore, we asked whether sustained gene expression following exposure cessation was specific to *CYP1A*, a gene with a high induction response (>100-fold). If so, we would not see similarly sustained responses at AHR target genes *CYP1B* and *CYP1C*, both of which are induced <10-fold by PAH in exposure-naïve embryos from both populations (**Fig. 1D-E**). Contrary to this hypothesis, we observed similarly sustained expression for both genes in tolerant adults (**Figs. 3D-I, Supp. Fig. 5, Supp. Tables 11-12**).

Next, we asked whether this secondary epigenetic memory was associated with altered regulation of the promoter-enhancer bivalent domain. Bivalent domains were initially described as key regulators of developmental gene expression(61, 78, 82–84). At most loci, bivalency resolves during lineage determination and cellular differentiation; loci that initiate transcription lose H3K27me3 and retain H3K4me3(85–92), and loci that are silenced retain H3K27me3 and lose H3K4me3(78). However, not all bivalent regions resolve in differentiated tissues(61, 93–95). To test whether the bivalent domain persists at *CYP1A* in adult mummichog, we measured histone PTMs in liver tissue from the same wild-caught adult male fish that showed residual CYP gene expression (**Fig. 3A-I**). We confirmed the persistence of the bivalent domain encompassing all three XREs and TSS in adult fish from both populations (**Fig. 4D-F, Supp. Table 1**.) As compared to sensitive males, tolerant male fish showed approximately equal levels of activating H3K4me1 (**Supp. Table 15**), moderately lower levels of activating H3K4me3 (**Supp. Table 16**), and moderately higher levels of repressive H3K27me3 (**Supp. Table 17**), none of which correlated well with the increased *CYP1A* expression in these males (**Fig. 3A-C**). In addition, the H3K27me3: H3K4me1 ratio was moderately elevated in the two XREs nearest to the TSS (**Supp. Table 18**), and the H3K27me3: H3K4me3 ratio was elevated across the promoter-enhancer region, particularly at the TSS (**Supp. Table 19**). These data indicate a relative increase in repressive signal at this bivalent domain in tolerant animals. To confirm that these results were not driven by differences in baseline histone PTMs or in changes in histone modifications with development, we compared histone PTM levels within each group of adult fish to levels in population-matched, exposure-naïve embryos and then compared developmental trajectories between populations. In agreement with the results from our initial analyses, tolerant male adults showed smaller decreases in activating PTMs and substantial increases of the repressive PTM when compared to their matched embryos, as compared to sensitive male fish (**Fig. 4, Supp. Figs. 6-8, Supp. Tables 20-23**). Similarly, we observed increases (both in mean enrichment and in Hedge’s g values that account for population variation) in both ratios in PAH-tolerant adults, as compared to PAH-sensitive ones (**Fig. 5, Supp. Figs. 9-10, Supp. Tables 24-25**). We speculate that this result indicates a progressive loss of regulatory control of *CYP1A* transcription in PAH-tolerant adults, perhaps related to the differential PAH response that we observed in matched embryos. The relative increases in the mildly repressive H3K27me3 may represent a compensatory response in tolerant fish, in an attempt to shut off the persistently induced *CYP1A* gene. Together, these data indicate an additional, mitotically heritable epigenetic memory of sustained PAH exposure that manifests in sustained increases in repressive: activating histone PTM ratios in the *CYP1A* bivalent domain and a sustained increase in expression of the *CYP1A* gene after cessation of PAH stimulus.

**Figure 4.**
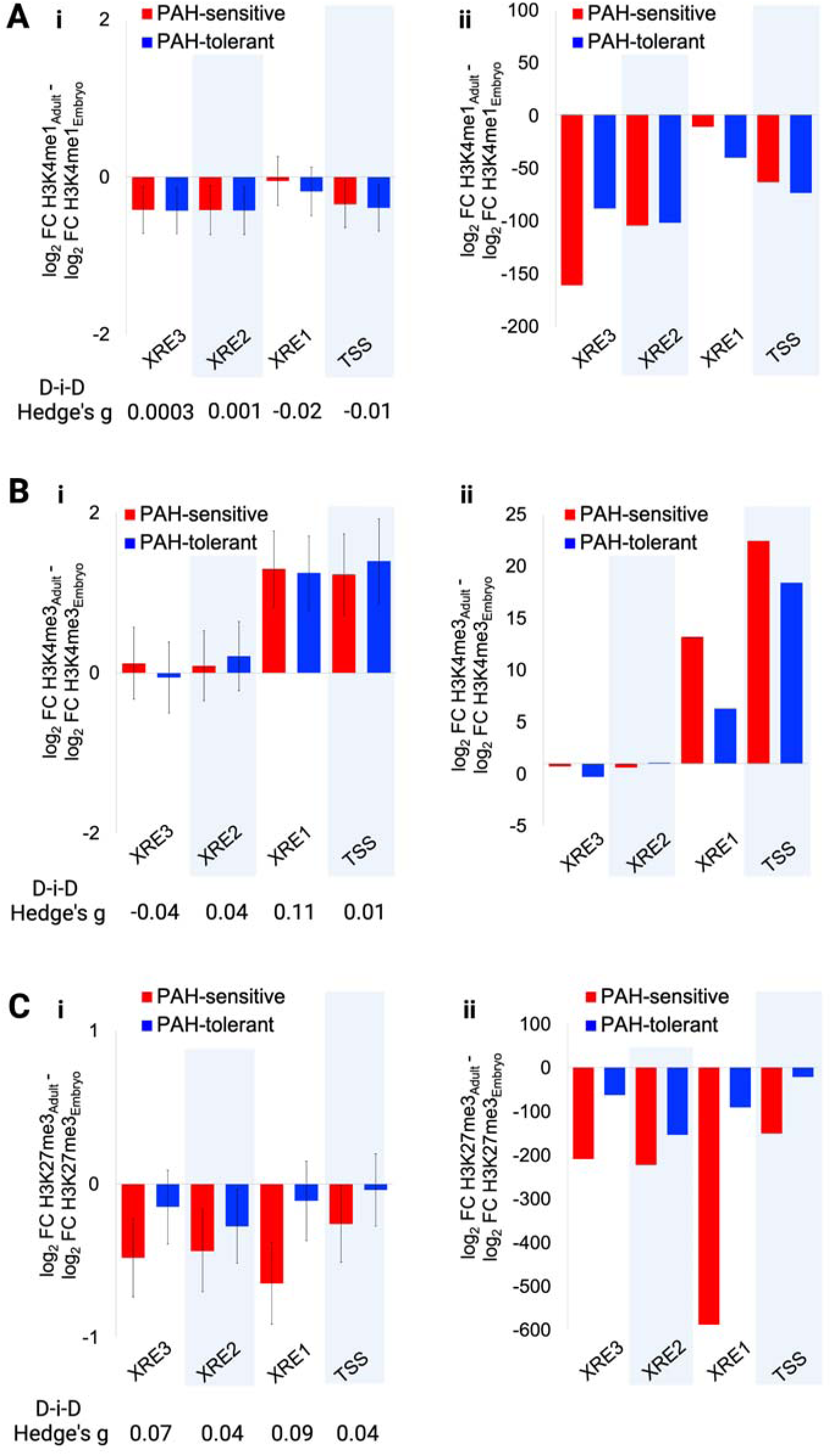
Histone modifications at the bivalent domain in *CYP1A* reflect active gene expression in PAH-tolerant fish with past but no current PAH exposure. **(A_i_, B_i_, C_i_)** Mean and SEM for differences in inverse-transformed histone PTM H3K4me1 (A_i_), H3K4me3 (B_i_), and H3K27me3 (C_i_) at the *CYP1A* proximal promoter-enhancer in adult fish compared to matched embryos from PAH-tolerant and PAH-sensitive populations. **(A_ii_, B_ii_, C_ii_)** Differences in back-transformed means for H3K4me1 (Aii), H3K4me3 (B_ii_), and H3K27me3 (C_ii_), for ease of interpretation. Hedge’s g values represent effect sizes for difference-in-difference (D-i-D) tests [PAH-tolerant (adult-embryo)] – [PAH-sensitive (adult-embryo)]. Hedge’s g = 0.2 is a small effect, Hedge’s g =0.4-0.6 is a moderate effect, Hedge’s g = or > 0.8 is a large effect. 95% confidence intervals shown in Supplemental Tables 9-10. XRE – xenobiotic response element, TSS – transcription start site.

**Figure 5.**
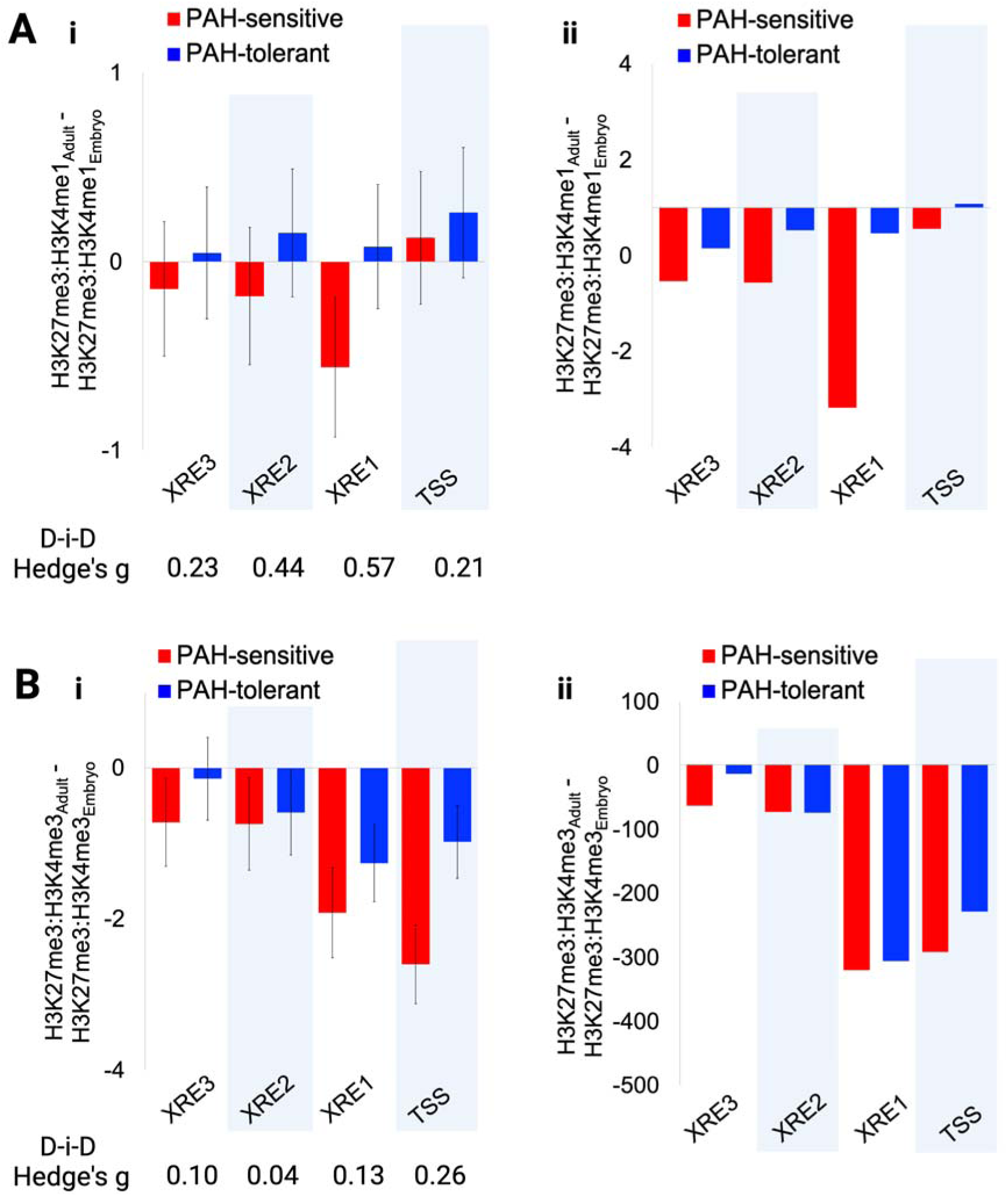
Ratios of repressive: activating histone modifications at *CYP1A* reflect active gene expression in PAH-tolerant fish with past but no current PAH exposure. **(A_i_, B_i_)** Mean and SEM for differences in inverse transformed ratios of repressive/activating histone PTMs ((A) H3K27me3/H3K4me1; (B) H3K27me3/H3K4me3) in adult vs. embryos from both populations. **(A_ii_, B_ii_)** Differences in back-transformed means in ratios. Hedge’s g values represent effect sizes for difference-in-difference (D-i-D) tests [PAH-tolerant (adult-embryo)] – [PAH-sensitive (adult-embryo)]. Hedge’s g = 0.2 is a small effect, Hedge’s g =0.4-0.6 is a moderate effect, Hedge’s g = or > 0.8 is a large effect. 95% confidence intervals shown in Supplemental Tables 24-25. XRE – xenobiotic response element, TSS – transcription start site.

### The human *CYP1A1* promoter contains a bivalent domain and a CpG island

To evaluate the generalizability of this phenotype to human populations, we asked whether a similar bivalent domain is present at the human orthologs, *CYP1A1/2* and *CYP1B1*. Humans have no known ortholog of *CYP1C*. Using publicly available Epigenome Roadmap reference datasets, we identified bivalent domains in the promoters of the *CYP1A1*, but not *CYP1A2,* gene in human embryonic stem cells (H1 ESCs) and terminally differentiated, karyotypically normal lung fibroblasts (IMR90) (**Supp. Figs. 11-12**). These findings support a role for this bivalent domain in regulation of this gene in both embryonic and adult tissues. Because bivalent chromatin preferentially forms in DNA sequence that contains a high density of cytosine-guanine dinucleotides, or CpG islands,(61) we tested for the presence of a CpG island at the mummichog *CYP1A* and the human *CYP1A1*. We observed two CpG islands upstream of the mummichog *CYP1A*, one of which overlaps the *CYP1A* promoter-enhancer region (**Supp. Fig. 13**). Similarly, we observed a CpG island upstream of the human *CYP1A1* isoform, which is consistent with formation of bivalent domains at the human *CYP1A*(*1*) ortholog (**Supp. Fig. 11**). Therefore, human *CYP1A1* contains the known structural chromatin features that characterize the *CYP1A* gene in mummichog, which supports the generalizability of the mummichog *CYP1A* memory phenotype to humans.

## Discussion

Here, we characterize an epigenetic memory at the *CYP1A* gene that arose naturally in a wild Atlantic mummichog population as it adapted to chronic, extreme PAH pollution. In PAH-tolerant fish, *CYP1A* induction is blunted. However, tolerant fish reared in clean water in the laboratory for one generation partially recover *CYP1A* inducibility. To explore the underlying mechanism, we caught wild mummichog from PAH-tolerant and PAH-sensitive populations, manually fertilized eggs, and exposed half of the embryos in each group to 5% Elizabeth River sediment extract for 14 days post-fertilization. We identified a bivalent domain, characterized by both activating and repressive histone post-translational modifications, in the *CYP1A* proximal promoter-enhancer in both sensitive and tolerant embryos and adults. In response to PAH, sensitive embryos showed more marked decreases in ratios of repressive: activating histone PTMs (consistent with gene activation), as compared to tolerant embryos, indicating active regulation by this domain. In addition to blunted induction of *CYP1A*, we observed that unexposed, PAH-tolerant adult males with a history of extreme PAH exposure showed persistent upregulation of *CYP1A*, which is consistent with a defect in gene recovery to baseline. Notably, PAH-sensitive fish showed greater variation than did PAH-tolerant fish in *CYP1A* expression and histone PTM enrichment both at baseline and in response to PAH. This result mirrors the loss of genetic diversity previously reported in PAH-tolerant fish(57) and suggests that genetic differences between tolerant and sensitive populations play a partial role in the observed epigenetic memory at *CYP1A*.

Overall, our data are consistent with two related epigenetic memories at *CYP1A*. The first memory is characterized by blunted PAH-triggered induction of *CYP1A* in PAH-tolerant embryos that correlates well with changes in histone PTMs in the promoter bivalent domain. This memory may be unique to *CYP1A*; alternatively, it may reflect protective epigenetic downregulation of AhR signaling that is reflected in localized chromatin responses at target genes, including *CYP1A, CYP1B, and CYP1C*. In the second scenario, epigenetic downregulation likely develops to protect against teratogenic effects, rather than cancer; regardless of its etiology, this memory protects against cancer. PAH-tolerant *Fundulus* living in PAH-contaminated areas of the Elizabeth River show higher rates of cancer in liver (the primary PAH target tissue in fish), as compared to PAH-sensitive fish living in a clean environment. However, clean water-reared F_1_ larvae of wild-caught sensitive fish exposed to a standardized dose of PAH developed more juvenile liver tumors, as compared to similarly treated tolerant F^1^ larvae(58). Second, PAH-tolerant adult fish with a history of continuous PAH exposure show persistent induction of *CYP1A* after exposure cessation, which is consistent with defective *CYP1A* gene shutoff and recovery to baseline. The second memory is characterized by persistent induction of *CYP1A* after exposure cessation in PAH-tolerant adult fish with a history of chronic, continuous PAH exposure, which is consistent with defective *CYP1A* gene shutoff and recovery to baseline. Since *CYP1A* expression is positively correlated with cancer risk in both mummichog and humans, these results indicate that the PAH-tolerant fish have epigenetic protection against PAH-induced cancer in early life that degrades over time in response to continuous gene activation.

These findings on naturally occurring epigenetic memories at *Fundulus CYP1A* represent an important advance over existing data on experimentally induced epigenetic memories at *CYP1A1/Cyp1a1* in mammals in response to acute exposures(96–99). Four prior studies report mammalian epigenetic memories at this gene; three of these four studies show than an initial stimulus triggered sustained induction of *Cyp1a1* in exposure naïve rodents(96, 97) or *CYP1A1* in exposure naïve human cells(98) and the fourth showed both sustained induction and superinduction on secondary challenge(99). Those studies model initial responses in human populations without significant historical PAH exposures. However, few human populations today are naïve to PAH exposure, and most are likely to experience episodic, possibly chronic, exposure as wildfires increase in frequency. Therefore, PAH-tolerant *Fundulus* represent a more informative model of human population responses. Our data suggest that chronically exposed human populations may develop protective epigenetic memories that increase adult cancer risk.

Increased understanding of the epigenetic memory at mummichog *CYP1A* provides insight into approaches for protecting human health in the context of increased population exposure to PAHs. Specifically, we can leverage fundamental understanding of the *CYP1A* memory to develop preventive or therapeutic approaches and to protect individuals that naturally form this memory from deleterious side effects. CYP1A1 is druggable with small molecule inhibitors(100–103), including dietary polyphenols(100–103), or with synthetic substrates that are converted to protective molecules (e.g., compounds that are cytotoxic to cancer cells) at a higher rate when *CYP1A1* is induced(104–112). Preventive or therapeutic treatment with these compounds may protect individuals that are either at high cancer risk from wildfire smoke exposure or at high risk of exacerbation of pre-existing cancer.

## Conclusion

Our findings represent an important advance in understanding the formation and persistence of an epigenetic memory at *CYP1A* in response to chronic PAH exposure. Specifically, we showed altered regulation of the bivalent domain at *CYP1A* in embryos with a cancer-protective epigenetic memory. In addition, a secondary memory in adult fish leads to potentially harmful, sustained activation of *CYP1A* after sustained exposure to PAH. Since *CYP1A* induction is a critical step in PAH-related carcinogenesis, if similar memories form in human populations, they are likely to substantially impact cancer risk. Therefore, these results raise a critical new consideration in predicting the human health risks of increasing air pollution and wildfire events due to climate change.

## Methods

### Fish collection

We caught wild mummichog from a PAH-tolerant population from the site of former creosote wood treatment facility (Republic Creosoting) in the southern of the Elizabeth River in Virginia (36° 79’ 31.0’’ N, 76° 29’ 41.3’’ W) and from a PAH-sensitive reference population from King’s Creek, a relatively uncontaminated tributary of the Severn River in Virginia (37° 30’ 47.6” N, 76° 41’ 63.9” W). We depurated fish for at least 4 weeks prior to breeding in flow-through systems comprising a series of 30-40L tanks containing 20% artificial sea water (ASW, Instant Ocean, Foster & Smith, Rhinelander, WI, USA). We maintained adult fish at 23-25°C on a 14:10 light: dark cycle and ad libitum pelleted feed (Aquamax Fingerling Starter 300, PMI Nutrition International LLC, Brentwood, MO, USA). We obtained eggs from each population by manual spawning of females and fertilized eggs *in vitro* with expressed sperm from males in a beaker containing ASW. Embryos were held for one hour after spawning to allow for fertilization, then washed briefly with 0.3% hydrogen peroxide solution. For exposure experiments, we used a previously collected, processed, and characterized(113) sediment extract (Elizabeth River sediment extract, ERSE) from the Atlantic Woods Industries Superfund site, a former creosote wood treatment facility, in the Elizabeth River (VA, USA). This extract is a real-world mixture of water and suspended solids with a total PAH content of 5,073 ng/mL PAHs, summed from analyses of 36 different PAHs. We exposed half of the embryos in each group to 5% ERSE (diluted in 20% ASW) and the remaining half to clean water only. We chose this ERSE concentration based on previous studies showing CYP induction in both sensitive and tolerant mummichog with no lethality in sensitive fish. We dosed embryos in 20 mL glass scintillation vials (VWR, Westchester, PA, USA) at 27°C beginning at 24 hours post-fertilization (hpf). After 14 days, we flash froze embryos in liquid nitrogen and stored at −80°C. We dissected liver tissue from depurated, wild-caught adult fish, flash froze tissue in liquid nitrogen and stored at −80°C. All care, reproductive techniques and rearing techniques were non-invasive and approved by the Duke University Institutional Animal Care and Use Committee (A139-16-06).

### Genomic DNA isolation

We extracted genomic DNA from adult liver tissue with DNeasy Blood & Tissue Kits (cat. No. 69504, Qiagen, Hilden, Germany). Briefly, we cut ∼20 mg of liver tissue over dry ice. We added 180 µL of buffer ATL, minced the tissue over wet ice and transferred the sample to a 1.5mL safe lock tube. After adding 20 µL of Proteinase K, we incubated samples at 56°C overnight. We followed the kit protocol for on-column purification, and eluted the final genomic DNA in 200 µL milliq-H_2_O and incubated at room temperature before centrifuging at 8,000 rpm for 1 minute and storing at −20°C.

### *CYP1A* resequencing and contig assembly

We re-sequenced the *CYP1A* gene and 7Kb upstream of the *CYP1A* TSS via primer walking and Sanger sequencing in *N*=4 wild-caught adult mummichog, one male and one female each from PAH-tolerant and -sensitive populations. The completed sequence spans chromosomal coordinates chr4:1,265,806-1,276,399 (NCBI RefSeq assembly Fundulus_heteroclitus-3.0.2 (2015), accession NW_012234324.1). We performed PCR in 12.5-25 µl reactions using Platinum SuperFi II Green PCR Master Mix (#12359010; Thermo Fisher Scientific, Waltham, MD), 0.5 µl of sample DNA, and forward and reverse primers to a final concentration of 500nM on a Biometra T Advanced Thermocycler (Analytik Jena, Jena, Germany) under the following conditions: 98°C for 30 seconds, followed by 35 cycles of 98°C for 5 seconds, primer annealing temperature for 20 seconds, extension at 72°C for 30 seconds, with a final extension at 72°C for 5 minutes. We diluted the products 1:4 with MilliQ-H_2_O and submitted them for Sanger sequencing through the OHSU Vollum DNA Sequencing Core using an ABI (Applied Biosystems) 3730*xl* 96-capillary DNA Analyzer. We manually edited the resulting ABI files before assembling overlapping sequences into contigs utilizing the PRABI Cap3 Sequence Assembly Program (https://doua.prabi.fr/software/cap3) and aligned contigs and the published reference sequence (GCF_000826765.1) using the CLUSTALW Multiple Sequence Alignment tool (https://www.genome.jp/tools-bin/clustalw). We analyzed the resulting alignments via the FIMO tool within the MEME Suite to identify transcription factor motifs (https://meme-suite.org/meme/tools/fimo) using published TFBS consensus sequences for AHR(114), GR(115), Sp1(116, 117), HNF-3(118), CREB(119), and NF-I(120, 121).

### Total RNA isolation and cDNA synthesis

We extracted total RNA from both adult and embryo mummichog samples using the following protocol. Briefly, we cut ∼10mg of liver tissue or minced whole embryo pools over dry ice. We placed samples into round bottom 2mL safe lock tubes with 1mL of TRI Reagent (cat. no. T9424; Sigma-Aldrich, St. Louis, MO, USA) containing 1% (v/v) molecular biology grade β-mercaptoethanol (cat. no. M3148; Sigma-Aldrich, St. Louis, MO, USA). We homogenized samples with 5mm stainless steel beads via two 2-minute 25Hz bursts in a TissueLyser system (cat. no. 85210; Qiagen, Hilden, Germany). We transferred lysates to 1.5mL tubes, incubated samples at room temperature for 5 minutes, added 200µl of Chloroform (cat. no. 25666; Sigma-Aldrich, St. Louis, MO, USA), mixed thoroughly, transferred tissue digests into 2mL 5PRIME Phase Lock Gel Heavy tubes (cat. no. 2302830; Quantabio, Beverly, MA, USA), incubated at room temperature for 3 minutes, and centrifuged at 10,000 rpm for 15 minutes at 4°C. We transferred supernatants to new 1.5 mL tubes and precipitated RNA using 10% (v/v) 3M sodium acetate (pH 5.5) (cat. no. AM9740; Invitrogen, Waltham, MA, USA), molecular biology grade glycogen to a final concentration of 1 µg/µl (cat. no. R0561; Thermo Scientific, Waltham, MA, USA), and 100% ethanol for at least 10 minutes at −20°C. We pelleted RNA at 11,000 rpm for 10 minutes at 4°C, washed the pellet twice with 75% ethanol, and resuspended in MilliQ water. We synthesized cDNA with qScript cDNA SuperMix kits (cat. no. 95048; Quantabio, Beverly, MA, USA) on Biometra TAdvanced Thermocyclers (cat. no. 846-x-070-280; Analytik Jena, Jena, Germany), quantified cDNA using Quant-iT RNA Assay (cat. no. Q10213; Invitrogen, Waltham, MA) and Quant-iT PicoGreen dsDNA Assay Kits (cat. no. P7589; Invitrogen, Waltham, MA) to quantify both cDNA concentration and possible interfering cDNA-RNA hybrids, respectively. We used these measurements to standardize samples to a concentration of 50 ng/µl for input into real-time qPCR.

### Real-time qPCR for gene expression

We measured gene expression of *CYP1A*, *CYP1B* and *CYP1C* genes using the following protocol. Briefly, we performed qPCR on cDNA generated from total RNA isolated from adult liver tissue (*N*=5 males and *N*=5 females per population) and embryos (*N*=3 pools of 10 embryos per population per exposure status). We designed primers spanning exon-exon junctions that captured all known gene isoforms using the NCBI Primer-BLAST tool (CYP1A forward 5’- GACCTCTTTGGAGCTGGTTT −3’, reverse 5’- CCAGACCGACTTTCTCCTTGATT −3’, 124 bp product; CYP1B forward 5’- ATATTTGGAGCCAGCCAGGAC −3’, reverse 5’- GTACTTGACAAAAATGAGGATGATCCACTG −3’, 69 bp product; CYP1C forward 5’- AGCCAGGGCATGACATCAAC −3’, reverse 5’- ACACGATGACCAGGAGTTCAG −3’, 96 bp product). We ran qPCR in 10 µl reactions in triplicate with PerfeCTa SYBR Green FastMix (cat. no. 95072-012; Quantabio, Beverly, MA), 50 ng cDNA, and 500 nM forward and reverse primers on a qTower^3^G real-time thermal cycler (Analytik Jena, Jena, Germany) under the following conditions: 95°C for 30 seconds, followed by 44 cycles of 95°C for 5 seconds, annealing temperature for 30 seconds (CYP1A = 65°C, CYP1B = 60°C, CYP1C = 50°C), with a final melt curve analysis to confirm single products. We calculated cycles to threshold (C(t)) with qPCRsoftv.4.0 and averaged sample triplicate values (standard deviation ≤0.5). We were unable to validate a reference (“housekeeping”) gene as invariant across populations and exposure status; therefore, we report relative fold-change values (PAH-tolerant relative to PAH-sensitive).

### Chromatin Immunoprecipitation (ChIP) assays

We characterized histone modifications at the *CYP1A* proximal promoter-enhancer via ChIP-qPCR. Prior to ChIP experiments, we confirmed antibody specificity and efficiency with EpiCypher designer recombinant nucleosome panels (SNAP-ChIP K-MetStat Panel, cat. no. 19-1001) (**Supp. Table 27**). We performed ChIP-qPCR on four regions of *CYP1A*: three AHR-binding xenobiotic response elements (XREs) in the proximal promoter-enhancer, plus the transcription start site (TSS) for three histone modifications (H3K4me1, H3K4me3, and H3K27me3) on *N*=5 pools of 10 embryos each per population per exposure and on *N*=5 adult male liver samples per population using the following antibodies (H3K4me1, cat. no. 13-0040, Epicypher, Durham, NC; H3K4me3, cat. no. 13-0041, Epicypher, Durham, NC; H3K27me3, cat. no. MA5-1198, Invitrogen, Carlsbad, CA). We adapted the MAGnify Chromatin Immunoprecipitation System (cat. no. 49-2024, Invitrogen, Carlsbad, CA, USA) protocol for ChIP-qPCR. Briefly, we minced either 50 mg of adult liver tissue or whole embryo pools with razor blades over ice with 250 µL of chilled 1X PBS, crosslinked chromatin with 37% formaldehyde (cat. no. BP531-25, Fisher Scientific, Hampton, NJ, USA) diluted with PBS to a final concentration of 1%, incubated samples for 5 minutes, quenched reactions with 1.25 M glycine to a final concentration of 0.125 M, and homogenized samples with 0.5 mm stainless steel beads via two 2-minute, 25Hz bursts on a TissueLyser (cat. no. 85210; Qiagen, Hilden, Germany) before immediately transferring to a DNA LoBind Tube 1.5ml (cat. no. 022431021, Eppendorf, Hamburg, Germany). We pelleted samples at 1,4000 rpm for 10 minutes at 4°C, washed pellets with 500 μl chilled PBS, and centrifuged at 4,000 rpm for 10 minutes 4°C. We lysed cells with lysis buffer supplemented with protease inhibitors and mechanically sheared chromatin to 200-500bp fragments (median fragment sizes 394-536 bp in all samples) in Covaris fiber pre-slit microtubes (cat. no. 520045, Covaris, Woburn, MA, USA) with a Covaris S220 sonicator (50 μl sample, temperature 3-9°C, peak incidence power 105 W, 200 cycles per burst, duty factor 1%, for a total of 450 seconds). We verified fragment lengths on a QIAxcel Advanced system (Qiagen, Hilden, Germany) using the QIAXcel High Resolution kit (cat. no. 929002), Qiagen 15 bp-3 kb alignment marker (cat. no. 929522), and Qiagen 100-2.5 kB size marker (cat. no. 929559) using smear detection M400 method settings and stored sonicated chromatin at −80°C. We performed antibody enrichment with 1µg antibody per ChIP reaction, IgG negative controls, and Epicypher SNAP-ChIP nucleosomes as spike-in positive controls. We bound chromatin to the magnetic dynabeads, washed the bound chromatin, reversed the chromatin crosslinks, purified the chromatin, and eluted 195 µl of DNA Elution Buffer and stored samples at −20°C. We quantified enrichment of each histone PTM at each *CYP1A* region using qPCR in triplicate (10 μl DNA input, annealing temperature 50°C for all reactions, with the following primer sets: XRE3 forward 5’- CGGTTTGATCACTGCGCTCT −3’, reverse 5’- TCTCCGCGGGAGTTAAAGAT −3’, 54 bp product; XRE2 forward 5’- AACTCCCGCGGAGAGCATGC −3’, reverse 5’- AAGGTTGCGCAGTGCTGATA −3’, 69 bp product; XRE1 forward 5’- AAGGCGGTAGACACTTTGT −3’, reverse 5’- GCCATGAATGAAGTTTGGAGCA −3’, 110 bp product; TSS forward 5’- GTAGCCAATAAGATTGCGCAGC −3’, reverse 5’- AATTCCAGAGATGCGTGTCCAA −3’, 130 bp product). We evaluated ChIP performance by pooling samples from each ChIP experimental batch and running qPCR for Epicypher nucleosome barcodes specific to on-target modified nucleosomes (H3K4me1, H3K4me3, and H3K27me3) and unmodified nucleosomes (H3K4me0 and H3K27me0, respectively).

## Statistical analysis

To analyze gene expression data, we stratified raw C(t) data by embryo or adult status, by sex (for adults only), by treatment (for embryos only), and by target gene (*CYP1A, CYP1B, CYP1C*). In embryo data, we evaluated difference-in-difference (D-i-D) in each gene’s expression response to PAH challenge between PAH-tolerant and PAH-sensitive populations, denoted as [PAH-tolerant (treatment-control)] – [PAH-sensitive (treatment-control)]. In adult data, we evaluated D-i-D in sex differences in each gene’s expression between PAH-tolerant and PAH-sensitive populations, denoted as [PAH-tolerant (males - females)] – [PAH-sensitive (males - females)]. We chose Hedge’s g as a sample-sized corrected Cohen’s d value (recommended for samples N<20)(70). Here, Hedge’s g values refer to differences in least squared means in terms of standard deviations. We computed D-i-D values following Feingold’s(122) approach; we first computed Hedge’s g for each Difference (e.g., treatment-control for each population), based on raw standard deviations, and then subtracted Hedge’s g values for each population group to yield final D-i-D estimates. We applied a series of independent sample t-tests and computed Hedge’s g for treatment and sex effects within each population group (PAH-tolerant and PAH-sensitive), respectively. Next, we subtracted the Hedge’s g values between each population group, as outlined above, to yield D-i-D Hedge’s g estimates. Hedge’s g estimates of effect size can be interpreted as follows: 0.2 - small effect, 0.4-0.6 - moderate effect, >0.8 - large effect. We calculated 95% confidence intervals around the D-i-D effect sizes (if the confidence interval does not contain zero, then the effect sizes can be interpreted as statistically significant at p<0.05). We tested for differences in basal *CYP1A* expression between PAH-tolerant and PAH-sensitive embryos with an independent samples t-test. These analyses were conducted in SPSS Version 29.0 (IBM Corp., Armonk, NY), R Studio (using the “psych” R package) and Microsoft Excel.

To analyze ChIP data, we stratified fold-change values (relative to IgG negative controls) by embryo or adult status, by treatment (for embryos only), and by target region (XRE3, XRE2, XRE1, TA/TSS). Each sample ID contained four repeated measures, one for each region within the *CYP1A* proximal promoter-enhancer, and each ChIP assay was run in one of several experimental batches. Therefore, there were two dependencies in the data that may have biased results run with traditional tests (e.g., independent samples t-tests). Therefore, we ran a series of three mixed models each for embryo and adult datasets, respectively, including main effects for population group and for treatment (embryo only), an interaction term between population group and treatment (embryo only), covariates for experimental batch and gene region, and a random intercept to account for correlation among gene regions within a sample ID, nested within experimental batch. In addition, we included a three-way interaction term (population group*treatment*gene region) for embryo data and a two-way interaction term (population group*gene region) for adult data, to estimate least squares means for fold-change values for each region for a given treatment group in each population group. Given the small sample and high variability in the data, instead of directly interpreting model results, we computed statistical effect sizes (Hedge’s g) for each hypothesized difference (e.g., [PAH-tolerant – PAH-sensitive] for each gene region within adult data) and for each hypothesized D-i-D (e.g., [PAH-tolerant (treatment-control)] – [PAH-sensitive (treatment-control)] for each gene region within embryo data). We computed 95% confidence intervals for each Difference and D-i-D estimate. In addition, we computed differences in fold-change ratios (H3K27me3/H3K4me3 and H3K27me3/H3K4me1) by computing ratios from fold-change values and replicating the prior analyses. Although batches were assay-specific, in some cases, there was complete or near-complete overlap in batches when computing ratios. In these instances, we chose one assay within each batch as an indicator to control for experimental batch. We conducted a sensitivity analysis for this approach by running tests for the H3K27me3/H3K4me1 ratio twice, using either assay as the batch indicator, and observed no difference in results. These analyses were conducted in SAS Version 9.4 (Cary, NC), R Studio (using the “psych” R package) and Microsoft Excel.

## Funding and Acknowledgements

The authors thank Drs. R. Stephen Lloyd and Mitchell Turker for critical review of the manuscript. C.W. acknowledges support from the Oregon Institute of Occupational Health Sciences at Oregon Health & Science University via funds from the Division of Consumer and Business Services of the State of Oregon (ORS 656.630), as well as through a Triangle Center for Evolutionary Medicine seed grant. N.J., R.T., J.N.M., and R.T.D. were supported by the NIH/NIEHS-funded Duke University Superfund Research Center (P42ES010356). All figures were created in Biorender.

## Author Contributions

C.W. and N.J. designed the experiments, in consultation with R.T.D., J.N.M, and R.T. R.T. and N.J. caught wild fish and performed breeding and exposure experiments. N.P. and S.C. performed *CYP1A* resequencing. S.C. and E.W. performed gene expression experiments. S.C. performed ChIP experiments. The OHSU Biostatistical Design Program staff performed statistical analyses. C.W. wrote the manuscript and all co-authors reviewed and edited the manuscript. C.W. oversaw the project.

## References

1. Toxicological profile for polycyclic aromatic hydrocarbons. Agency for Toxic Substances and Disease Registry, U.S. Department of Health and Human Services, Public Health Service Atlanta, Georgia (1995).

2. Diesel and Gasoline Engine Exhausts and Some Nitroarenes. IARC Monographs on the Evaluation of Carcinogenic Risks to Humans. IARC Monogr Eval Carcinog Risks Hum 105, 9–699 (2014).

3. IARC Monographs on the Identification of Carcinogenic Hazards to Humans. Report of the Advisory Group to Recommend Priorities for the IARC Monographs During 2020-2024. (2019).

4. Outdoor Air Pollution: IARC Working Group on the Evaluation of Carcinogenic Risks to Humans. IARC Monogr Eval Carcinog Risks Hum 109, 9-444 (2016).

5. Tobacco Smoke and Involuntary Smoking: IARC Working Group on the Evaluation of Carcinogenic Risks to Humans. IARC Monogr Eval Carcinog Risks Hum 83, 1–1438 (2004).

6. D. M. DeMarini, W. P. Linak, Mutagenicity and carcinogenicity of combustion emissions are impacted more by combustor technology than by fuel composition: A brief review. Environ Mol Mutagen 63, 135–150 (2022).

7. Spreading like Wildfire - The Rising Threat of Extraordinary Landscape Fires. A UNEP Rapid Response Assessment. Nairobi. (2022).

8. R. E. Albert, Comparative carcinogenic potencies of particulates from diesel engine exhausts, coke oven emissions, roofing tar aerosols and cigarette smoke. Environ Health Perspect 47, 339–341 (1983).

9. L. T. Cupitt, W. G. Glen, J. Lewtas, Exposure and risk from ambient particle-bound pollution in an airshed dominated by residential wood combustion and mobile sources. Environ Health Perspect 102 Suppl 4, 75–84 (1994).

10. Tumour Site Concordance and Mechanisms of Carcinogenesis, R. A. Baan, B. W. Stewart, K. Straif, Eds. (Lyon (FR), 2019).

11. The Lancet Editorial, Lung cancer: some progress, but still a lot more to do. Lancet 394, 1880 (2019).

12. C. Frick et al., Quantitative estimates of preventable and treatable deaths from 36 cancers worldwide: a population-based study. Lancet Glob Health 11, e1700–e1712 (2023).

13. R. Doll, A. B. Hill, Smoking and carcinoma of the lung; preliminary report. Br Med J 2, 739–748 (1950).

14. K. Yoshida et al., Tobacco smoking and somatic mutations in human bronchial epithelium. Nature 578, 266–272 (2020).

15. X. Dai et al., Health effects associated with smoking: a Burden of Proof study. Nat Med 28, 2045–2055 (2022).

16. G. B. D. C. R. F. Collaborators, The global burden of cancer attributable to risk factors, 2010-19: a systematic analysis for the Global Burden of Disease Study 2019. Lancet 400, 563–591 (2022).

17. W. Hill et al., Lung adenocarcinoma promotion by air pollutants. Nature 616, 159–167 (2023).

18. H. Sung et al., Global Cancer Statistics 2020: GLOBOCAN Estimates of Incidence and Mortality Worldwide for 36 Cancers in 185 Countries. CA Cancer J Clin 71, 209–249 (2021).

19. J. Ferlay et al., Cancer statistics for the year 2020: An overview. Int J Cancer 10.1002/ijc.33588 (2021).

20. J. Ferlay, Ervik M., Lam F., Laversanne M., Colombet M., Mery L., Pineros M., Znaor A., Soerjomataram I., Bray F., Global Cancer Observatory: Cancer Today. International Agency for Research on Cancer, *Lyon, France*. (2024).

21. R. L. Siegel, K. D. Miller, N. S. Wagle, A. Jemal, Cancer statistics, 2023. CA Cancer J Clin 73, 17–48 (2023).

22. R. Stading, G. Gastelum, C. Chu, W. Jiang, B. Moorthy, Molecular mechanisms of pulmonary carcinogenesis by polycyclic aromatic hydrocarbons (PAHs): Implications for human lung cancer. Semin Cancer Biol 76, 3–16 (2021).

23. G. Gastelum et al., Polycyclic Aromatic Hydrocarbon-induced Pulmonary Carcinogenesis in Cytochrome P450 (CYP)1A1- and 1A2-Null Mice: Roles of CYP1A1 and CYP1A2. Toxicol Sci 177, 347–361 (2020).

24. T. Shimada, Y. Oda, E. M. Gillam, F. P. Guengerich, K. Inoue, Metabolic activation of polycyclic aromatic hydrocarbons and other procarcinogens by cytochromes P450 1A1 and P450 1B1 allelic variants and other human cytochromes P450 in Salmonella typhimurium NM2009. Drug Metab Dispos 29, 1176–1182 (2001).

25. T. Shimada, Y. Fujii-Kuriyama, Metabolic activation of polycyclic aromatic hydrocarbons to carcinogens by cytochromes P450 1A1 and 1B1. Cancer Sci 95, 1–6 (2004).

26. T. Shimada et al., Arylhydrocarbon receptor-dependent induction of liver and lung cytochromes P450 1A1, 1A2, and 1B1 by polycyclic aromatic hydrocarbons and polychlorinated biphenyls in genetically engineered C57BL/6J mice. Carcinogenesis 23, 1199–1207 (2002).

27. Y. Matsumoto et al., Aryl hydrocarbon receptor plays a significant role in mediating airborne particulate-induced carcinogenesis in mice. Environ Sci Technol 41, 3775–3780 (2007).

28. K. L. Willett, P. R. Gardinali, J. L. Sericano, T. L. Wade, S. H. Safe, Characterization of the H4IIE rat hepatoma cell bioassay for evaluation of environmental samples containing polynuclear aromatic hydrocarbons (PAHs). Arch Environ Contam Toxicol 32, 442–448 (1997).

29. M. Till, D. Riebniger, H. J. Schmitz, D. Schrenk, Potency of various polycyclic aromatic hydrocarbons as inducers of CYP1A1 in rat hepatocyte cultures. Chem Biol Interact 117, 135–150 (1999).

30. M. Machala, J. Vondracek, L. Blaha, M. Ciganek, J. V. Neca, Aryl hydrocarbon receptor-mediated activity of mutagenic polycyclic aromatic hydrocarbons determined using in vitro reporter gene assay. Mutat Res 497, 49–62 (2001).

31. R. E. Kouri et al., Correlation of inducibility of aryl hydrocarbon hydroxylase with susceptibility to 3-methylcholanthrene-induced lung cancers. Cancer Lett 9, 277–284 (1980).

32. O. Pelkonen, A. R. Boobis, R. C. Levitt, R. E. Kouri, D. W. Nebert, Genetic differences in the metabolic activation of benzo[a]pyrene in mice. Attempts to correlate tumorigenesis with binding of reactive intermediates to DNA and with mutagenesis in vitro. Pharmacology 18, 281–293 (1979).

33. J. Malhotra, M. Malvezzi, E. Negri, C. La Vecchia, P. Boffetta, Risk factors for lung cancer worldwide. Eur Respir J 48, 889–902 (2016).

34. P. B. Bach et al., Variations in lung cancer risk among smokers. J Natl Cancer Inst 95, 470–478 (2003).

35. A. A. G. Gabriel et al., Genetic Analysis of Lung Cancer and the Germline Impact on Somatic Mutation Burden. J Natl Cancer Inst 114, 1159–1166 (2022).

36. D. Sengupta et al., A comprehensive meta-analysis and a case-control study give insights into genetic susceptibility of lung cancer and subgroups. Sci Rep 11, 14572 (2021).

37. N. Ezzeldin et al., Genetic polymorphisms of human cytochrome P450 CYP1A1 in an Egyptian population and tobacco-induced lung cancer. Genes Environ 39, 7 (2017).

38. L. P. Zhang, C. P. Wang, L. H. Li, Y. F. Tang, W. C. Li, The interaction between smoking and CYP1A1 MspI polymorphism on lung cancer: a meta-analysis in the Chinese population. Eur J Cancer Care (Engl) 26 (2017).

39. X. F. He et al., Association between the CYP1A1 T3801C polymorphism and risk of cancer: evidence from 268 case-control studies. Gene 534, 324–344 (2014).

40. K. Nakachi, K. Imai, S. Hayashi, J. Watanabe, K. Kawajiri, Genetic susceptibility to squamous cell carcinoma of the lung in relation to cigarette smoking dose. Cancer Res 51, 5177–5180 (1991).

41. K. Kawajiri, K. Nakachi, K. Imai, S. Hayashi, J. Watanabe, Individual differences in lung cancer susceptibility in relation to polymorphisms of P-450IA1 gene and cigarette dose. Princess Takamatsu Symp 21, 55–61 (1990).

42. D. W. Nebert, R. A. McKinnon, A. Puga, Human drug-metabolizing enzyme polymorphisms: effects on risk of toxicity and cancer. DNA Cell Biol 15, 273–280 (1996).

43. P. P. Shah, K. Saurabh, M. C. Pant, N. Mathur, D. Parmar, Evidence for increased cytochrome P450 1A1 expression in blood lymphocytes of lung cancer patients. Mutat Res 670, 74–78 (2009).

44. D. Schwarz, P. Kisselev, I. Cascorbi, W. H. Schunck, I. Roots, Differential metabolism of benzo[a]pyrene and benzo[a]pyrene-7,8-dihydrodiol by human CYP1A1 variants. Carcinogenesis 22, 453–459 (2001).

45. S. Hayashi, J. Watanabe, K. Nakachi, K. Kawajiri, Genetic linkage of lung cancer-associated MspI polymorphisms with amino acid replacement in the heme binding region of the human cytochrome P450IA1 gene. J Biochem 110, 407–411 (1991).

46. G. Cosma, F. Crofts, E. Taioli, P. Toniolo, S. Garte, Relationship between genotype and function of the human CYP1A1 gene. J Toxicol Environ Health 40, 309–316 (1993).

47. N. Ishibe et al., Susceptibility to lung cancer in light smokers associated with CYP1A1 polymorphisms in Mexican- and African-Americans. Cancer Epidemiol Biomarkers Prev 6, 1075–1080 (1997).

48. T. Oyama et al., Cytochrome P450 expression (CYP) in non-small cell lung cancer. Front Biosci 12, 2299–2308 (2007).

49. T. Oyama et al., Increased cytochrome P450 and aryl hydrocarbon receptor in bronchial epithelium of heavy smokers with non-small cell lung carcinoma carries a poor prognosis. Front Biosci 12, 4497–4503 (2007).

50. S. Anttila et al., Methylation of cytochrome P4501A1 promoter in the lung is associated with tobacco smoking. Cancer Res 63, 8623–8628 (2003).

51. W. D. J. Van Veld P.A., Evidence for depression of cytochrome P4501A in a population of chemically resistant mummichog (Fundulus heteroclitus). Environ Sci 3, 221–234 (1995).

52. J. N. Meyer, D. E. Nacci, R. T. Di Giulio, Cytochrome P4501A (CYP1A) in killifish (Fundulus heteroclitus): heritability of altered expression and relationship to survival in contaminated sediments. Toxicol Sci 68, 69-81 (2002).

53. J. N. Meyer, D. M. Wassenberg, S. I. Karchner, M. E. Hahn, R. T. Di Giulio, Expression and inducibility of aryl hydrocarbon receptor pathway genes in wild-caught killifish (Fundulus heteroclitus) with different contaminant-exposure histories. Environ Toxicol Chem 22, 2337–2343 (2003).

54. L. P. Wills, C. W. Matson, C. D. Landon, R. T. Di Giulio, Characterization of the recalcitrant CYP1 phenotype found in Atlantic killifish (Fundulus heteroclitus) inhabiting a Superfund site on the Elizabeth River, VA. Aquat Toxicol 99, 33–41 (2010).

55. B. W. Clark, A. J. Bone, R. T. Di Giulio, Resistance to teratogenesis by F1 and F2 embryos of PAH-adapted Fundulus heteroclitus is strongly inherited despite reduced recalcitrance of the AHR pathway. Environ Sci Pollut Res Int 21, 13898–13908 (2014).

56. R. T. Di Giulio, B. W. Clark, The Elizabeth River Story: A Case Study in Evolutionary Toxicology. J Toxicol Environ Health B Crit Rev 18, 259–298 (2015).

57. N. M. Reid et al., The genomic landscape of rapid repeated evolutionary adaptation to toxic pollution in wild fish. Science 354, 1305–1308 (2016).

58. L. P. Wills et al., Comparative chronic liver toxicity of benzo[a]pyrene in two populations of the atlantic killifish (Fundulus heteroclitus) with different exposure histories. Environ Health Perspect 118, 1376–1381 (2010).

59. D. M. Wassenberg, R. T. Di Giulio, Synergistic embryotoxicity of polycyclic aromatic hydrocarbon aryl hydrocarbon receptor agonists with cytochrome P4501A inhibitors in Fundulus heteroclitus. Environ Health Perspect 112, 1658–1664 (2004).

60. J. N. Meyer, Di Giulio, R.T., Heritable adaptation and fitness costs in killifish (Fundulus heteroclitus) inhabiting a polluted estuary. Ecological Applications 13, 490–503 (2003).

61. P. Voigt, W. W. Tee, D. Reinberg, A double take on bivalent promoters. Genes Dev 27, 1318–1338 (2013).

62. M. Ptashne, Binding reactions: epigenetic switches, signal transduction and cancer. Curr Biol 19, R234–241 (2009).

63. M. Ptashne, On the use of the word ‘epigenetic’. Curr Biol 17, R233–236 (2007).

64. S. Henikoff, J. M. Greally, Epigenetics, cellular memory and gene regulation. Curr Biol 26, R644–648 (2016).

65. B. E. Bernstein, A. Meissner, E. S. Lander, The mammalian epigenome. Cell 128, 669–681 (2007).

66. A. Haghani et al., DNA methylation networks underlying mammalian traits. Science 381, eabq5693 (2023).

67. Y. Takahashi et al., Transgenerational inheritance of acquired epigenetic signatures at CpG islands in mice. Cell 186, 715–731 e719 (2023).

68. N. Reveron-Gomez et al., Accurate Recycling of Parental Histones Reproduces the Histone Modification Landscape during DNA Replication. Mol Cell 72, 239–249 e235 (2018).

69. V. Flury et al., Recycling of modified H2A-H2B provides short-term memory of chromatin states. Cell 186, 1050–1065 e1019 (2023).

70. D. Lakens, Calculating and reporting effect sizes to facilitate cumulative science: a practical primer for t-tests and ANOVAs. Front Psychol 4, 863 (2013).

71. R. L. Jirtle, M. K. Skinner, Environmental epigenomics and disease susceptibility. Nat Rev Genet 8, 253–262 (2007).

72. A. R. Timme-Laragy, J. N. Meyer, R. A. Waterland, R. T. Di Giulio, Analysis of CpG methylation in the killifish CYP1A promoter. Comp Biochem Physiol C Toxicol Pharmacol 141, 406–411 (2005).

73. H. Wang et al., H3K4me3 regulates RNA polymerase II promoter-proximal pause-release. Nature 615, 339–348 (2023).

74. N. Kubo et al., H3K4me1 facilitates promoter-enhancer interactions and gene activation during embryonic stem cell differentiation. Mol Cell 10.1016/j.molcel.2024.02.030 (2024).

75. R. T. Coleman, G. Struhl, Causal role for inheritance of H3K27me3 in maintaining the OFF state of a Drosophila HOX gene. Science 356 (2017).

76. A. Sankar et al., Histone editing elucidates the functional roles of H3K27 methylation and acetylation in mammals. Nat Genet 54, 754–760 (2022).

77. A. Scacchetti, R. Bonasio, Histone gene editing probes functions of H3K27 modifications in mammals. Nat Genet 54, 746–747 (2022).

78. B. E. Bernstein et al., A bivalent chromatin structure marks key developmental genes in embryonic stem cells. Cell 125, 315–326 (2006).

79. N. Nalabothula et al., Archaeal nucleosome positioning in vivo and in vitro is directed by primary sequence motifs. BMC Genomics 14, 391 (2013).

80. J. N. Meyer, D. C. Volz, J. H. Freedman, R. T. Di Giulio, Differential display of hepatic mRNA from killifish (Fundulus heteroclitus) inhabiting a Superfund estuary. Aquat Toxicol 73, 327–341 (2005).

81. D. R. Burrill, M. C. Inniss, P. M. Boyle, P. A. Silver, Synthetic memory circuits for tracking human cell fate. Genes Dev 26, 1486–1497 (2012).

82. V. Azuara et al., Chromatin signatures of pluripotent cell lines. Nat Cell Biol 8, 532–538 (2006).

83. E. Shema et al., Single-molecule decoding of combinatorially modified nucleosomes. Science 352, 717–721 (2016).

84. E. Blanco, M. Gonzalez-Ramirez, A. Alcaine-Colet, S. Aranda, L. Di Croce, The Bivalent Genome: Characterization, Structure, and Regulation. Trends Genet 36, 118–131 (2020).

85. A. P. Bracken, N. Dietrich, D. Pasini, K. H. Hansen, K. Helin, Genome-wide mapping of Polycomb target genes unravels their roles in cell fate transitions. Genes Dev 20, 1123–1136 (2006).

86. G. Caretti, M. Di Padova, B. Micales, G. E. Lyons, V. Sartorelli, The Polycomb Ezh2 methyltransferase regulates muscle gene expression and skeletal muscle differentiation. Genes Dev 18, 2627–2638 (2004).

87. K. Cui et al., Chromatin signatures in multipotent human hematopoietic stem cells indicate the fate of bivalent genes during differentiation. Cell Stem Cell 4, 80–93 (2009).

88. E. Ezhkova et al., Ezh2 orchestrates gene expression for the stepwise differentiation of tissue-specific stem cells. Cell 136, 1122–1135 (2009).

89. P. B. Rahl et al., c-Myc regulates transcriptional pause release. Cell 141, 432–445 (2010).

90. I. H. Su et al., Ezh2 controls B cell development through histone H3 methylation and Igh rearrangement. Nat Immunol 4, 124–131 (2003).

91. M. G. Lee et al., Demethylation of H3K27 regulates polycomb recruitment and H2A ubiquitination. Science 318, 447–450 (2007).

92. K. Agger et al., UTX and JMJD3 are histone H3K27 demethylases involved in HOX gene regulation and development. Nature 449, 731–734 (2007).

93. T. S. Mikkelsen et al., Genome-wide maps of chromatin state in pluripotent and lineage-committed cells. Nature 448, 553–560 (2007).

94. F. Mohn et al., Lineage-specific polycomb targets and de novo DNA methylation define restriction and potential of neuronal progenitors. Mol Cell 30, 755–766 (2008).

95. A. Harikumar, E. Meshorer, Chromatin remodeling and bivalent histone modifications in embryonic stem cells. EMBO Rep 16, 1609–1619 (2015).

96. Y. Tian et al., Early Life Short-Term Exposure to Polychlorinated Biphenyl 126 in Mice Leads to Metabolic Dysfunction and Microbiota Changes in Adulthood. Int J Mol Sci 23 (2022).

97. W. Jiang et al., Persistent induction of cytochrome P450 (CYP)1A enzymes by 3-methylcholanthrene in vivo in mice is mediated by sustained transcriptional activation of the corresponding promoters. Biochem Biophys Res Commun 390, 1419–1424 (2009).

98. I. S. Fazili et al., Persistent induction of cytochrome P4501A1 in human hepatoma cells by 3-methylcholanthrene: evidence for sustained transcriptional activation of the CYP1A1 promoter. J Pharmacol Exp Ther 333, 99–109 (2010).

99. H. Z. Amenya, C. Tohyama, S. Ohsako, Dioxin induces Ahr-dependent robust DNA demethylation of the Cyp1a1 promoter via Tdg in the mouse liver. Sci Rep 6, 34989 (2016).

100. H. P. Ciolino, G. C. Yeh, Inhibition of aryl hydrocarbon-induced cytochrome P-450 1A1 enzyme activity and CYP1A1 expression by resveratrol. Mol Pharmacol 56, 760–767 (1999).

101. H. P. Ciolino, G. C. Yeh, The flavonoid galangin is an inhibitor of CYP1A1 activity and an agonist/antagonist of the aryl hydrocarbon receptor. Br J Cancer 79, 1340–1346 (1999).

102. H. P. Ciolino, P. J. Daschner, G. C. Yeh, Dietary flavonols quercetin and kaempferol are ligands of the aryl hydrocarbon receptor that affect CYP1A1 transcription differentially. Biochem J 340 (Pt 3), 715–722 (1999).

103. H. P. Ciolino, T. T. Wang, G. C. Yeh, Diosmin and diosmetin are agonists of the aryl hydrocarbon receptor that differentially affect cytochrome P450 1A1 activity. Cancer Res 58, 2754–2760 (1998).

104. M. S. Chua et al., Role of Cyp1A1 in modulation of antitumor properties of the novel agent 2-(4-amino-3-methylphenyl)benzothiazole (DF 203, NSC 674495) in human breast cancer cells. Cancer Res 60, 5196–5203 (2000).

105. E. Brantley et al., Fluorinated 2-(4-amino-3-methylphenyl)benzothiazoles induce CYP1A1 expression, become metabolized, and bind to macromolecules in sensitive human cancer cells. Drug Metab Dispos 32, 1392–1401 (2004).

106. T. D. Bradshaw, A. D. Westwell, The development of the antitumour benzothiazole prodrug, Phortress, as a clinical candidate. Curr Med Chem 11, 1009–1021 (2004).

107. V. Trapani et al., DNA damage and cell cycle arrest induced by 2-(4-amino-3-methylphenyl)-5-fluorobenzothiazole (5F 203, NSC 703786) is attenuated in aryl hydrocarbon receptor deficient MCF-7 cells. Br J Cancer 88, 599–605 (2003).

108. A. I. Loaiza-Perez et al., Aryl hydrocarbon receptor activation of an antitumor aminoflavone: basis of selective toxicity for MCF-7 breast tumor cells. Mol Cancer Ther 3, 715–725 (2004).

109. L. H. Meng et al., Activation of aminoflavone (NSC 686288) by a sulfotransferase is required for the antiproliferative effect of the drug and for induction of histone gamma-H2AX. Cancer Res 66, 9656–9664 (2006).

110. V. Androutsopoulos, R. R. Arroo, J. F. Hall, S. Surichan, G. A. Potter, Antiproliferative and cytostatic effects of the natural product eupatorin on MDA-MB-468 human breast cancer cells due to CYP1-mediated metabolism. Breast Cancer Res 10, R39 (2008).

111. V. Androutsopoulos, N. Wilsher, R. R. Arroo, G. A. Potter, Bioactivation of the phytoestrogen diosmetin by CYP1 cytochromes P450. Cancer Lett 274, 54–60 (2009).

112. V. P. Androutsopoulos, S. Mahale, R. R. Arroo, G. Potter, Anticancer effects of the flavonoid diosmetin on cell cycle progression and proliferation of MDA-MB 468 breast cancer cells due to CYP1 activation. Oncol Rep 21, 1525–1528 (2009).

113. M. Fang et al., Effect-directed analysis of Elizabeth River porewater: developmental toxicity in zebrafish (Danio rerio). Environ Toxicol Chem 33, 2767–2774 (2014).

114. W. H. Powell et al., Cloning and analysis of the CYP1A promoter from the atlantic killifish (Fundulus heteroclitus). Mar Environ Res 58, 119–124 (2004).

115. H. M. Jantzen et al., Cooperativity of glucocorticoid response elements located far upstream of the tyrosine aminotransferase gene. Cell 49, 29–38 (1987).

116. W. S. Dynan, R. Tjian, The promoter-specific transcription factor Sp1 binds to upstream sequences in the SV40 early promoter. Cell 35, 79–87 (1983).

117. J. A. Segal, J. L. Barnett, D. L. Crawford, Functional analyses of natural variation in Sp1 binding sites of a TATA-less promoter. J Mol Evol 49, 736–749 (1999).

118. G. Zeruth, R. S. Pollenz, Functional analysis of cis-regulatory regions within the dioxin-inducible CYP1A promoter/enhancer region from zebrafish (Danio rerio). Chem Biol Interact 170, 100–113 (2007).

119. M. R. Montminy, K. A. Sevarino, J. A. Wagner, G. Mandel, R. H. Goodman, Identification of a cyclic-AMP-responsive element within the rat somatostatin gene. Proc Natl Acad Sci U S A 83, 6682–6686 (1986).

120. R. M. Gronostajski, S. Adhya, K. Nagata, R. A. Guggenheimer, J. Hurwitz, Site-specific DNA binding of nuclear factor I: analyses of cellular binding sites. Mol Cell Biol 5, 964–971 (1985).

121. A. Yanagida, K. Sogawa, K. I. Yasumoto, Y. Fujii-Kuriyama, A novel cis-acting DNA element required for a high level of inducible expression of the rat P-450c gene. Mol Cell Biol 10, 1470–1475 (1990).

122. A. Feingold, Effect sizes for growth-modeling analysis for controlled clinical trials in the same metric as for classical analysis. Psychol Methods 14, 43–53 (2009).

