## Supplemental Figures for "An epigenetic memory at the *CYP1A* gene in cancer-resistant, pollution-adapted killifish"

**This PDF file includes:**

- Supporting text
- Supporting text references
- Figures S1 to S13

**Other supporting materials for this manuscript include the following:**

- Tables S1-27

### Supporting text

#### AhR sequence motifs in the *CYP1A* promoter-enhancer are conserved

The *CYP1A* gene in PAH-tolerant mummichog shows marked sequence deviation from the PAH-sensitive gene[1], which suggests the possibility of genetic contribution to the *CYP1A* memory phenotype. We reasoned that genetic variation in AHR sequence motifs (xenobiotic response elements, or XREs) could alter AHR affinity for the *CYP1A* gene's proximal promoter-enhancer. Therefore, we re-sequenced *CYP1A*, including 7Kb upstream of the transcription start site (TSS), in four adult, wild-caught mummichog, one male and one female each from a PAH-tolerant population and a PAH-sensitive population (**Fig. 1A**). Although we observed high sequence variation at *CYP1A* in fish from the two populations (**Supp. Fig. 1, Supp. Table 1**), which is consistent with prior reports[1], we did not observe any sequence changes in any of the three well-characterized, functional XREs[2] in the proximal promoter-enhancer (located at positions -784, -735, and -194 from the TSS, respectively), except for a single base pair change in a PAH-sensitive male (**Fig. 1B, Supp. Fig. 1, Supp. Table 1**). These three XREs are the best-characterized AHR binding sites in *CYP1A*; however, it is possible that additional XREs are present that regulate expression of this gene. To evaluate this possibility, we screened the gene and ~7kB upstream sequence for additional XRE motifs. We identified one additional consensus and three additional partial XRE motifs (**Supp. Fig. 1, Supp. Table 1**), but none are likely to explain observed population differences in *CYP1A* inducibility because all additional partial motifs were absent in one of two PAH-sensitive fish that we tested, despite known *CYP1A* inducibility in these fish (**Supp. Fig. 1, Supp. Table 1)**. The single additional consensus motif was present in all fish tested, as well as the reference, (**Supp. Fig. 1, Supp. Table 1**) and is unlikely to explain phenotypic differences between the Virginia populations. Therefore, decreased sequence affinity for AHR in the *CYP1A* promoter-enhancer region does not explain reduced induction of *CYP1A* in PAH-tolerant fish.

Next, we evaluated sequence motifs for transcription factors which are known to bind cooperatively with AhR or to moderate AHR induction of *CYP1A*, including specificity protein 1 (Sp1)[3,4], glucocorticoid receptor (GR)[5-7], cAMP response element-binding protein (CREB)[8], nuclear factor I (NF-I)[9,10], and hepatocyte nuclear factor-3 (HNF-3)[8]. We reasoned that if one or more of these motifs were mutated in PAH-tolerant fish and these mutations altered sequence affinity for one or more of these factors under certain cellular conditions, then these mutations could explain the reversible blunting of AHR induction of *CYP1A*. We identified several motifs within the *CYP1A* gene body and proximal promoter-enhancer that showed mutation patterns consistent with altered transcriptional regulation. Specifically, in the gene body, we identified four full-length, consensus co-factor motifs: one exonic Sp1 site, one intronic HNF-3 site and two exonic NF-I sites (**Supp. Fig. 1, Supp. Table 1**). The Sp1 site was present only in PAH-tolerant sequences, which is consistent with gain of novel positive Sp1 regulation of AhR binding in fish with evolved tolerance to PAH (**Supp. Fig. 1, Supp. Table 1**). The HNF-3 site was present only in PAH-sensitive and reference sequences, which is consistent with loss of HNF-3 positive regulation of AhR binding in fish with evolved tolerance to PAH (**Supp. Fig. 1, Supp. Table 1**). One of the exonic NF-I sites was perfectly conserved in all sequences; the second NF-I site was conserved across all sequences except in the PAH-sensitive female (**Supp. Fig. 1, Supp. Table 1**). This second finding may reflect a standing genetic variant in the PAH-sensitive population that is lost in the tolerant population. NF-I binding decreases *CYP1A* expression[10], so it is possible that this additional NF-I site caused decreased *CYP1A* expression that conferred a fitness advantage in a PAH contaminated environment and increased in frequency in PAH-tolerant fish. In the proximal promoter-enhancer (1500 bp upstream from the TSS), which is known to regulate basal[11] and inducible *CYP1A* expression[2], we identified one full-length and 11 partial Sp1 sites, including three partial Sp1 sites near the TSS; one full-length HNF-3 site; nine GR half-sites; and one CREB half-site (**Supp. Fig. 1, Supp. Table 1**). Neither proximal promoter-enhancer region full-length Sp1 and HNF-3 sites is implicated in the *CYP1A* memory phenotype, since both were completely conserved across all four fish and the reference sequence (**Supp. Fig. 1, Supp. Table 1**). This result is similar for the XREs found within the promoter-enhancer region, indicating strong conservation of regulatory sequences immediately upstream of the gene (**Supp. Fig. 1, Supp. Table 1**); neither of these motifs likely contributes to population differences in *CYP1A* inducibility because they are conserved across fish from both populations. However, three of the partial Sp1 sites, including the one nearest to the TSS, and one GR half-site in the proximalpromoter enhancer, were present in one or both PAH-sensitive fish but were absent in one or both tolerant fish, which is consistent with partial loss of regulatory control at *CYP1A* in PAH-tolerant fish. We did not observe any additional, informative motifs in sequence farther upstream of the proximal promoter-enhancer. We observed only two additional conserved, full-length HNF-3 motifs, one additional conserved, full-length GR motif, and one additional conserved, full-length NF-I motif (**Supp. Fig. 1, Supp. Table 1**). Like other highly conserved motifs, none of these motifs is likely responsible for population differences in *CYP1A* inducibility, since they are conserved across fish from both populations. In summary, most predicted co-factor motifs with likely regulatory potential were conserved across all four fish and showed no deviation from the reference genome (**Supp. Fig. 1, Supp. Table 1**). Therefore, genetic variation in transcription factor binding sites is unlikely to be solely responsible for the memory at *CYP1A*. However, we did observe limited evidence for loss of regulatory sequence motifs in PAH-tolerant fish that is consistent with combined genetic and epigenetic effects at this locus.

Supporting text references


1 Reid, N. M. *et al.* The genomic landscape of rapid repeated evolutionary adaptation to toxic pollution in wild fish. *Science* **354**, 1305-1308, doi:10.1126/science.aah4993 (2016).

2 Powell, W. H. *et al.* Cloning and analysis of the CYP1A promoter from the atlantic killifish (Fundulus heteroclitus). *Mar Environ Res* **58**, 119-124, doi:10.1016/j.marenvres.2004.03.005 (2004).

3 Zhang, W., Shields, J. M., Sogawa, K., Fujii-Kuriyama, Y. & Yang, V. W. The gut-enriched Kruppel-like factor suppresses the activity of the CYP1A1 promoter in an Sp1-dependent fashion. *J Biol Chem* **273**, 17917-17925, doi:10.1074/jbc.273.28.17917 (1998).

4 Fujii-Kuriyama, Y., Imataka, H., Sogawa, K., Yasumoto, K. & Kikuchi, Y. Regulation of CYP1A1 expression. *FASEB J* **6**, 706-710, doi:10.1096/fasebj.6.2.1537460 (1992).

5 Vrzal, R. *et al.* Dexamethasone controls aryl hydrocarbon receptor (AhR)-mediated CYP1A1 and CYP1A2 expression and activity in primary cultures of human hepatocytes. *Chem Biol Interact* **179**, 288-296, doi:10.1016/j.cbi.2008.10.035 (2009).

6 Linder, M. W., Falkner, K. C., Srinivasan, G., Hines, R. N. & Prough, R. A. Role of canonical glucocorticoid responsive elements in modulating expression of genes regulated by the arylhydrocarbon receptor. *Drug Metab Rev* **31**, 247-271, doi:10.1081/dmr-100101917 (1999).

7 Harvey, J. L., Paine, A. J. & Wright, M. C. Disruption of endogenous regulator homeostasis underlies the mechanism of rat CYP1A1 mRNA induction by metyrapone. *Biochem J* **331 ( Pt 1)**, 273-281, doi:10.1042/bj3310273 (1998).

8 Zeruth, G. & Pollenz, R. S. Functional analysis of cis-regulatory regions within the dioxin-inducible CYP1A promoter/enhancer region from zebrafish (Danio rerio). *Chem Biol Interact* **170**, 100-113, doi:10.1016/j.cbi.2007.07.003 (2007).

9 Morel, Y., Mermod, N. & Barouki, R. An autoregulatory loop controlling CYP1A1 gene expression: role of H(2)O(2) and NFI. *Mol Cell Biol* **19**, 6825-6832, doi:10.1128/MCB.19.10.6825 (1999).

10 Morel, Y. & Barouki, R. Down-regulation of cytochrome P450 1A1 gene promoter by oxidative stress. Critical contribution of nuclear factor 1. *J Biol Chem* **273**, 26969-26976, doi:10.1074/jbc.273.41.26969 (1998).

11 Yanagida, A., Sogawa, K., Yasumoto, K. I. & Fujii-Kuriyama, Y. A novel cis-acting DNA element required for a high level of inducible expression of the rat P-450c gene. *Mol Cell Biol* **10**, 1470-1475, doi:10.1128/mcb.10.4.1470-1475.1990 (1990).

**Supplemental Figure 1. *Fundulus heteroclitus CYP1A* gene sequence.** We re-sequenced the *CYP1A* gene, as well as 7 kilobases upstream of the transcription start site (chromosomal coordinates chr1:1,265,805-1,276,398 (accession NW_012234324.1), via primer walking in four adult fish of species *Fundulus heteroclitus*, including one male and one female each from two populations: a PAH-sensitive population from the King's Creek tributary of the Severn River in Virginia, USA, and a PAH-tolerant population from the Republic tributary of the Elizabeth River in Virginia, USA. All four sequences are shown aligned to the reference sequence derived from a related *F. heteroclitus* population in Maine. The sequences are annotated as follows. Exonic sequence is colored green, intronic sequence is colored red, the TATAA box is colored fuchsia, and transcription start site is colored blue. Insertions are highlighted in green, deletions are highlighted in red, and deviations from reference are highlighted in cerulean. Transcription factor motifs are colored for easy comparisons among fish sequences. Xenobiotic response elements (XREs) are purple, glucocorticoid response elements (GREs) are orange, Sp1 motifs are light blue, CREB motifs are royal blue, HNF-3 motifs are brown, HNF-1 motifs are hot pink. Overlapping motifs are distinguished by underlining the first motif and italicizing the second motif.

```
PAH-sensitive female---------        ----------|AAGGTAA-TTTATTTTTTTATGATCAACCCTCAAAATCA
PAH-sensitive male-----------        ----------|AAGGTAA-TTTATTTTTT-ATGATCAACCCTCAAAATCA
PAH-tolerant female----------        ----------|AAGGTAA-TTTATTTTTT-ATGATCAACCCTCAAAATCA
PAH-tolerant male------------        ----------|AAGGTAAATTTATTTTTT-ATGATCAACCCTCAAAATCA
CYP1Arefseq------------------        ----------|AAGGTAAATTTATTTTTTTATGATCAACCCTCAAAATCA
                                               |******* ********** ********************

PAH-sensitive female---------        GTCACAAATAAAAAAAAAAC--TCTTTTTGGGCTTGTATGCACAGGAATA
PAH-sensitive male-----------        GTCACAAATAAAAATAAACCCCTCTTTTTGGGCTTGTATGCACAGGAATA
PAH-tolerant female----------        GTCACAAATAAAAATAAACCC-TCTTTTTGGGCTTGTATGCACAGGAATA
PAH-tolerant male------------        GTCACAAATAAAAATAAACCC-TCTTTTTGGGCTTGTATGCACAGGAATA
CYP1Arefseq------------------        ATCACAAATAAAAAGAAACCC-TCTTTTTGGGCTTGTATGCACAGGAATA
                                      ************* *** *  ****************************
```

```
PAH-sensitive female---------       GAGTCACATACTAAGAATGTGATCCACTCCTGCTGTTAGGTTGTGTGCGT
PAH-sensitive male-----------       GAGTCACATATTAAGAATGTGATCCACTCCTGCTGTTAGGTTGTGTGCGT
PAH-tolerant female----------       GAGTCACATACTAAGAATGTGATCCACTCCTGCTGTTAGGTTGTGTGCGT
PAH-tolerant male------------       GAGTCACATACTAAGAATGTGATCCACTCCTGCTGTTAGGTTGTGTGCGT
CYP1Arefseq------------------       GAGTCACATACTAAGAATGTGATCCACTCCTGCTGTTAGGTTGCGTGCGT
                                    ********** ******************************** ******

PAH-sensitive female---------       GACAACCTTTGTATAACTTGACAACCCTCTCA-GTCACTGCGCACACTTT
PAH-sensitive male-----------       GACAACCTTTGTATAACTTGACAACCTTCTCACGTCACTGCGCACACTTT
PAH-tolerant female----------       GACAACCTTTGTATAACTTGACAACCTTCTCA-GTCACTGCGCACACTTT
PAH-tolerant male------------       GACAACCTTTGTATAACTTGACAACCTTCTCA-GTCACTGCGCACACTTT
CYP1Arefseq------------------       GACAACCTTTGTATAACTTGACAACCCTCTCA-GTCACTGCGCACACTTT
                                    ************************** ***** *****************

PAH-sensitive female---------       TATGTTCACAAGTTAGGGCGTAAATGACTGCAGGCAAAACCTTTTTTCTT
PAH-sensitive male-----------       TATGTTCACAAGTTAGGGCGTAAATGACTGCAGGCAAAACCTTTTTTCTT
PAH-tolerant female----------       TATGTTCACAAGTTAGGGCGTAAATGACTGCAGGCAAAACCTTTTTTCTT
PAH-tolerant male------------       TATGTTCACAAGTTAGGGCGTAAATGACTGCAGGCAAAACCTTTTTTCTT
CYP1Arefseq------------------       TATGTTCACAAGTTAGGGCGTAAATGACTGCAGGCAAAACCTTTTTTCTT
                                    **************************************************

PAH-sensitive female---------       ACATATTTCTGTCCAATTCGGGTAAAACAGACACAGGGAATCTAGCACCG
PAH-sensitive male-----------       ACATATTTCTGTCCAATTCTGGTAAAATAGACACAGGGAATCTAGCACCG
PAH-tolerant female----------       ACATATTTCTGTCCAATTCTGGTAAAATAGACACAGGGAATCTAGCACCG
PAH-tolerant male------------       ACATATTTCTGTCCAATTCTGGTAAAATAGACACAGGGAATCTAGCACCG
CYP1Arefseq------------------       ACATATTTCTGTCCAATTCGGGTAAAACAGACACAGGGAATCTAGCACCT
                                    ******************* ******* ********************* 

PAH-sensitive female---------       TAAAAAAATGGTCGGGTCTCAGAATCCACTGCTTTTTTTCCCCTGCTTCA
PAH-sensitive male-----------       TAAAA---TGGTCGGGTCTCAGAGTCCACTGCTATTTTTCCC-TGCTGCA
PAH-tolerant female----------       TAAAA---TGGTCGGGTCTCAGAGTCCACTGCTATTTTTCCC-TGCTGCA
PAH-tolerant male------------       TAAAA---TGGTCGGGTCTCAGAGTCCACTGCTATTTTTCCC-TGCTGCA
CYP1Arefseq------------------       CAAAA---TGGTCGGGTCTCAGAGTCCACTGCTTTTTTTCCCCTGCTGCA
                                     ****   *************** ********* ******** **** **

PAH-sensitive female---------       CTCATGTCCACGCACACCTTGGATCTGCACAGTGGCACAGATAACTTTCT
PAH-sensitive male-----------       CTCATGTCCACACACACCTTGGATCTGCACAGTGGCACAGATAACTTTCT
PAH-tolerant female----------       CTCATGTCCACACACACCTTGGATCTGCACAGTGGCACAGATAACTTTCT
PAH-tolerant male------------       CTCATGTCCACACACACCTTGGATCTGCACAGTGGCACAGATAACTTTCT
CYP1Arefseq------------------       CTCATGTCCACGCACACCTTGGATCTGCACAGTGGCACAGATAACTTTCT
                                    *********** **************************************
```

```
PAH-sensitive female---------        GTTTCGTCTTTAATCTCTTAATCTTGTAAAGAATTCATGAACTACATCAC
PAH-sensitive male-----------        GTTTCGTCTTTAATCTCTTAATCTTGTAAAGAATTCATGAACTACATCAG
PAH-tolerant female----------        GTTTCGTCTTTAATCTCTTAATCTTGTAAAGAATTCATGAACTACATCAG
PAH-tolerant male------------        GTTTCGTCTTTAATCTCTTAATCTTGTAAAGAATTCATGAACTACATCAG
CYP1Arefseq------------------        GTTTCGTCTTTAATCTCTTAATCTTGTAAAGAATTCATGAACTACATCAC
                                     ************************************************* 

PAH-sensitive female---------        TCCCATCACTCGCATTCGACATTCTTCACTGCTCCACCAGTCACCAGACG
PAH-sensitive male-----------        TCCCATCACTCGCATTCGACATTCTTCACTGCTCCACCAGTCACCAGACG
PAH-tolerant female----------        TCCCATCACTCGCATTCGACATTCTTCACTGCTCCACCAGTCACCAGACG
PAH-tolerant male------------        TCCCATCACTCGCATTCGACATTCTTCACTGCTCCACCAGTCACCAGACG
CYP1Arefseq------------------        TCCCATCACTCGCATTCGACATTCTTCACTGCTCCACCAGTCACCAGACG
                                     **************************************************

PAH-sensitive female---------        AGACCTCCTACTTTTGGTCCAAGAAACACACTCTTATTTTCCAGAAAACA
PAH-sensitive male-----------        AGACCTCCTACTTTTGGTCCAAGAAACACACTCTTATTTTCCAGAAAACA
PAH-tolerant female----------        AGACCTCCTACTTTTGGTCCAAGAAACACACTCTTATTTTCCAGAAAACA
PAH-tolerant male------------        AGACCTCCTACTTTTGGTCCAAGAAACACACTCTTATTTTCCAGAAAACA
CYP1Arefseq------------------        AGACCTCCTACTTTTGGTCCAAGAAACACACTCTTATTTTCCAGAAAACA
                                     **************************************************

PAH-sensitive female---------        ACTCGTATTTATGTTGCAAATGACGCAGAACTGTACAATATGAGTTAAAC
PAH-sensitive male-----------        ACTCTTATTTATGTTGAAAATGACGCAGAACTGTACAATATGGGTTAAAC
PAH-tolerant female----------        ACTCTTATTTATGTTGAAAATGACGCAGAACTGTACAATATAGGTTAAAC
PAH-tolerant male------------        ACTCTTATTTATGTTGAAAATGACGCAGAACTGTACAATATAGGTTAAAC
CYP1Arefseq------------------        ACTCGTATTTATGTTGCAAATGACGCAGAACTGTACTATATGGGTTAAAC
                                     **** *********** ******************* ****  *******

PAH-sensitive female---------        CTAACTTTGTTAAAACATTACAAAAAAAGTTACTTTTTT-CAGTAATTCA
PAH-sensitive male-----------        CTAACTTTGTTAAAACATTACAAAAAAAGTAACTTTTTTTCAGTAATTCA
PAH-tolerant female----------        CTAACTTTGTTAAAACATTACAAAAAAAGTAACTTTTTTTCAGTAATTCA
PAH-tolerant male------------        CTAACTTTGTTAAAACATTACAAAAAAAGTAACTTTTTTTCAGTAATTTA
CYP1Arefseq------------------        CTAACTTTGTTAAAACATTACAAAAAAAGTGACTTTTTT-CAGTAATTCA
                                     ****************************** ******** ******** *

PAH-sensitive female---------        AATCAACATTTGTACTCCACACAGAGTTTCATTCTGGCACGTTTCAATTA
PAH-sensitive male-----------        AATCAACATTTGTACTCCACACAGAGTTTGATTCTGGCACGTTTCAATTA
PAH-tolerant female----------        AATCAACATTTGTACTCCACACAGAGTTTGATTCTGGCACGTTTCAATTA
PAH-tolerant male------------        AATCAACATTTGTACTCCACACAGAGTTTGATTCTGGCACGTTTCAATTA
CYP1Arefseq------------------        AATCAACATTTGTACTCCACACAGAGTTTCATCCTGGCACTTTTCAATTA
                                     ***************************** ** ******* *********
```

```
PAH-sensitive female---------        ATTTGGGTGCCCTACAGCCAGTAACACCCCTTCCCCATTACTGCATCATT
PAH-sensitive male-----------        ATTTGGGTGCCCTACAGCCAGTAACACCCCTTCCCCATTACTGCATCATC
PAH-tolerant female----------        ATTTGGGTGCCCTACAGCCAATAACACCACTTCCCCAT--------CATT
PAH-tolerant male------------        ATTTGGGTGCCCTACAGCCAATAACACCACTTCCCCAT--------CATT
CYP1Arefseq------------------        ATTTGGGTGCCCTACAGCCAGTAACACCCCTTCCCCATTACTGCATCATT
                                     ******************** ******* *********        *** 

PAH-sensitive female---------        GCGGTGTGGTATGTAGACAGCCTGTGGCTCGATGAGGTGATGAGGAAGCC
PAH-sensitive male-----------        GCAGTGTGGTATGTAGACAGCCTGTGGCTCGATGAGGTGATGAAGAAGCC
PAH-tolerant female----------        GCAGTGTGGTGTGTAGACAGCCTGTGGCTCAATGAGGTGATGAGGAAGCC
PAH-tolerant male------------        GCAGTGTGGTATGTAGACAGCCTGTGGCTCAATGAGGTGATGAGGAAGCC
CYP1Arefseq------------------        GCGGTGTGGTATGTAGACAGCCTGTGGCTCGATGAGGTGATGAGGAAGCC
                                     ** ******* ******************* ************ ******

PAH-sensitive female---------        CAGGTTAAAAGAGACCCTCAGCTGATCTGCACTGTTGGGTCTGACTCTGG
PAH-sensitive male-----------        CAGGTTAAAAGAGACCCTCAGCTGATCTGCACTGTGGTGTCTG--TCTGG
PAH-tolerant female----------        CAGGTTAAAAGAGACCCTCAGCTGATCTGCACTGTTGGGTCTGACTCTGG
PAH-tolerant male------------        CAGGTTAAAAGAGACCCTCAGCTGATCTGCACTGTTGGGTCTGACTCTGG
CYP1Arefseq------------------        CAGGTTAAAAGAGACCCTCAGCTGATCTGCACTGTTGTGTCTGACTCTGG
                                     *********************************** * *****  *****

PAH-sensitive female---------        TGTTTCTTGTCTTGCCATTGACAATAACCTACAGTCCCTCAATGGGGTCT
PAH-sensitive male-----------        TGTTTCTTGTCTTGCATTTGACAATAACCTACAGTCCCTCAATGGGGTCT
PAH-tolerant female----------        TGTTTCTTGTCTTGCCATTGACAATAACCTACAGTCCCTCAATGGGGTGT
PAH-tolerant male------------        TGTTTCTTGTCTTGCCATTGACAATAACCTACAGTCCCTCAATGGGGTGT
CYP1Arefseq------------------        TGTTTCTTGTCTTGCCATTGACAATAACCTACAGTCCCTCAATGGGGTCT
                                     ***************  ******************************* *

PAH-sensitive female---------        AGATCATACAAGTTTGCTGTCAAATCAAGCACAGTAATACCATGGCCACT
PAH-sensitive male-----------        AGATCATAAAAGTTTGCTGTCAA-TCGAGCACAGTAATACCATGGCCACT
PAH-tolerant female----------        AGATCATACAAGTTTGCTGTCAAATCAAACACAGTAATACCATGGCCACT
PAH-tolerant male------------        AGATCATACAAGTTTGCTGTCAAATCAAACACAGTAATACCATGGCCACT
CYP1Arefseq------------------        AGATCATACAAGTTTGCTGTCAAATCAAGCACAGTAATACCATGGCCACT
                                     ******** ************** ** * *********************

PAH-sensitive female---------        GAACCACAAGTTGGTACGT-TTGGCCAAGTGGCCCAATCATCCTGAGAAA
PAH-sensitive male-----------        GAACCACAAGTTGGTATGCCTTGGCCTAGTGGCCCAATCATCCTGAGAAA
PAH-tolerant female----------        GAACCACAAGTTGGTACGT-TTGGCCTAGTGGCCCAATCATCCTGAGAAA
PAH-tolerant male------------        GAACCACAAGTTGGTACGT-TTGGCCTAGTGGCCCAATCATCCTGAGAAA
CYP1Arefseq------------------        GAACCACAAGTTGGTACGT-TTGGCCAAGTGGCCCAATCATCCTGAGAAA
                                     **************** *  ****** ***********************
```

```
PAH-sensitive female---------      TGAAGTCCTAAAAGCTTGTCAGCATCGGGAATCATGAAATTCTTAGGAAG
PAH-sensitive male-----------      TGAAGTCCAAAAAGCTTGTCAGCATCGGGAATCATGAAATTCTTAGGAAG
PAH-tolerant female----------      TGAAGTCCAAAAAGCTTGTCAGCATCGGGAATCATGAAATTCTTAGGAAG
PAH-tolerant male------------      TGAAGTCCAAAAAGCTTGTCAGCATCGGGAATCATGAAATTCTTAGGAAG
CYP1Arefseq------------------      TGAAGTCCTAAAAGCTTGTCAGCATCGGGAATCATGAAATTCTTAGGAAG
                                   ******** *****************************************

PAH-sensitive female---------      ACAGCTAACTTATTCTTTCAGATGTTTTCCTTCCACTCAACTTTCCATCA
PAH-sensitive male-----------      ACGGCTAACTTATTCTTTCAGATGTTTTCCTTCCACTCAACTTTCCATCA
PAH-tolerant female----------      ACGGCTAACTTATTCTTTCAGATGTTTTCCTTCCACTCAACTTTCCATCA
PAH-tolerant male------------      ACGGCTAACTTATTCTTTCAGATGTTTTCCTTCCACTCAACTTTCCATCA
CYP1Arefseq------------------      ACAGCTAACTTATTCTTTCAGATGTTTTCCTTCCACTCAACTTTCCATCA
                                   ** ***********************************************

PAH-sensitive female---------      ATACGGGTGGAGACAGCACTCTGTAAGCAGCTTCTTCAGCAGAAGTACTC
PAH-sensitive male-----------      ATACGGGTGGAGACAGCACTCTGTAAGCAGCTACTTTAGCAGAAGTTCTC
PAH-tolerant female----------      ATACGGGTGGAGACAGCACTCTGTAAGCAGCTACTTTAGCAGAAGTTCTC
PAH-tolerant male------------      ATACGGGTGGAGACAGCACTCTGTAAGCAGCTACTTTAGCAGAAGTTCTC
CYP1Arefseq------------------      ATACGGGTGGAGACAGCACTCTGTAAGCAGCTTCTTCAGCAGAAGT----
                                   ******************************** *** *********

PAH-sensitive female---------      AGTGTCAAGTCAGCAAGCTCACCCACGGTTGTGTAGGCCACAAACCTGCA
PAH-sensitive male-----------      CATGTCAAGTCAGCAAGCTCACCCATGGTTGTGTAGGCCACAAATCTGAA
PAH-tolerant female----------      CATGTCAAGTCAGCAAGCTCACCCATGGTTGTGTAGGCCACAAATCTGAA
PAH-tolerant male------------      CATGTCAAGTCAGCAAGCTCACCCATGGTTGTGTAGGCCACAAATCTGAA
PAH-sensitive female---------      AGTGTCAAGTCAGCAAGCTCACCCACGGTTGTGTAGGCCACAAACCTGCA
CYP1Arefseq------------------      ----------CAGCAAGCTCACCCACGGTTGTGTAGGCCACAAACCTGCA
                                             *************** ****************** *** *

PAH-sensitive female---------      TTTCTGTATGAAACATAATTTATGTATTGATCTTATGAAACTGAAATGTT
PAH-sensitive male-----------      TTTCTGTATGAAAAATAATTTATGTATTGATCTTATGAAACTGAAATTTT
PAH-tolerant female----------      TTTCTGTATGAAAAATAATTTATGTATTGATCTTATGAAACTGAAATTTT
PAH-tolerant male------------      TTTCTGTATGAAAAATAATTTATGTATTGATCTTATGAAACTGAAATTTT
CYP1Arefseq------------------      TTTCTGTATGAAACATAATTTATGTATTGATCTTATGAAACTGAAATGTT
                                   ************* ********************************* **

PAH-sensitive female---------      ATTTGCATTAGTTGTAAGCAATTATACATTTGAGGTTTTCTGCTTTCTTA
PAH-sensitive male-----------      ATTTGCATTAGTTGTAAGCAATTATACATTTGAGGTTTTCTGCTTTCTTA
PAH-tolerant female----------      ATTTGCATTAGTTGTAAGCAATTATACATTTGAGGTTTTCTGCTTTCTTA
PAH-tolerant male------------      ATTTGCATTAGTTGTAAGCAATTATACATTTGAGGTTTTCTGCTTTCTTA
CYP1Arefseq------------------      ATTTGCATTAGTTGTAAGCAATTATACATTTGAGGTTTTCTGCTTTCTTA
                                   **************************************************
```

```
PAH-sensitive female---------        CTTTATTAGTCCACATTGATTAACTCTTGGCCAGTTATTTGTTTCAAATT
PAH-sensitive male-----------        CTTTATTAGTCCACATTGATTAACTCTTGGCCAGTTATTTGTTTCAAATT
PAH-tolerant female----------        CTTTATTAGTCCACATTGATTAACTCTTGGCCAGTTATTTGTTTCAAATT
PAH-tolerant male------------        CTTTATTAGTCCACATTGATTAACTCTTGGCCAGTTATTTGTTTCAAATT
CYP1Arefseq------------------        CTTTATTAGTCCAAATTGATTAACTCTTGGTCAGTTATTTGTTTCAAATT
                                     ************* **************** *******************

PAH-sensitive female---------        CATTTTCACTCTTGTTATGTGATGCCTAAAATTGACAAATATTCTTATTG
PAH-sensitive male-----------        CATTTTCACTCTTGTTATGTGATGCTTAAAATTGACAAATATTCTTATTG
PAH-tolerant female----------        CATTTTCACTCTTGTTATGTGATGCTTAAAATTGACAAATATTCTTATTG
PAH-tolerant male------------        CATTTTCACTCTTGTTATGTGATGCTTAAAATTGACAAATATTCTTATTG
CYP1Arefseq------------------        CATTTTCACTCTTGTTATGTGATGCCTAAAATTGACAAATATTCTTATTG
                                     ************************* ************************

PAH-sensitive female---------        TGATATTAATACACAAGTACATCTCCAAAAAAATTATATTGCGTAAACAT
PAH-sensitive male-----------        TGATAGTAATGTGCAAGTACATCTCCAAAAAATTCATGTTGCGTAAACAT
PAH-tolerant female----------        TGATAGTAATGTGCAAGTACATCTCCAAAAAATTCATGTTGCGTAAACAT
PAH-tolerant male------------        TGATAGTAATGTGCAAGTACATCTCCAAAAAATTCATGTTGCGTAAACAT
CYP1Arefseq------------------        TGATATTAATACACAAGTACATCTCCAAAAAAATTATATTGCGTAAACAT
                                     ***** ****   ******************* * ** ************

PAH-sensitive female---------        TTTTTGCTGGTCATTTCAGAAAAGGAAATTATTATAATTATTTTATAGAT
PAH-sensitive male-----------        TTTTTGCTGGTCATTTCAGAAAAGGAAATTATTATAATTATTTTATAAAT
PAH-tolerant female----------        TTTTTGCTGGTCATTTCAGAAAAGGAAATTATTATAATTATTTTATAAAT
PAH-tolerant male------------        TTTTTGCTGGTCATTTCAGAAAAGGAAATTATTATAATTATTTTATAAAT
CYP1Arefseq------------------        TTTTTGCTGGTCATTTCAGAAAAGGAAATTATTATAATTATTTTATAGAT
                                     *********************************************** **

PAH-sensitive female---------        ATTCATTACACATAGAGTGAAAATAATATTTTAGTCTTTTATTTCTTGTA
PAH-sensitive male-----------        AGTCATTACACATAGAGTGAAAATAATATTTTAGCCTTTTATTTCTTGTA
PAH-tolerant female----------        AGTCATTACACATAGAGTGAAAATAATATTTTAGCCTTTTATTTCTTGTA
PAH-tolerant male------------        AGTCATTACACATAGAGTGAAAATAATATTTTAGCCTTTTATTTCTTGTA
CYP1Arefseq------------------        ATTCATTACACATAGAGTGAAAATAATATTTTAGTCTTTTATTTCTTGTA
                                     * ******************************** ***************

PAH-sensitive female---------        TTTTTGATGATTACAACAAATAAATAAGTTCAGTGTCTCAGAAAAGGATG
PAH-sensitive male-----------        TTTTTGATGATTACAACAAATAAATAAGTTCAGTGTCTCAGAAAAGGATG
PAH-tolerant female----------        TTTTTGATGATTACAACAAATAAATAAGTTCAGTGTCTCAGAAAAGGATG
PAH-tolerant male------------        TTTTTGATGATTACAACAAATAAATAAGTTCAGTGTCTCAGAAAAGGATG
CYP1Arefseq------------------        TTTTTGATGATTACAACAAATAAATAAGTTCAGTGTCTCAGAAAAGGATG
                                     **************************************************
```

```
PAH-sensitive female---------       TTTTGAGAACAAATGTCCAGCATGTGAAAAGTATGTCCATTTCTATGCAC
PAH-sensitive male-----------       TTTTAAGCACAAATGTCCACCATCTGAAAAGTGGGTCCATTTCTATGCAC
PAH-tolerant female----------       TTTTAAGCACAAATGTCCACCATCTGAAAAGTGGGTCCATTTCTATGCAC
PAH-tolerant male------------       TTTTAAGCACAAATGTCCACCATCTGAAAAGTGGGTCCATTTCTATGCAC
CYP1Arefseq------------------       TTTTGAGAACAAATGTCCAGCATGTGAAAAGTATGTCCATTTCTATGCAC
                                    **** ** *********** *** ********  ****************

PAH-sensitive female---------       CCAACACTTGGTTGGGCCTCCTCTTGCGTGAATTACTGCATTGATGTGCC
PAH-sensitive male-----------       ACAATACTTGGTTGGGCCTCCTCTTGCATCAATTACTGCATCGATGTGCC
PAH-tolerant female----------       ACAATACTTGGTTGGGCCTCCTCTTGCATCAATTACTGCATCGATGTGCC
PAH-tolerant male------------       ACAATACTTGGTTGGGCCTCCTCTTGCATCAATTACTGCATCGATGTGCC
CYP1Arefseq------------------       CCAACACTTGGTTGGGCCTCCTCTTGCGTGAATTACTGCATTGATGTGCC
                                     *** ********************* * *********** ********

PAH-sensitive female---------       GAAATGATCTGCCTGTGGTGCTGCATAGATGTAATGGAAGCCCAGGTTGC
PAH-sensitive male-----------       GAAGTGATCTGCCTGTGGTGCTGCAT----GTAATGGAAGCCCAGATTGC
PAH-tolerant female----------       GAAGTGATCTGCCTGTGGTGCTGCAT----GTAATGGAAGCCCAGATTGC
PAH-tolerant male------------       GAAGTGATCTGCCTGTGGTGCTGCAT----GTAATGGAAGCCCAGATTGC
CYP1Arefseq------------------       GAAGTGATCTGCCTGTGGTGCTGCATAGATGTAATGGAAGCCCAGGTTGC
                                    *** *********************    *************** ****

PAH-sensitive female---------       TTTAATAAGATCCTTCTGGGTCATCTGCATTGCTGGATCTGGTGTCCTCA
PAH-sensitive male-----------       TTTGATAAGATCCTT-TGGGTCATCTGCATTGCTGGATTTGGTGTCCTCA
PAH-tolerant female----------       TTTGATAAGATCCTT-TGGGTCATCTGCATTGCTGGATTTGGTGTCCTCA
PAH-tolerant male------------       TTTGATAAGATCCTT-TGGGTCATCTGCATTGCTGGATTTGGTGTCCTCA
CYP1Arefseq------------------       TTTAATAAGATCCTTCTGGGTCATCTGCATTGCTGGATCTGGTGTCCTCA
                                    *** *********** ********************* ***********

PAH-sensitive female---------       ACTTCTTCTTGACAACAGACAATAGATTTTCTATGGGATTCAGGTCAGGG
PAH-sensitive male-----------       ACTTCTTCTTGACAACAGCCAATAGATTTTCTATGGGATTCAGGTCAGGG
PAH-tolerant female----------       TCTTCTTCTTGACAACAGCCAATAGATTTTCTATGGGATTCAGGTCAGGG
PAH-tolerant male------------       TCTTCTTCTTGACAACAGCCAATAGATTTTCTATGGGATTCAGGTCAGGG
CYP1Arefseq------------------       ACTTCTTCTTGACAACAGACAATAGATTTTCTATGGGATTCAGGTCAGGG
                                     ***************** *******************************

PAH-sensitive female---------       GAGTTGGCTGGCCAATCAAGAACAGGAACACCACTTTCATTGAACCATTC
PAH-sensitive male-----------       GAGTTTGCTGGCCAATCAAGAACAGGAACACCACTTTCATTGATCCATTC
PAH-tolerant female----------       GAGTTTGCTGGCCAATCAAGAACAGGGACACCACTTTCATTGATCCATTC
PAH-tolerant male------------       GAGTTTGCTGGCCAATCAAGAACAGGAACACCACTTTCATTGAACCATTC
CYP1Arefseq------------------       GAGTTGGCTGGCCAATCAAGAACAGGAACACCACTTTCATTGAACCATTC
                                    ***** ******************** **************** ******
```

```
PAH-sensitive female---------        ATTGGTACCTTTGGCAGTGTGGGCAGGTGCCGTGTCCTGCTGGAAAATAA
PAH-sensitive male-----------        ATTGGTACCTTTGGTAGTGTGGGCAGGTGCCGAGTCCTGCTGGAAAATTA
PAH-tolerant female----------        ATTCGTACCTTTGGCAGTGTGGGCAGGTGCCGAGTCCTGCTGGAAAATTA
PAH-tolerant male------------        ATTCGTACCTTTGGCAGTGTGGGCAGGTGCCGAGTCCTGCTGGAAAATTA
CYP1Arefseq------------------        ATTGGTACCTTTGGCAGTGTGGGCAGGTGCCGTGTCCTGCTGGAAAATAA
                                     *** ********** ***************** *************** *

PAH-sensitive female---------        AATTAGCATCTCCATACAGCTTGTCTGCAGAAGGAAACATGAAGTGCTCT
PAH-sensitive male-----------        AATTAGCATCTCCATACAGCTTGTCTGCAGAAGGAAACATGAAGTGCTCT
PAH-tolerant female----------        AATTAGCATCTCCATACAGCTTGTCTGCAGAAGGAAACATGAAGTGCTCT
PAH-tolerant male------------        AATTAGCATCTCCATACAGCTTGTCTGCAGAAGGAAACATGAAGTGCTCT
CYP1Arefseq------------------        AATTAGCATCTCCATACAGCTTGTCTGCAGAAGGAAACATGAAGTGCTCT
                                     **************************************************

PAH-sensitive female---------        ATAATGTCT---TAGTAGCAGCATTGACTGTGGAAATACAGTGGACCAAC
PAH-sensitive male-----------        AAAAAGGCAGAATAGCAGGAGCAGTGACTGTGGAAATACTGTGGACCAAC
PAH-tolerant female----------        ATAATTTCT---TAGTAGGAGCATTGACTGTGGAAATACAGTGGACCAAC
PAH-tolerant male------------        ATAATTTCT---TAGTAGGAGCATTGACTGTGGAAATACAGTGGACCAAC
CYP1Arefseq------------------        ATAATGTCT---TAGTAGCAGCATTGACTGTGGAAATACAGTGGACCAAC
                                     * **   *    *** ** **** *************** **********

PAH-sensitive female---------        ACCAGCAGATTGCATGGCTCTCTAAACCATCACTGACTGTGGAAACTGCA
PAH-sensitive male-----------        ACCAGCAGATTGCATGGCTCTCTAAACCATCACTGACTGTGGGAACTGCA
PAH-tolerant female----------        ACCAGCAGATTGCATGGCTCTCCAAACCATCACTGACTGTGGAAACTGCA
PAH-tolerant male------------        ACCAGCAGATTGCATGGCTCTCCAAACCATCACTGACTGTGGAAACTGCA
CYP1Arefseq------------------        ACCAGCAGATTGCATGGCTCTCTAAACCATCACTGACTGTGGAAACTGCA
                                     ********************** ******************* *******

PAH-sensitive female---------        CTTCAATTAACATGGATTCTGGGCCTCTCCACTCTTCCTCTCCACTCTTC
PAH-sensitive male-----------        CTTCAATTAACATGGATTCTGGACCTCTCCACTCTTCCTCTCCACTCTTC
PAH-tolerant female----------        CTTCAATTAACATGGATTCTGGACCTCTCCACTCTTCCTCTCCACTCTTC
PAH-tolerant male------------        CTTCAATTAACATGGATTCTGGACCTCTCCACTCTTCCTCTCCACTCTTC
CYP1Arefseq------------------        CTTCAATTAACATGGATTCTGGGCCTCTCCACTCTTCCTCTCCACTCTTC
                                     ********************** ***************************

PAH-sensitive female---------        CTCCAGAATCTGGGGCCTTGATTTCCAAATAAAATGAAGAATTCAATTTC
PAH-sensitive male-----------        CTCCAGAATCTGGGGCCTTGATTTCCAAATAAAATGAAGAATTCAATTTC
PAH-tolerant female----------        CTCCAGAATCTGGGGCCTTGATTTCCAAATAAAATGAAGAATTCAATTTC
PAH-tolerant male------------        CTCCAGAATCTGGGGCCTTGATTTCCAAATAAAATGAAGAATTCAATTTC
CYP1Arefseq------------------        CTCCAGAATCTGGGGCCTTGATTTCCAAATAAAATGAAGAATTCAATTTC
                                     **************************************************
```

```
PAH-sensitive female---------       ATCTGAAAAAA-CATCTTTGGATCACTGAGCAACAGTCCAGATCCTTTTC
PAH-sensitive male-----------       ATCTGAAAAAAACATCTTTGGATCACTGAGCAACAGTCCAGATCCTTTTC
PAH-tolerant female----------       ATCTGAAAAAAACATCTTTGGATCACTGAGCAACAGTCCAGATCCTTTTC
PAH-tolerant male------------       ATCTGAAAAAAACATCTTTGGATCACTGAGCAACAGTCCAGATCCTTTTC
CYP1Arefseq------------------       ATCTGAAAAAA-CATCTTTGGATCACTGAGCAACAGTCCAGATCCTTTTC
                                    *********** **************************************

PAH-sensitive female---------       TCCACAGACCAGGTAAGACGTTATCTGTGCAGGACCCGTCTGTGTGGTGG
PAH-sensitive male-----------       TCCATAGACCAGGTAAGACGTTACCTGTGCAGGACCTGTCTGTGTGGTGG
PAH-tolerant female----------       TCCATAGACCAGGTAAGACGTTACCTGTGCAGGACCTGTCTGTGTGGTGG
PAH-tolerant male------------       TCCATAGACCAGGTAAGACGTTACCTGTGCAGGACCTGTCTGTGTGGTGG
CYP1Arefseq------------------       TCCACAGACCAGGTAAGACGTTATCTGTGCAGGACCCGTCTGTGTGGTGG
                                    **** ****************** ************ *************

PAH-sensitive female---------       CTCTCGATGCACCGACTCCAGCCTCAGTCCACTTCTTAAGAAGCTCCCAC
PAH-sensitive male-----------       CTCTCCATGCACCGACTCCAGCCTCAGTCCACTCATTAAGAAGCTCCCAC
PAH-tolerant female----------       CTCTCGATGCACCGACTCCAGCCTCAGTCCACTCCTTAAGAAGCTCCCAC
PAH-tolerant male------------       CTCTCGATGCACCGACTCCAGCCTCAGTCCACTCCTTAAGAAGCTCCCAC
CYP1Arefseq------------------       CTCTCGATGCACCGACTCCAGCCTCAGTCCACTTCTTAAGAAGCTCCCAC
                                    ***** ***************************  ***************

PAH-sensitive female---------       AGATTCTTAAATAGATTTTGGTTGGTAATCCTGTCAAGGCTGCAGTTCTC
PAH-sensitive male-----------       AGATTCTTAAATAGATTTCGGTTGGCAATCCTCTCAAGGCTGCAGTTCTC
PAH-tolerant female----------       AGATTCTTAAATAGATTTCGGTTGGCAATCCTCTCAAGGCTGCAGTTCTC
PAH-tolerant male------------       AGATTCTTAAATAGATTTCGGTTGGCAATCCTCTCAAGGCTGCAGTTCTC
CYP1Arefseq------------------       AGATTCTTAAATAGATTTTGGTTGGTAATCCTGTCAAGGCTGCAGTTCTC
                                    ****************** ****** ****** *****************

PAH-sensitive female---------       TCTGTTGCTGGGGCACCTTTCCTACCGCATTTCTTCCTTCCACTCAATTT
PAH-sensitive male-----------       CCTGTTGCTGGGGCACCTTTCCTACCACATTTCTTCCTTCCACTCAATTT
PAH-tolerant female----------       CCTGTTGCTGGGGCACCTTTCCTACCACATTTCTTCCTTCCACTCAATTT
PAH-tolerant male------------       CCTGTTGCTGGGGCACCTTTCCTACCACATTTCCTCCTTCCACTCAATTT
CYP1Arefseq------------------       TCTGTTGCTGGGGCACCTTTCCTACCGCATTTCTTCCTTCCACTCAATTT
                                     ************************* ****** ****************

PAH-sensitive female---------       TCTGTTAATATGTTTGGATATAGAATTCTGTGAACAACCAGCTTCTTTCA
PAH-sensitive male-----------       TCTGTTAATATGTTTGGATATAGAATTCTGTGAACAACCAGCTTCTTTCA
PAH-tolerant female----------       TCTGTTAATATGTTTGGATATAGAATTCTGTGAACAACCAGCTTCTTTCA
PAH-tolerant male------------       TCTGTTAATATGTTTGGATATAGAATTCTGTGAACAACCAGCTTCTTTCA
CYP1Arefseq------------------       TCTGTTAATATGTTTGGATATAGAATTCTGTGAACAACCAGCTTCTTTCA
                                    **************************************************
```

```
PAH-sensitive female---------       GGAATGACCTTTCGTGGCTTTTCCTGCTTGTGAAGGATGTCGATGACTGT
PAH-sensitive male-----------       GGAATGACCTTTCGTGGCTTTTCCTGCTTGTGAAGGATGTCGATGA----
PAH-tolerant female----------       GGAATGACCTTTCGTGGCTTTTCCTGCTTGTGAAGGATGTCGATGACTGT
PAH-tolerant male------------       GGAATGACCTTTCGTGGCTTTTCCTGCTTGTGAAGGATGTCGATGACTGT
CYP1Arefseq------------------       GGAATTACCTTTCGTGGCTTTTCCTGCTTGTGAAGGATGTCGATGACTGT
                                    ***** ****************************************

PAH-sensitive female---------       CTTCAAGACATCTGTCACGTCAGCCGTCTTGTCCATGATTGTGTCGCATA
PAH-sensitive male-----------       -----------------------CCGTCTTGTCCATGATTGTGTCGCATA
PAH-tolerant female----------       CTTC--------TGTCACGTCAGCCGTCTTCCCCATGATTGTGTCGCATA
PAH-tolerant male------------       CTTC--------TGTCACGTCAGCCGTCTTCCCCATGATTGTGTCGCATA
CYP1Arefseq------------------       CTTC--------TGTCACGTCAGCCGTCTTCCCCATGATTGTGTCGCATA
                                                           *******  ******************

PAH-sensitive female---------       TTGCCCCAGAGCGAGAGACCGTCTGAGGGCTCAGGAAATATTTGCAGGTA
PAH-sensitive male-----------       TTGCCCCAGAGCGAGAGACCGTCTGAGGGCTCAGGAAATATTTACAGGTA
PAH-tolerant female----------       CTGCCCCAGAGCGAGAGACCGTCTGAGGGTTCAGGAAATATTTGCAGGTA
PAH-tolerant male------------       CTGCCCCAGAGCGAGAGACCGTCTGAGGGTTCAGGAAATATTTGCAGGTA
CYP1Arefseq------------------       CTGCCCCAGAGCGAGAGACCGTCTGAGGGCTCAGGAAATATTTGCAGGTA
                                     **************************** ************* ******

PAH-sensitive female---------       TTTTGTGTTAACTAGCTGATTAGGGTGTGACACTAAGATTTTCCAATAAT
PAH-sensitive male-----------       TTTTGTGTTAATTAGCTGATTAGGGTGTGACACTAAGATTTTCCAATAAT
PAH-tolerant female----------       TTTTGCGTTAATTAGCTGAATAGGGTGTGACACTAAGATTTTCTAATCAT
PAH-tolerant male------------       TTTTGCGTTAATTAGCTGAATAGGGTGTGACACTAAGATTTTCTAATCAT
CYP1Arefseq------------------       TTTTGCGCTAATTAGCTGAATAGGGTGTGACACTAAGATTTTCTAATCAT
                                    ***** * *** ******* *********************** *** **

PAH-sensitive female---------       GAACATTTTCACAGAATTCTAATTTTTTGGGACACTGATTTTTTCTTTAT
PAH-sensitive male-----------       GAACATTTTCACAGAATTCTAATTTTTTGGGACACTGATTTTTTCTTTAT
PAH-tolerant female----------       GAACATTTTCACAGAATTCTAATTTTCTGGGACACTGATTTTTTCTTTAT
PAH-tolerant male------------       GAACATTTTCACAGAATTCTAATTTTCTGGGACACTGATTTTTTCTTTAT
CYP1Arefseq------------------       GAACATTTTCACAGAATTCTAATTTTCTGGGACACTGATTTTTTCTTTAT
                                    ************************** ***********************

PAH-sensitive female---------       TATTTTTATCTGTAAGCCATAATTATCCAAATTAAAAGAAACAAACGTTT
PAH-sensitive male-----------       TATTTTTATCTGTAAGCCATAATTATCCAAATTAAAAGAAACAAACGTTT
PAH-tolerant female----------       TATTTTTATCTGTAAGCCATAATTATCCAAATTAAAAGAAACAAACGTTT
PAH-tolerant male------------       TATTTTTATCTGTAAGCCATAATTATCCAAATTAAAAGAAACAAACGTTT
CYP1Arefseq------------------       TATTTTTATCTGTAAGCCATAATTATCCAAATTAAAAGAAACAAACGTTT
                                    **************************************************
```

```
PAH-sensitive female---------         AGGATAATATAATATATGAGT-----------------------------
PAH-sensitive male-----------         AGGATAATATAATATATGAGT-----------------------------
PAH-tolerant female----------         AGGATAATATAATATATGAGT-----------------------------
PAH-tolerant male------------         AGGATAATATAATATATGAGT-----------------------------
CYP1Arefseq------------------         AGGATCATATAATATATGAGTTTCACTTTCTGAGATGACTGGCAAAAAAT
                                      ***** ***************

PAH-sensitive female---------         ----------------------------ATTGAGATATACCTGTACTCAT
PAH-sensitive male-----------         ----------------------------ATTGAGATATACCTGTACTCAT
PAH-tolerant female----------         ----------------------------ATTGAGATATACCTGTACTCAT
PAH-tolerant male------------         ----------------------------ATTGAGATATACCTGTACTCAT
CYP1Arefseq------------------         ATTGCACTTTTAGAGAATATACAAACGTATTGAGATACACCTGTACTCAT
                                                                  ********* ************

PAH-sensitive female---------         TAGCTTTCTTTTTTGTTGTTGTT---TCCCTGTAAAGCACTGCACTTTAA
PAH-sensitive male-----------         TAGCTTTCTTTTTTGTTGTT------TCCCTGTAAAGCACTGCACTTTAA
PAH-tolerant female----------         TAGCTTTCTTTTTTGTTGTTGTT---TCCCTGTAAAGCACTGCACTTTAA
PAH-tolerant male------------         TAGCTTTCTTTTTTGTTGTTGTT---TCCCTGTAAAGCACTGCACTTTAA
CYP1Arefseq------------------         TAGCTTTCTTTTTTGTTGTTGTTGTTTCTCTGTAAAGCACTGCACTTTAA
                                      ********************      ** *********************

PAH-sensitive female---------         ATTACCTTGCATA----GGGTGCTTTAACAATAAACTTTCCTTGTGAGAC
PAH-sensitive male-----------         ATTACCTTGCATA----GGGTGCTTCAACAATAAACTTTCCTTGTGAGAC
PAH-tolerant female----------         ATTACCTTGCTTA----GGGTGCTTTAACAATAAACTTTCCTTGTGAGAC
PAH-tolerant male------------         ATTACCTTGCTTA----GGGTGCTTTAACAATAAACTTTCCTTGTGAGAC
CYP1Arefseq------------------         ATTACCTTGCATATAAAGGGTGCTTTAACAATAAACTTTCCTTGTGAGAC
                                      ********** **    ******** ************************

PAH-sensitive female---------         AAAAAAA-TATTGACAGAAATTATTAAAAATGCTGGAAAGAGAGGAAGTT
PAH-sensitive male-----------         AAAAAAAATATTGACAGAAATTATTAAAAATGCTGGAAAGAGTGGAAGTT
PAH-tolerant female----------         AAAAAAA-TATTGACAGAAATTATTAAAAATGCTGGAAAGAGAGGAAGTT
PAH-tolerant male------------         AAAAAAA-TATTGACAGAAATTATTAAAAATGCTGGAAAGAGAGGAAGTT
CYP1Arefseq------------------         AAAAAAC-TATTGACAGACATTATTAAAAGTGCTGGAAAGAGTGGAAGTT
                                      ******  ********** ********** ************ *******

PAH-sensitive female---------         ACAAATATTTTACCTCTCAAAATGATGTTATATTTTGTTCTTATGTGAAA
PAH-sensitive male-----------         ACAAATATTTTACCTCTCAAAATGATGTTATATTTTGTTCTTATGTGAAA
PAH-tolerant female----------         ACAAATATTTTACCTCTCAAAATGATGTTATATTTTGTTCTTATGTGAAA
PAH-tolerant male------------         ACAAATATTTTACCTCTCAAAATGATGTTATATTTTGTTCTTATGTGAAA
CYP1Arefseq------------------         ACAAATATTTTACCCCTCAAAATGATGTTATATTTTGTTCTTATGTGAAA
                                      ************** ***********************************
```

```
PAH-sensitive female---------        AACGTACAATGATATAAAAGTGGAGAATTATGATTTATTGGTGACCCTGC
PAH-sensitive male-----------        AACGTACAATGATATAAAACTGGAGAATTATGAATTATTGGTGACCCTGC
PAH-tolerant female----------        AACGTACAATGATATAAAAGTGGAGAATTATGATTTATTGGTGACCCTGC
PAH-tolerant male------------        AACGTACAATGATATAAAAGTGGAGAATTATGATTTATTGGTGACCCTGC
CYP1Arefseq------------------        AACGTACAATGATATAAAACTGAAGAATAATGATTTATTGGTGACCCTGC
                                     ******************* ** ***** **** ****************

PAH-sensitive female---------        TCATCCATTCCAGCAGGCTCATCTCTGAGACCCTGAAACGGGGCAGGAGG
PAH-sensitive male-----------        TCATCCATTCCAGCAGGCTCATCTCTGAGACCCTGAAACGGGGCAGGAGG
PAH-tolerant female----------        TCATCCATTCCAGCAGGCTCATCTTTGAGACCCTGAAACGGGGCAGGAGG
PAH-tolerant male------------        TCATCCATTCCAGCAGGCTCATCTTTGAGACCCTGAAACGGGGCAGGAGG
CYP1Arefseq------------------        TCATCCATTCCAGCAGGCTCATCTCTGAGACCCTGAAACGGGGCAGGAGG
                                     ************************ *************************

PAH-sensitive female---------        GCAGACGAGGTCATCTTATTCAAAGTTATCGTGGTATTTGCCCTACAGCG
PAH-sensitive male-----------        GCAGACGAGGTCATCTTATTCAAAGTTATCGTGGTATTTGCCCTACAGCG
PAH-tolerant female----------        GCAGACGAGGTCATCTTATTCAAAGTTATCGTGGTATTTGCCCTACAGCG
PAH-tolerant male------------        GCAGACGAGGTCATCTTATTCAAAGTTATCGTGGTATTTGCCCTACAGCG
CYP1Arefseq------------------        GCAGACGAGGTCATCTTATTCAAAGTTATCGTGGTATTTGCCCTACAGCG
                                     **************************************************

PAH-sensitive female---------        ACCTCAGCGGGTTGCATGACTGCAGCGAGTGAAACCTGAGTGGCAGATTG
PAH-sensitive male-----------        ACCTCAGCGGGTTGCATGACTGCAGCGAGTGAAACCTGAGTGGCAGATTG
PAH-tolerant female----------        ACCTCAGCGGGTTGCATGACTGCAGCGAGTGAAACCTGAGTGGCAGATTG
PAH-tolerant male------------        ACCTCAGCGGGTTGCATGACTGCAGCGAGTGAAACCTGAGTGGCAGATTG
CYP1Arefseq------------------        ACCTCAGCGGGTTGCATGACTGCAGCGAGTGAAACCTGAGTGGCAGATTG
                                     **************************************************

PAH-sensitive female---------        CATGACTCCAAAGCTCCAAGATCCTCGCATTTACAGAAGCTATAGCGCGC
PAH-sensitive male-----------        CATGACTCCAAAGCTCCAAGATCCTCGCATTTACAGAAGCTATAGCGCGC
PAH-tolerant female----------        CATGACTCCAAAGCTCCAAGATCCTCGCATTTACAGAAGCTATAGCGCGC
PAH-tolerant male------------        CATGACTCCAAAGCTCCAAGATCCTCGCATTTACAGAAGCTATAGCGCGC
CYP1Arefseq------------------        CATGACTCCAAAGCTCCAAGATCCTCGCATTTACAGAAGCTATAGCGCGC
                                     **************************************************

PAH-sensitive female---------        AAATATGTGTACCTTCGTGCAGGCAAGGAGTGTGGGGCGAAAAA----AA
PAH-sensitive male-----------        AAATATGTGTACCTTCGTGCAGGCAAGGAGTGTGGGGCGAAAAAGAAAAA
PAH-tolerant female----------        AAATATGTGTACCTTCGTGCAGGCAAGGAGTGTGGGGCGAAAAA----AA
PAH-tolerant male------------        AAATATGTGTACCTTCGTGCAGGCAAGGAGTGTGGGGCGAAAAA----AA
CYP1Arefseq------------------        AAATATGTGTGCCTTCGTGCAGGCAAGGAGTGTGGGGCGAAAAA----AA
                                     ********** *********************************    **
```

```
PAH-sensitive female---------       AAAAGACTCGAGCTTTTCTCTCTTTGTTCTGCGAGTTTAAAACCTTTGTG
PAH-sensitive male-----------       AAAAGACTCGAGCTTTTCTCTCTTTGTTCTGCGAGTTTAAAACCTTTGTG
PAH-tolerant female----------       AAAAGACTCGAGCTTTTCTCTCTTTGTTCTGCGAGTTTAAAACCTTTGTG
PAH-tolerant male------------       AAAAGACTCGAGCTTTTCTCTCTTTGTTCTGCGAGTTTAAAACCTTTGTG
CYP1Arefseq------------------       AAAAGACTCGAGCTTTTCTCTCTTTGTTCTGCGAGTTTAAAACCTTTGTG
                                    **************************************************

PAH-sensitive female---------       TCTTTTCCAGAGGCTACAGCCTT-CGCGTGACTTAACCAGGCAGATTTAG
PAH-sensitive male-----------       TCTTTTCCAGAGGCTACAGCCTGGCGCGTGACTTAACCAGGCAGATTTAG
PAH-tolerant female----------       TCTTTTCCAGAGGCTACAGCCTTGCGCGTGACTTAACCAGGCAGATTTAG
PAH-tolerant male------------       TCTTTTCCAGAGGCTACAGCCTT-CGCGTGACTTAACCAGGCAGATTTAG
CYP1Arefseq------------------       TCTTTTCCAGAGGCTACAGCCTT-CGCGTGACTTAACCAGGCAGATTTAG
                                    **********************  **************************

PAH-sensitive female---------       GATTGTCCAGAAGACCGAGACCCTCCCCTCTGGACGGCAGCTCCGCTTCT
PAH-sensitive male-----------       GATTGTCCAGAAGACCGAGACCCTCCCCTCTGGACGGCAGCTCCGCTTCT
PAH-tolerant female----------       GATTGTCCAGAAGACCGAGACCCTCCCCTCTGGACGGCAGCTCCGCTTCT
PAH-tolerant male------------       GATTGTCCAGAAGACCGAGACCCTCCCCTCTGGACGGCAGCTCCGCTTCT
CYP1Arefseq------------------       GATTGTCCAGAAGACCGAGACCCTCCCCTCTGGACGGCAGCTCCGCTTCT
                                    **************************************************

PAH-sensitive female---------       CACGCAAGCTGCAGCACCAGAGGCGCAGCGCCGGGGCGCACGGAGAAAGA
PAH-sensitive male-----------       CACGCAAGCTGCAGCACCAGAGGCGCAGCGCCGGGGCGCACGGAAAAAGA
PAH-tolerant female----------       CACGCAAGCTGCAGCACCAGAGGCGCAGCGCCGGGGCGCACGGAGAAAGA
PAH-tolerant male------------       CACGCAAGCTGCAGCACCAGAGGCGCAGCGCCGGGGCGCACGGAGAAAGA
CYP1Arefseq------------------       CACGCAAGCTGCAGCACCAGAGGCGCAGCGCCGGGGCGCACGGAGAAAGA
                                    ******************************************** *****

PAH-sensitive female---------       GGCCCTGCAGAGTCAACACATGCAGCCCCCCAGGG--GATCAGGACAGCG
PAH-sensitive male-----------       GGCCCTGCAGAGTCAACACATGCAGCCCCCTTAGACAAAACAGGACAGCG
PAH-tolerant female----------       GGCCCTGCAGAGTCAACACATGCAGCTCCCCAGGG--GATCAGGACTGCG
PAH-tolerant male------------       GGCCCTGCAGAGTCAACACATGCAGCTCCCCAGGG--GATCAGGACTGCG
CYP1Arefseq------------------       GGCCCTGCAGAGTCAACACATGCAGCCCCCCAGGG--GATCAGGACAGCG
                                    ************************** ***   *    * ****** ***

PAH-sensitive female---------       CTGCGCTGCGCCGCGCCGACATAGTGGCACTGACGAACAAATTAAGTTTT
PAH-sensitive male-----------       CT-----GCGCTGCGCCGACATAGTGGCACTGACGAACAAATTAAGTTTT
PAH-tolerant female----------       CT-----GCGCTGCGCCGACATAGTGGCACTGACGAACAAATTAAGTTTT
PAH-tolerant male------------       CT-----GCGCTGCGCCGACATAGTGGCACTGACGAACAAATTAAGTTTT
CYP1Arefseq------------------       CT-----GCGCCGCGCCGACATAGTGGCACTGACGAACAAATTAAGTTTT
                                    **     **** **************************************
```

```
PAH-sensitive female---------        GCCTGTTTGACTTCGTTTAAAAAAAAAAGTCTAACTTTCTTGGAGGTTTG
PAH-sensitive male-----------        GCCTGTTTGACTTCGTTTAAAAAAAAAAGTCTAACTTTCTTGGAGGTTTG
PAH-tolerant female----------        GCCTGTTTGACTTCGTTTAAAAAAAAAAGTCTAACTTTCTTGGAGGTTTG
PAH-tolerant male------------        GCCTGTTTGACTTCGTTTAAAAAAAAAAGTCTAACTTTCTTGGAGGTTTG
CYP1Arefseq------------------        GCCTGTTTGACTTCGTATAAAAGAAAA-GTCTAACTTTCTTGGAGGTTTG
                                     **************** ***** **** **********************

PAH-sensitive female---------        TGTCCGGGGCCATTGGTTGTCAGCGCCCACGCTGTTT-AATGCGTTAACA
PAH-sensitive male-----------        TGTCCGGGGCCATTGGTTGTCAGCGCCCACGCTGTTTTAATGCGTTAACA
PAH-tolerant female----------        TGTCCGGGGCCATTGGTTGTCAGCGCCCACGCTGTTT-AATGCGTTAACA
PAH-tolerant male------------        TGTCCGGGGCCATTGGTTGTCAGCGCCCACGCTGTTT-AATGCGTTAACA
CYP1Arefseq------------------        TGTCCGGGGCCGTTGGTTGTCAGCGCCCACGCTGTTT-AATGCGTTAACA
                                     *********** ************************* ************

PAH-sensitive female---------        GGTTTTTGGTTTGGTTTGGTTTGTTGTTTTTTACGCATTCAGTTGGCAAA
PAH-sensitive male-----------        A-TTTTTTGTTTTGTTTTGTTTTTTGTTTTTTACGCATTCAGTTGGCAAA
PAH-tolerant female----------        G-TTTTTTGTTTTGTTTTGTTTTTTGTTTTTTACGCATTCAGTTGGCAAA
PAH-tolerant male------------        A-TTTTTTGTTTTGTTTTGTTTTTTGTTTTTTACGCATTCAGTTGGCAAA
CYP1Arefseq------------------        G-TTTTTTGTTTTGTTTTGTTTTTTGTTTTTTACGCATTCAGTTGGCAAA
                                       ***** **** **** **** ***************************

PAH-sensitive female---------        AACATATTTTGACAGCCTTCTCCATGAACGCCAAATAATTTATCACTGTC
PAH-sensitive male-----------        AACATATTTTGACAGCCTTCTCCATGAATGCCAAATAATTTATCACTGTC
PAH-tolerant female----------        AACATATTTTGACAGCCTTCTCCATGAATGCCAAATAATTTATCACTGTC
PAH-tolerant male------------        AACATATTTTGACAGCCTTCTCCATGAATGCCAAATAATTTATCACTGTC
CYP1Arefseq------------------        AACATATTTTGACAGCCTTCTCCATGAATGCCAAATAATTTATCACTGTC
                                     **************************** *********************

PAH-sensitive female---------        CTCTGATTGATCGTGCTATTTATGCCTCATTGCAAAGCTTTTTTTAATAC
PAH-sensitive male-----------        CTCTGATTGATCGTGCTATTTATGCCTCATTGTAAAGCTTTTTTTAATAC
PAH-tolerant female----------        CTCTGATTGATCGTGCTATTTATGCCTCATTGTAAAGCTTTTTTTAATAC
PAH-tolerant male------------        CTCTGATTGATCGTGCTATTTATGCCTCATTGTAAAGCTTTTTTTAATAC
CYP1Arefseq------------------        CTCTGATTGATCGTGCTATTTATGCCTCATTGTAAAGCTTTTTTTAATAC
                                     ******************************** *****************

PAH-sensitive female---------        ATATAATAAACTGTTGTCAAAATAGGTTATTTTAAACTATTGTACCAATA
PAH-sensitive male-----------        ATATAATAAACTGTTGTCAAAATAGGTTATTTTAAACTATTGTACCAATA
PAH-tolerant female----------        ATATAATAAACTGTTGTCAAAATAGGTTATTTTAAACTATTGTACCAATA
PAH-tolerant male------------        ATATAATAAACTGTTGTCAAAATAGGTTATTTTAAACTATTGTACCAATA
CYP1Arefseq------------------        ATATAATAAACTGTTGTCAAAATAGGTTATTTTAAACTATTGTACCAATA
                                     **************************************************
```

```
PAH-sensitive female---------      GCTGTCTGATTTTATTCACCCATTGCCTTCAAATTGTGTCACTTAACATG
PAH-sensitive male-----------      GCTGTCTGATTTTATTCACCCATTGCCTTCAAATTGTGTCACTTAACATG
PAH-tolerant female----------      GCTGTCTGATTTTATTCACCCATTGCCTTCAAATTGTGTCACTTAACATG
PAH-tolerant male------------      GCTGTCTGATTTTATTCACCCATTGCCTTCAAATTGTGTCACTTAACATG
CYP1Arefseq------------------      GCTGTCTGATTTTATTCACCCATTTCCTTTGAATTATGTCGCTTAACATG
                                   ************************ ****  **** **** *********

PAH-sensitive female---------      CAAATAAATAAATAAATAAATAGAATTTTGGGCAAAAATAGTTTTTGCCA
PAH-sensitive male-----------      CAAATAAATAAATAAATAAATAGAATTTTGGGCAAAAATAGTTTTTGCCA
PAH-tolerant female----------      CAAATAAATAAATAAATAAATAGAATTTTGGGCAAAAATAGTTTTTGTCA
PAH-tolerant male------------      CAAATAAATAAATAAATAAATAGAATTTTGGGCAAAAATAGTTTTTGTCA
CYP1Arefseq------------------      CAAATAAATAAATAAATAAATAGAATTTTGGGCAAAAATAGTTTTTGCCA
                                   *********************************************** **

PAH-sensitive female---------      CAAATTCTTTCCATAGATTGTTTAAATTGCATTTTATTTGACAGACAAAT
PAH-sensitive male-----------      CAAATTCTTTCCATAGATTGTTTAAATTGCATTTTATTTGACAGACAAAT
PAH-tolerant female----------      CAAATTCTTTCCATAGATTGTTTAAATTGCATTTTATTTGACAGACAAAT
PAH-tolerant male------------      CAAATTCTTTCCATAGATTGTTTAAATTGCATTTTATTTGACAGACAAAT
CYP1Arefseq------------------      CAAATTCTTTCCATAGATTGTTTAAATTGCATTTTATTTGACAGACAAAT
                                   **************************************************

PAH-sensitive female---------      ATGTTTAAAGTCCATATATCACCATGGAGCTAATTTATGTTAATATAACA
PAH-sensitive male-----------      ATGTTTAAAGTCCATATATCACCATGGAGCTAATTTATGTCAATATAACA
PAH-tolerant female----------      ATGTTTAAAGTCCATATATCACCATGGAGCTAATTTATGTCAATATAACA
PAH-tolerant male------------      ATGTTTAAAGTCCATATATCACCATGGAGCTAATTTATGTCAATATAACA
CYP1Arefseq------------------      ATGTTTAAAGTCCATATATCACCATGGTGCTAATTTATGTCAACAATACA
                                   *************************** ************ ** *  ***

PAH-sensitive female---------      ATTCAGAAAGTTACTGTATATTGATTCATTTCACACACGGCGTTAGAGTG
PAH-sensitive male-----------      ATTCAGAAAGTTACTG-ATATTGATTAATTTCACACACGGCGTTATATTT
PAH-tolerant female----------      ATTCAGAAAGTTACTG-ATATTGATTCATTTCACACACGGCGTTATATTT
PAH-tolerant male------------      ATTCAGAAAGTTACTG-ATATTGATTCATTTCACACACGGCGTTATATTT
CYP1Arefseq------------------      ATTCAGAAAGTTACTG-ATATTGATTCATTTCACACACGGCGTTATATTT
                                   **************** ********* ****************** * *

PAH-sensitive female---------      CGACGGGTATATTCATGATTGTGGCTTGATGATGACCCACAATTTAGTTT
PAH-sensitive male-----------      CTACTG-TTTATTCATGATTGTGGCTTGATGATGACCCACAATTTAGTTT
PAH-tolerant female----------      CTACTG-TTTATTCATGATTGTGGCTTGATGATGACCCACAATTTAGTTT
PAH-tolerant male------------      CTACTG-TTTATTCATGATTGTGGCTTGATGATGACCCACAATTTAGTTT
PAH-sensitive female---------      CGACGGGTATATTCATGATTGTGGCTTGATGATGACCCACAATTTAGTTT
CYP1Arefseq------------------      CTACTG-TTTATTCATGATTGTGGCTTGATGATGACCCACAATTTAGTTT
                                   * ** * * *****************************************
```

```
PAH-sensitive female---------       ATCAGAAAACTAGAATATAAGGCTGATAAAAACATGAATTGTTTTTCAAA
PAH-sensitive male-----------       ATCAGAAAACTAGAATATAAGGCTGATAAAAACATGAATTGTTTTTCAAA
PAH-tolerant female----------       ATCAGAAAACTAGAATATAAGGCTGATAAAAACATGAATTGTTTTTCAAA
PAH-tolerant male------------       ATCAGAAAACTAGAATATAAGGCTGATAAAAACATGAATTGTTTTTCAAA
CYP1Arefseq------------------       ATCAGAAAACTAGAATATAAGGCTGATAAAAACATGAATTGTTTTTCAAA
                                    **************************************************

PAH-sensitive female---------       CAGAAATATCAGCCCACAGACAGATACATGTACTGTATGGTAAAAACACT
PAH-sensitive male-----------       CAGAAATATCAGCCCACAGACAGATACATGTACTGTATGGTAAAAACACT
PAH-tolerant female----------       CAGAAATATCAGCCCACAGACAGATACATGTACTGTATGGTAAAAACACT
PAH-tolerant male------------       CAGAAATATCAGCCCACAGACAGATACATGTACTGTATGGTAAAAACACT
CYP1Arefseq------------------       CAGAAATATCAGCCCACAGACAGATACATGTACTGTATGGTAAAAACACT
                                    **************************************************

PAH-sensitive female---------       CAATACTTGGTCTGAGCTTGTCATCAATAACTGCATCGATGCAGTGTGGC
PAH-sensitive male-----------       CAATACTTGGTCTGAGCTTGTCATCAATAACAGCATCGATGCAGTGTGGC
PAH-tolerant female----------       CAATACTTGGTCTGAGCTTGTCATCAATAACAGCATCGATGCAGTGTGGC
PAH-tolerant male------------       CAATACTTGGTCTGAGCTTGTCATCAATAACAGCATCGATGCAGTGTGGC
CYP1Arefseq------------------       CAATACTTGGTCTGAGCTTGTCATCAATAACTGCATCGATGCAGTGTGGC
                                    ******************************* ******************

PAH-sensitive female---------       ATGAAGTCCGTGGTGCTGCTGAAGTCTTCAGGATCCAATATTGCTTTGAT
PAH-sensitive male-----------       ATGAAGTCTGTGGTGCTGCTGAAGTCTTCAGGATCCAATATTGCTTTGAC
PAH-tolerant female----------       ATGAAGTCTGTGGTGCTGCTGAAGTCTTCAGGATCCAATATTGCTTTGAC
PAH-tolerant male------------       ATGAAGTCTGTGGTGCTGCTGAAGTCTTCAGGATCCAATATTGCTTTGAC
CYP1Arefseq------------------       ATGAAGTCTGTGGTGCTGCTGAAGTCTTTAGGATCCATTATTGCTTTGAT
                                    ******** ******************* ******** ***********

PAH-sensitive female---------       AGCAGCTATAAGCGCATCTACATTTCTGCGTCTAGTGTTGCTCGTCGTCC
PAH-sensitive male-----------       AGCAGCTATAAGCGCATCTGCATTTCTGGGTCTGGTGTTGCTCGTCGTCC
PAH-tolerant female----------       AGCAGCTATAAGCGCATCTGCATTTCTGGGTCTGGTGTTGCTCGTTGTCC
PAH-tolerant male------------       AGCAGCTATAAGCGCATCTGCATTTCTGGGTCTGGTGTTGCTCGTTGTCC
CYP1Arefseq------------------       AGCAGCTATAAGCGCATCTGCATTTCTGCGTCTGGTGTTGCTCGTCGTCC
                                    ******************* ******** **** *********** ****

PAH-sensitive female---------       TCTTGACCGTGGTTTCTTAATTAGGTTTATGTAAGGGATATTTGCTGGCT
PAH-sensitive male-----------       TCTTGACCGTGGTTTCTTAATTAGGTTTAGGTAAGGGATATTTGCTGGCC
PAH-tolerant female----------       TCTTGACCGTGGTTTCTTAATTAGGTTTAGGTAAGGGATATTTGCTGGCC
PAH-tolerant male------------       TCTTGACCGTGGTTTTTTAATTAGGTTTAGGTAAGGGATATTTGCTGGCC
CYP1Arefseq------------------       TCTTGACCGTGGTTTCTTAATTAGGTTTAGGTAAGGGATATTTGCTGGCT
                                    *************** ************* *******************
```

```
PAH-sensitive female---------      AAACAAACGCTGTGATACTCTGGTTATTATAAGTTTAACTTGACCATCTG
PAH-sensitive male-----------      AAACAAACGCTGTGATACTCTGGTTATTATAAGTTTAACTTGACCGTCTG
PAH-tolerant female----------      AAACAAACGCTGTGATACTCTGGTTATTATAAGTTTAACTTGACCGTCTG
PAH-tolerant male------------      AAACAAACGCTGTGATACTCTGGTTATTATAAGTTTAACTTGACCGTCTG
CYP1Arefseq------------------      AAACAAACGCTGTGATACTCTGGTTATTATAAGTTTAACTTGACCATCTG
                                   ********************************************* ****

PAH-sensitive female---------      GTCAGGTGTCGTCCTGTTAAGAAATAAAAAACGGCGTCTTCTTAAAGCTT
PAH-sensitive male-----------      GTCAGGTGTCGTCCTGTTTAGAAATAAAAAACGGTGTCTTCTTAAAGCTT
PAH-tolerant female----------      GTCAGGGGTCGTCCTGTTTAGAAATAAAAAACGGTGTCTTCTTAAAGCTT
PAH-tolerant male------------      GTCAGGGGTCGTCCTGTTTAGAAATAAAAAACGGTGTCTTCTTAAAGCTT
CYP1Arefseq------------------      GTCAGGTGTCGTCCTGTTTAGAAATAAAAAACGGCGTCTTCTTAAAGCTT
                                   ****** ************ ************** ***************

PAH-sensitive female---------      ATCTGCAGAAGAAGAAGAATGACGTGCTCCAAAATATCCTGGTAATCAGT
PAH-sensitive male-----------      ATCTGCAGAGGAAGAAGAATGACGTGCTCCAAAATATCCTGGCAATCAGT
PAH-tolerant female----------      ATCTGCAGAGGAAGAAGAAAGACGTGCTCCAAAATATCCTGGCAATCAGT
PAH-tolerant male------------      ATCTGCAGAGGAAGAAGAATGACGTGCTCCAAAATATCCTGGCAATCAGT
CYP1Arefseq------------------      ATCTGCAGAAGAAGAAGAATGACGTGCTCCAAAATATCCTGGTAATCAGT
                                   ********* ********* ********************** *******

PAH-sensitive female---------      TGCACTGATTTTGTACTTTCAAATTAAAAAAA------CTAAATAAAAAA
PAH-sensitive male-----------      TGCACTGATTTTGTACTTTCCAATTAAAAAAA------CTAAATAAAAAA
PAH-tolerant female----------      TGCACTGATTTTGTACTTTCAAATAAAAAAAAAAAATAAAAAAAAAAAAA
PAH-tolerant male------------      TGCACTGATTTTGTACTTTCAAATAAAAAAAAAAAATAAAAAAAAAAAAA
CYP1Arefseq------------------      TGCACTGATTTTGTACTTTCAAATTAAAAAAA------CTAAATAAAAAA
                                   ******************** *** *******        *** ******

PAH-sensitive female---------      CACCGGACCAACATCAGCACATAACATGGTTCCCCAAATCATCACAGACT
PAH-sensitive male-----------      CACTGGACCAACATCAGCAAATAACATGGTTCCCCAAATCATCACAGACT
PAH-tolerant female----------      CACTGGACCAACATCAGCAAATAACAGGGTTCCCCAAATCATCACAGACT
PAH-tolerant male------------      CACTGGACCAACATCAGCAAAAAACAGGGTTCCCCAAATCATCACAAACT
CYP1Arefseq------------------      CACTGGACCAACATCAGCACATAACCTGGTTCCCCAAATCATCACAGACT
                                   *** *************** * ***  ******************* ***

PAH-sensitive female---------      GAGCTCTGAAAGCTGAATTCTCACGGTATTTGTTTGTTCACAAACACACA
PAH-sensitive male-----------      GAGCTCTGAAAGCTGAATTCTCACGGTATTTGTTTGATCACAAACACACA
PAH-tolerant female----------      GAGCTCTGAAAGCTGAATTCTCACGGTATTTGTTTGATCACAAACACACA
PAH-tolerant male------------      GAGCTCTGAAAGCGGAATTCCCACGGTATTTGTTTGATCACAAACACACA
CYP1Arefseq------------------      GAGCTCTGAAAGCTGAATTCTCACGGTATTTGTTTGTTCACAAACACACA
                                   ************* ****** *************** *************
```

```
PAH-sensitive female---------        AGAATCCATCTTGTTCCACTCCCGCCTCAACCATTCGAGACCGACCAATC
PAH-sensitive male-----------        AGAATCCATCTTGTTCCGCTCCCGCCTCAACCATTCGAGACCGACCAATC
PAH-tolerant female----------        AAAATCCATCTTGTTCCGCTCCCGCCTCAACCATTCGAGACCGACCAATC
PAH-tolerant male------------        AAAATCCATCTTGTTCCGCTCCCGCCTCAACCATTCGAGACCGACCAATC
CYP1Arefseq------------------        AGAATCCATCTTGTTCCGCTCCCGCCTCAACCATTCGAGACCGAGCAATC
                                     * *************** *************************** ****

PAH-sensitive female---------        ACGTTGGAGTATTATTATTTAAGTTGAGACATACAGCTGTGACAAATACA
PAH-sensitive male-----------        ACGTTGGAGTATTATTATTTAAGTTGAGACATACAGCTGTGACAAATACA
PAH-tolerant female----------        ACGTTGGAGTATTATTATTTAAGTTGAAACATACAGCTGTGACAAAAACA
PAH-tolerant male------------        ACGTTGGAGTATTATTATTTAAGTTGAAACATACAGCTGTGACAAAAACA
CYP1Arefseq------------------        ACGTTGGAGTATTATTATTTAAGTTGAGACATACAGCTGTGACAAATACA
                                     *************************** ****************** ***

PAH-sensitive female---------        ATAGCTGCCAAACC------------AAAATAAATATGGCAATGCAGGAG
PAH-sensitive male-----------        ATAGCTGCCAAACCCACAGCCAAACCAAAATGAATATGGCAATGCAGGAG
PAH-tolerant female----------        ATAGCTGCCAAACCCACAGCCAAACCAAAATAAATATGGCAATGCAGGAG
PAH-tolerant male------------        ATAGCTGCCAAACCCACAGCCAAACAAAAAAAAATATGGCAATGCAGGAG
CYP1Arefseq------------------        ATAGCTGCCAAACCCACAGCCAAACCAAAATAAATATGGCAATGCAGGAG
                                     **************            ****  ******************

PAH-sensitive female---------        GATGCATGCATAGCTGCTGCTATCACATGTGTGTTTTGTCAGAATCCATG
PAH-sensitive male-----------        GATGCATGCATAGCTGCTGCTATCACATGTGTGTTTTGTCAGAATCCATG
PAH-tolerant female----------        GATGCATGCATAGCTGCTGCTATCACATGTGTGTTTTGTCAGAATCCATG
PAH-tolerant male------------        GATGCATGCATAGCTGCTGCTATCACATGTGTGTTTTGTCAGAATCCATG
CYP1Arefseq------------------        GATGCATGCATAGCTGCTGCTATCACATGTGTGTTCTGTCAGAATCCATG
                                     *********************************** **************

PAH-sensitive female---------        ATTGTC---------ATTATTCAAATAAGTGCCCCTTGACCAGCTTAACT
PAH-sensitive male-----------        ATTGTCATGAGTGTCATTATTCAAATAAGTGCCCCTTGACCAGCTTAACT
PAH-tolerant female----------        ATTGTCATGAGTGTCATTATTCAAATAAGTGCCCCTTGACCAGCTTAACT
PAH-tolerant male------------        ATTGTCATGAGTGTCATTATTCAAATAAGTGCCCCTTGACCAGCTTAACT
CYP1Arefseq------------------        ATTGTC---------ATTATTCAAATAAGTGCCCCTTGACCAGCTTAACT
                                     ******         ***********************************

PAH-sensitive female---------        TTTTGTGGTTTTTCATGGCGCCTGCTGCTTACAGCATTTTCATTTGATAT
PAH-sensitive male-----------        TGTTGTGGTTTTTCATGGCACCTGCTGCTTCTAACATTTTCATTTGATAT
PAH-tolerant female----------        TGTTGTGGTTTTTCATGGCACCTGCTGCTTCTAACATTTTCATTTGATAT
PAH-tolerant male------------        TGTTGTGGTTTTTCATGGCACCTGCTGCTTCTAACATTTTCATTTGATAT
CYP1Arefseq------------------        TTTTGTGGTTTTTCATGGCGCCTGCTGCTTACAGCATTTTCATTTGATAT
                                     * ***************** **********  * ****************
```

```
PAH-sensitive female---------        CGTGATTGGCTAAAACCAACCTTTGGTAGGCGGGCTATAGGTGTCACTTC
PAH-sensitive male-----------        TGTGATTGGCTAAAACCACCCTTTGGTAAGCGGGCTATAGGTGTCACTTC
PAH-tolerant female----------        TGTGATTGGCTAAAACCACCCTTTGGTAAGCGGGCTATAGGTGTCACTTC
PAH-tolerant male------------        TGTGATTGGCTAAAACCACCCTTTGGTAAGCGGGCTATAGGTGTCACTTC
CYP1Arefseq------------------        CGTGATTGGCTAAAACCAACCTTTGGTAGGCGGGCTATAGGTGTCACTTC
                                      ***************** ********* *********************

PAH-sensitive female---------        ACCCAAACAGCTTGGAAAGTTCTGCTGGATGTCCCTCCTTTCCCCGAACA
PAH-sensitive male-----------        ACCCAAACAGCTTGGAAAGTTCTGCTGGATGTCCCTCCTTTCCCCGAACA
PAH-tolerant female----------        ACCCAAACAGCTTGGAAAGTTCTGCTGGATGTCCCTCCTTTCCCCGAACA
PAH-tolerant male------------        ACCCAAACAGCTTGGAAAGTTCTGCTGGATGTCCCTCCTTTCCCCGAACA
CYP1Arefseq------------------        ACCCAAACAGCTTGGAAAGTTCTGCTGGATGTCCCTCCTTTCCCCGAACA
                                     **************************************************

PAH-sensitive female---------        GATTTGATGGGAGCAAACCCAGATGGATGATGTGCAGAACGGAGAGTGAT
PAH-sensitive male-----------        GATTTGATGGGAGCAAACCCAGATGGAT---GTGCAGAACGGAGAGTGAT
PAH-tolerant female----------        GATTTGATGGGAGCAGACCCAGATGGATGATGTGCAGAACGGAGAGTGAT
PAH-tolerant male------------        GATTTGATGGGAGCAGACCCAGATGGATGATGTGCAGAACGGAGAGTGAT
CYP1Arefseq------------------        GATTTGATGGGAGCAAACCCAGATGGATGATGTGCAGAACGGAGAGTGAT
                                     *************** ************   *******************

PAH-sensitive female---------        ACACATCCCCACTACACTATCAATCTGGCTTGCCAGGTTAGCACTGGCCG
PAH-sensitive male-----------        ACACATCTCTGCTGCACTATCAATCTGGCTTGCCAGGTTAGCACTGGCCG
PAH-tolerant female----------        ACACATCCCCGCTACACTATCAATCTGGCTTGCCAGGTTAGCACTGGCCG
PAH-tolerant male------------        ACACATCCCCGCTACACTATCAATCTGGCTTGCCAGGTTAGCACTGGCCG
CYP1Arefseq------------------        ACACATCCCCACTACACTATCAATCTGGCTTGCCAGGTTAGCACTGGCCG
                                     ******* *  ** ************************************

PAH-sensitive female---------        TGGAAACTTCACACTGGACCTCAAGCAAATTGGATTCTATGCCTCTCTCT
PAH-sensitive male-----------        TGGAAACTTCACACTGGACCTCAAGCAAATTGGATTCTATGCCTCTCTCT
PAH-tolerant female----------        TGGAAACTTCACACTGGACCTCAAGCAAATTGGATTCTATGCCTCTCTCT
PAH-tolerant male------------        TGGAAACTTCACACTGGACCTCAAGCAAATTGGATTCTATGCCTCTCTCT
CYP1Arefseq------------------        TGGAAACTTCACACTGGACCTCAAGCAAATTGGATTCTATGCCTCTCTCT
                                     **************************************************

PAH-sensitive female---------        TCCTTCAGAGCTCAGGACCATCGTTTTCCACATGAAACGCAAAAGTTTGC
PAH-sensitive male-----------        TCCTTCAGAGCTCAGGACCATCGTTTTCCACATGAAACGCAAA-GTTTGC
PAH-tolerant female----------        TCCTTCAGAGCTCAGGACCATCGTTTTCCACATGAAACGCAAA-GTTTGC
PAH-tolerant male------------        TCCTTCAGAGCTCAGGACCATCGTTTTCCACATGAAACGCAAA-GTTTGC
CYP1Arefseq------------------        TCCTTCAGAGCTCAGGACCATCGTTTTCCACATGAAACGCAAA-GTTTGC
                                     ******************************************* ******
```

```
PAH-sensitive female---------        TTTTAGCAAAAAAGAATTAGCCACAGTCCAGCCATTTTCTCCTTAATCAA
PAH-sensitive male-----------        TTTTAGCAAAAAAGAATTAGCCACAGTCCAGTCATTTTCTCCTTGATCAA
PAH-tolerant female----------        TTTTAGCAAAAAAGAATTAGCCACAGTCCAGTCATTTTCTCCTTGATCAA
PAH-tolerant male------------        TTTTAGCAAAAAAGAATTAGCCACAGTCCAGTCATTTTCTCCTTGATCAA
CYP1Arefseq------------------        TTTTAGCAAAAAAGAATTAGCCACAGTCCAGCCATTTTCTCCTTAATCAA
                                     ******************************* ************ *****

PAH-sensitive female---------        AGTGCGTTTCTCTGGTGGTCAAAACGTTGATTCTTATTGCACCTTTTTGT
PAH-sensitive male-----------        AGTGCGTTTCTCTGGTGGTCAAAACGTTGATTCTTAGTGCACCTTTT-GT
PAH-tolerant female----------        AGTGCGTTTCTCTGGTGGTCAAAACGTTGATTCTTAGTGCACCTTTT-GT
PAH-tolerant male------------        AGTGCGTTTCTCTGGTGGTCAAAACGTTGATTCTTAGTGCACCTTTT-GT
CYP1Arefseq------------------        AGTGCGTTTCTCTGGTGGTCAAAACGTTGATTCTTAGTGCACCTTTTTGT
                                     ************************************ ********** **

PAH-sensitive female---------        TCCACACTTTTTCCTTCCACTAAACTTTCCACCATCATATTTTGTTACTG
PAH-sensitive male-----------        TCCACACTTTTTCCTTCCACTAAATTTTCCACCATCATATTTTGTTACTG
PAH-tolerant female----------        TCCACACTTTTTCCTTCCACTAAATTTTCCACCATCATATTTTGTTACTG
PAH-tolerant male------------        TCCACACTTTTTCCTTCCACTAAATTTTCCACCATCATATTTTGTTACTG
CYP1Arefseq------------------        TCCACACTTTTTCCTTCCACTAAACTTTCCACCATCATATTTTGTTACTG
                                     ************************ *************************

PAH-sensitive female---------        CACCATGTGAAAAGATGACTCGTTTT--TAGCAACGACCTCTTGTGGTTC
PAH-sensitive male-----------        CACCATG--AAAAGCCGACTCTTTTTTTTATCAACGACCTC---------
PAH-tolerant female----------        CACCATGTGAAAAGCCGACTCTTTTTTTTATCAACGACCTC---------
PAH-tolerant male------------        CACCATGTGAAAAGCCGACTCTTTTTTTTATCAACGACCTC---------
CYP1Arefseq------------------        CACCATGTGAAAAGATGACTCGTTTT--TAGCAACGACCTCTTGTGGTTC
                                     *******  *****  ***** ****  ** **********

PAH-sensitive female---------        ATCCTCCTGGTAGAATAAAATGCTCAAATAAGCAGTCTTTTTCATGATTG
PAH-sensitive male-----------        ------CTGGTAGAATAAAATGCTCAAATGAGCAGTCTTTTTCATGATTG
PAH-tolerant female----------        ------CTGGTAGAATAAAATGCTCAAATGAGCAGTCTTTTTCATGATTG
PAH-tolerant male------------        ------CTGGAACAATAAAATGCTCAAATGAGCAGTCTTTTTCATGAATG
CYP1Arefseq------------------        ATCTTCCTGGTAGAATAAAATGCTCAAATAAGCAGTCTTTTTCATGATTG
                                           **** * **************** ***************** **

PAH-sensitive female---------        TATTGGTCATGTCTATAATAAACTTCTACAAGCAATCCATTCTTATTGTT
PAH-sensitive male-----------        TATTGGTCATGGCTATAATAAACTTCTATAAGCAATCATTCCTTATCGTT
PAH-tolerant female----------        TATTGGTCATGGCTATAATAAACTTCTATAAGCAATCCATTCTTATCGTT
PAH-tolerant male------------        TATTGGTCATGGCTATAATAAACTTCTATAAGCAATCCATTCTTATCGTT
CYP1Arefseq------------------        TATTGGTCATGTCTATAATAAACTTCTACAAGCAATCCATTCTTATTGTT
                                     *********** **************** ********  * ***** ***
```

```
PAH-sensitive female---------       CTTAAGAAATATAATTTTCGTAAACAGTTTTTAATTTGTGATTTGCTTGA
PAH-sensitive male-----------       CTTAAGAAATATAATTTTTGTAAACAGTTTTGAGTT-GTGAT--------
PAH-tolerant female----------       CTTAAGAAATATCATTTTTGTAAACAGTTTTGAGTT-GTGAT--------
PAH-tolerant male------------       CTTAAGAAATATCATTTTTGTAAACAGTTTTGAGTT-GTGAT--------
CYP1Arefseq------------------       CTTAAGAAATATAATTTTCGTAAACAGTTTTTAATTTGTGATTTGCTTGA
                                    ************ ***** ************ * ** *****

PAH-sensitive female---------       AATCTATGTACAAGGAATCCTTATTGTAGCTAAACATATTTTCATGTCTA
PAH-sensitive male-----------       ---------------------------AGCTAAACATATTTCCATGTCTA
PAH-tolerant female----------       ---------------------------AGCTAAACATATTTCCATGTCTA
PAH-tolerant male------------       ---------------------------AGCTAAACATATTTCCATGTCTA
CYP1Arefseq------------------       AATCTATGTACAAGGAATCCTTATTGTAGCTAAACATATTTTCATGTCTA
                                                               ************** ********

PAH-sensitive female---------       AACCTTTCAGTGATACTCAGCTCCATTTAAATACACATTCTCTTCCAGGA
PAH-sensitive male-----------       AACTTTTCAGCGATACTCAGCTCCATTTAAATACACATTCTCTTCCAGGA
PAH-tolerant female----------       AACTTTTCAGCGATACTCAGCTCCATTTAAATACACATTCTCTTCCAGGA
PAH-tolerant male------------       AACTTTTCAGCGATACTCAGCTCCATTTAAATACACATTCTCTTCCAGGA
CYP1Arefseq------------------       AACCTTTCAGTGATACTCAGCTCCATTTAAATACACATTCTCTTCCAGGA
                                    *** ****** ***************************************

PAH-sensitive female---------       CATGTAAAACAAATTTCATCACAATAATGGATACCTGGAAGATTTTCTCT
PAH-sensitive male-----------       CATGTAAAACAAATTTCATCACAATAATGGATACCTGGAAGCTTTTCTCT
PAH-tolerant female----------       CATGTAAAACAAATTTCATCACAATAATGGATACCTGGAAGCTTTTCTCT
PAH-tolerant male------------       CATGTAAAACAAATTTCATCACAATAATGGATACCTGGAAGCTTTTCTCT
CYP1Arefseq------------------       CATGTAAAACAAATTTCATCACAATAATGGATACCTGGAAGATTTTCTCT
                                    ***************************************** ********

PAH-sensitive female---------       TGCTCTGCAGGGATTTAACAGATTTTTTTTTTTATTATTATTGTATAATG
PAH-sensitive male-----------       TGCTCTGCAGGGATTCAACAGATTTTTGTTTTTATT------GTATAATG
PAH-tolerant female----------       TGCTCTGCAGGGATTCAACAGATTTTTGTTTTAATT------GTATAATG
PAH-tolerant male------------       TGCTCTGCAGGGATTCAACAGATTTTTGTTTTAATT------GTATAATG
CYP1Arefseq------------------       TGCTCTGCAGGGATTTAACAGATTTTTTTTTTTATTATTATTGTATAATG
                                    *************** *********** **** ***      ********

PAH-sensitive female---------       TTACACCACATATCGGTTAAAGGGAGAAATATTTTAGGTTAACTAAAAAC
PAH-sensitive male-----------       TTACACCACATATCGGTAAAAGGAATAAATATTTCAGGTTAACTAAAAAC
PAH-tolerant female----------       TTACACCACATATCGGTAAAAGGAATAAATATTTCAGGTTAACTAAAAAC
PAH-tolerant male------------       TTACACCACATATCGGTAAAAGGAATAAATATTTCAGGTTAACTAAAAAC
CYP1Arefseq------------------       TTACACCACATATCGGTTAAAGGGAGAAATATTTTAGGTTAACTAAAAAC
                                    ***************** ***** * ******** ***************
```

```
PAH-sensitive female---------        TGAAGTATAGCTGAGAAACAGTTTGGTGCGCTCATTGTTCCCATTTTTTT
PAH-sensitive male-----------        TAAAGTATAGCTGAAAAACAGTTTGGTGCGCTCATTGTTCCCATTTTTTT
PAH-tolerant female----------        TAAAGTATAGCTGAGAAACAGTTTGGTGCGCTCATTGTTCCCATTTTTTT
PAH-tolerant male------------        TAAAGTATAGCTGAGAAACAGTTTGGTGCGCTCATTGTTCCCATTTTTTT
CYP1Arefseq------------------        TGAAGTATAGCTGAGAAACAGTTTGGTGCGCTCATTGTTCCCATTTTTTT
                                     * ************ ***********************************

PAH-sensitive female---------        TTTCCATTTACATGTTTAAAGCCGTTTATCCAAGTTGTAATGCTGCATTA
PAH-sensitive male-----------        ---CCATTTACTTGTTTAGAGCCGTTTATCCAAGTTGTAATGCTGCATTA
PAH-tolerant female----------        ---CCATTTACTTGTTTAGAGCCGTTTATCCAAGTTGTAATGCTGCATTA
PAH-tolerant male------------        ---CCATTTACTTGTTTAGAGCCGTTTATCCAAGTTGTAATGCTGCATTA
CYP1Arefseq------------------        TTTCCATTTACATGTTTAAAGCCGTTTATCCAAGTTGTAATGCTGCATTA
                                        ******** ****** *******************************

PAH-sensitive female---------        TTTAAGCCTCCCGTTGCGCAGCCTTTAATAAACGTAATGAACTCGTCTGC
PAH-sensitive male-----------        TTTAAGCCTCCCAGTGCGCAGCCTTTAATAAACGTAATGAACTCGTCTGC
PAH-tolerant female----------        TTTAAGCCTCCCAGTGCGCAGCCTTTAATAAACGTAATGAACTCGTCTGC
PAH-tolerant male------------        TTTAAGCCTCCCAGTGCGCAGCCTTTAATAAACGTAATGAACTCGTCTGC
CYP1Arefseq------------------        TTTAAGCCTCCCAGTGCGCAGCCTTTAATAAACGTAATGAACTCGTCTGC
                                     ************  ************************************

PAH-sensitive female---------        CCTTGCGCTCCGGCAGCAGTGGCGCCTTCTCCCCTCCCTGTCCGTGCCCA
PAH-sensitive male-----------        CCTTGCGCTCCGGCAGCAGTGGCGCCTCCTCCCCTCCCTGTCCGTGCCCA
PAH-tolerant female----------        CCTTGCGCTCCGGCAGCAGTGGCGCCTCCTCCCCTCCCTGTCCGTGCCCA
PAH-tolerant male------------        CCTTGCGCTCCGGCAGCAGTGGCGCCTCCTCCCCTCCCTGTCCGTGCCCA
CYP1Arefseq------------------        CCTTGCGCTCCGGCAGCAGTGGCGCCTCCTCCCCTCCCTGTCCGTGCCCA
                                     *************************** **********************

PAH-sensitive female---------        GCATCCTCCTCGAAGGGGAGAGGGCGGTTTGATCACTGCGCTCTCACGCA
PAH-sensitive male-----------        GCATCCTCCTCGAAGGGGAGAGGGCGGTTTGATCACTGCGCTCTCACGCA
PAH-tolerant female----------        GCATCCTCCTCGAAGGGGAGAGGGCGGTTTGATCACTGCGCTCTCACGCA
PAH-tolerant male------------        GCATCCTCCTCGAAGGGGAGAGGGCGGTTTGATCACTGCGCTCTCACGCA
CYP1Arefseq------------------        GCATCCTCCTCGAAGGGGAGAGGGCGGTTTGATCACTGCGCTCTCACGCA
                                     **************************************************
                                                                                  XRE3

PAH-sensitive female---------        ACTGGTCAATCTTTAACTCCCGCGGAGAGCATGCAGGTACAAGCACGCAA
PAH-sensitive male-----------        ACTGGTCAATCTTTAACTCCCGCGGAGAGCATGCAGGTACAAGCACGCAA
PAH-tolerant female----------        ACTGGTCAATCTTTAACTCCCGCGGAGAGCATGCAGGTACAAGCACGCAA
PAH-tolerant male------------        ACTGGTCAATCTTTAACTCCCGCGGAGAGCATGCAGGTACAAGCACGCAA
CYP1Arefseq------------------        ACTGGTCAATCTTTAACTCCCGCGGAGAGCATACAGGTACAAGCACGCAA
                                     ********************************* ****************
                                                                                XRE2
```

```
PAH-sensitive female---------        TTGCATCTGTTTT-ATCAGCACTGCGCAACCTTGCCCGGAAAATGCTGGC
PAH-sensitive male-----------        TTGCATCTGTTTTTATCAGCACTGCGCAACCTTGCCCGGAAAATGCTGGC
PAH-tolerant female----------        TTGCATCTGTTTTTATCAGCACTGCGCAACCTTGCCCGGAAAATGCTGGC
PAH-tolerant male------------        TTGCATCTGTTTTTATCAGCACTGCGCAACCTTGCCCGGAAAATGCTGGC
CYP1Arefseq------------------        TTGCATCTGTTTTTATCAGCACTGCGCAACCTTGCCCGGAAAATGCTGGC
                                     ************* ************************************

PAH-sensitive female---------        TGGCATGGCAAGCAGCAGCCCCGTTCTCACCCCCAAATCTGGGTGGTAAG
PAH-sensitive male-----------        TGGCATGGCAAGCAGCAGCCCGGTTCTCACCCCCAAATCTGGGGGGTAAG
PAH-tolerant female----------        TGGCATGGCAAGCAGCAGCCCCGTTCTCACCCCCAAATCTGGGTGGTAAG
PAH-tolerant male------------        TGGCATGGCAAGCAGCAGCCCCGTTCTCACCCCCAAATCTGGGTGGTAAG
CYP1Arefseq------------------        TGGCATGGCAAGCAGCAGCCCCGTTCTCACCCCCAAATCTGGGTGGTAAG
                                     ********************* ********************* ******

PAH-sensitive female---------        GTGGTTGAACGTCTGCCTGATGTCGCAACAGTCACAAGCACATAACCGTG
PAH-sensitive male-----------        GGGGTTGAACGTCTGCCTGATGTCCCAACAGTCACAAGCACATAACCGTG
PAH-tolerant female----------        GGGGTTGAACGTCTGCCTGATGTCGCAACAGTCACAAGCACATAACCGTG
PAH-tolerant male------------        GTGGTTGAACGTCTGCCTGATGTCGCAACAGTCACAAGCACATAACCGTG
CYP1Arefseq------------------        GTGGTTGAACGTCTGCCTGATGTCGCAACAGTCACAAGCACATAACCGTG
                                     * ********************** *************************

PAH-sensitive female---------        CTACTTAATAATAAGTTACTTATTTTTAATGCAAAGAAATAAAAAAATAA
PAH-sensitive male-----------        CTACTTAATAATAAGTTACTTATTTTTAATGCAAAGGAATAAAA----TA
PAH-tolerant female----------        CTACTTAATAATAAGTTACTTATTTTTAATGCAAAGGAATTAAA----AA
PAH-tolerant male------------        CTACTTAATAATAAGTTACTTATTTTTAATGCAAAGGAATTAAA----AA
CYP1Arefseq------------------        CTACTTAATAATAAGTTACTTATTTT-AATGCAAAGGAATAA-----TAT
                                     ************************** ********* *** *

PAH-sensitive female---------        AAAAAAAAGCTCATTCACAGATCGCGCTCGCACAACGCCTGCGTCAGCAA
PAH-sensitive male-----------        AAAAAAAAGCTCATTCACAGATCGCGCTCGCACAACGCCTGCGTCAGCAA
PAH-tolerant female----------        AAAAAAAAGCTCATTCACAGATCGCGCTCGCACAACGCCTGCGTCAGCAA
PAH-tolerant male------------        AAACAAAAGCTCATTCACAGATCGCGCTCGCACAACGCCAGCGTCAGCAA
CYP1Arefseq------------------        AAAAAAAAGCTCATTCACAGATCGCGCTCGCACAACGCCTGCGTCAGCAA
                                     *** ************************************ *********

PAH-sensitive female---------        AATGCCAACCTGCACAGATCAAGACCAAGTGCATTAGAATGGATACAAAC
PAH-sensitive male-----------        AATGCCAACCCGCACAGATCAAGACCAAGTGCATTAGAAAGGATACAAAC
PAH-tolerant female----------        AATGCCAACCAGCACAGATCAAGACCAAGTGCATTAGAATGGATACAACC
PAH-tolerant male------------        AATGCCAACCAGCACAGATCAAGACCAAGTGCATTAGAATGGATACAACC
CYP1Arefseq------------------        AATGCCAACCTGCACAGATCAAGACCAAGTGCATTAGAATGGATACAAAC
                                     ********** **************************** ******** *
```

```
PAH-sensitive female---------       CTTAAATTATA-CAACAGTATTAGATATTAGGGCTGGCAAAAAATAAAAT
PAH-sensitive male-----------       CTTAAATTATAACAACAGTATTAGATATTAGGGCTGGCAAAAAATAAAAT
PAH-tolerant female----------       CTTAAATTATA-CAATAGTATTAGAAAATAGGGCTGGCAAAAAG------
PAH-tolerant male------------       CTTAAATTATA-CAATAGTATTAGAAAATAGGGCTGGCAAAAAG------
CYP1Arefseq------------------       CTTAAATTATA-CAACAGTATTAGATATTAGGGCTGGCAAAAAATAACAT
                                    *********** *** ********* * ***************

PAH-sensitive female---------       AAAATAAA----ATATTGCACAGAAAATAAAAGCAAACACTGCAAACTCT
PAH-sensitive male-----------       AAAATAAA----ATATTGCACAGAAAATAAAAGCAAACACTGCAAACTCT
PAH-tolerant female----------       ------AA----ATATTGCACAGAAAATAAAAGCAAACACTGCAAACTCT
PAH-tolerant male------------       ------AA----ATATTGCACAGAAAATAAAAGCAAACACTGCAAACTCT
CYP1Arefseq------------------       AAAATAAATAAAATATTGCACAGAAAATAAAAGCAAACACTGCAAACTCT
                                          **    **************************************

PAH-sensitive female---------       GAACCACCTGCAGAAAGGCGCACAGTGATAAACATTTGAATGCTCTTATC
PAH-sensitive male-----------       GAACCACCTGCAGAAAGGCGCACAGTGATAAAGATTTGAATGCTCTTATC
PAH-tolerant female----------       GAACCACCTGCAGAAAGGCGCACAGTGATAAAGATTTGAATGCTCTTATC
PAH-tolerant male------------       GAACCACCTGCAGAAAGGCGCACAGTGATAAAGATTTGAATGCTCTTATC
CYP1Arefseq------------------       GAACCACCTGCAGAAAGGCGCACAGTGATAAACATTTGAATGCTCTTATC
                                    ******************************** *****************

PAH-sensitive female---------       GCATAACAGACGCTGATTTGCAGCCCGCCTTTGTCAGCATTGTGTCTCAT
PAH-sensitive male-----------       GCATAACAGACGCTGATTTGCAGCCCGCCTTTGACAGCATTGTGTCTCAT
PAH-tolerant female----------       GCATATCAGACGCTGATTTGCAGCCCGCCTTTGACAGCATTGTGTCTCAT
PAH-tolerant male------------       GCATATCAGACGCTGATTTGCAGCCCGCCTTTGACAGCATTGTGTCTCAT
CYP1Arefseq------------------       GCATATCAGACGCTGATTTGCCGCCCGCCTTTGACAGCATTGTGTCTCAT
                                    ***** *************** *********** ****************

PAH-sensitive female---------       GCGCACCTAAACTTGAAGAAGGCGGTAGACACTTTGTAATGCACGCGAAT
PAH-sensitive male-----------       GCGCACCTAAACTTGAACAAGGCGGTAGACACTTTGTAATGCCCGCGAAT
PAH-tolerant female----------       GCGCACCTAAACTTGAACAAGGCGGTAGACACTTTGTAATGCACGCGAAT
PAH-tolerant male------------       GCGCACCTAAACTTGAACAAGGCGGTAGACACTTTGTAATGCACGCGAAT
CYP1Arefseq------------------       GCGCACCTAAACTTGAAGAAGGCGGTAGACACTTTGTAATGCACGCGAAT
                                    ***************** ************************ *******
                                                                              XRE1
PAH-sensitive female---------       TGTGTACCGCCAGGACCACACACAGACACACCCACCAACTTTTTTTTTT-
PAH-sensitive male-----------       TGTGTACCGCCAGGACCACACACAGACACCCCCACCAACTTTTTTTTTTT
PAH-tolerant female----------       TGGGTACCCCCAGGACCACACACAGACACACCCACCAACTTTTTTTTTTT
PAH-tolerant male------------       TGTGTACCCCCAGGACCACACACAGACACACCCACCAACTTTTTTTTTTT
CYP1Arefseq------------------       TGTGTACCGCCAGGACCACACACAGACACACCCACCAACTTTTTTTTTTT
                                    ** ***** ******************** *******************
```

```
PAH-sensitive female---------     ---CTACTGCTCCAAACTTCATTCATGGCAGGGAATTAAAGACAGGCACT
PAH-sensitive male-----------     TTTCTACTGCTCCAAACTTCATTCATGGCAGGGAATTAAAGACAGGCACT
PAH-tolerant female----------     ---CTACTGCTCCAAACTTCATTCATGGCAGGGAATTAAAGACAGGCACT
PAH-tolerant male------------     ---CTACTGCTCCAAACTTCATTCATGGCAGGGAATTAAAGACAGGCACT
CYP1Arefseq------------------     TT-CTACTGCTCCAAACTTCATTCATGGCAGGGAATTAAAGACAGGCACT
                                     ***********************************************

PAH-sensitive female---------     CGGATGGAGGAGGGGAGATGATGTCAACCTCGGTAGCCAATAAGATTGCG
PAH-sensitive male-----------     CGGATGGAGGAGGGGAGATGATGTCAACCTCGGTAGCCAATAAGATTGCG
PAH-tolerant female----------     CAGATGGAGGAGGGGAGATGATGTCAACCTCGGTAGCCAATAAGATTGCG
PAH-tolerant male------------     CAGATGGAGGAGGGGAGATGATGTCAACCTCGGTAGCCAATAAGATTGCG
CYP1Arefseq------------------     CAGATGGAGGAGGGGAGATGATGTCAACCTCGGTAGCCAATAAGATTGCG
                                  * ************************************************

PAH-sensitive female---------     CAGCGCTCTATAAATCATACGTCCACTCGCGGCTTTGAAGACATCTGCAA
PAH-sensitive male-----------     CAGCGCTCTATAAATCATACGTCCACTCGCGGCTTTGAAGACATCTGCAA
PAH-tolerant female----------     CAGCGCTCTATAAATCATACGTCCACTCGCGGCTTTGAAGACATCTGCAA
PAH-tolerant male------------     CAGCGCTCTATAAATCATACGTCCACTCGCGGCTTTGAAGACATCTGCAA
CYP1Arefseq------------------     CAGCGCTCTATAAATCATACGTCCACTCGCGGCTTTGAAGACATCTGCAA
                                  **************************************************
                                         TATAA                               TSS

PAH-sensitive female---------     CGTTGAGGACACCTCTGCAAAACACATTTTTTTTCTGTTGTTGGACACGC
PAH-sensitive male-----------     CGTTGAGGACACCTCTGCAAAACACATTTTTTT-CTGTTGTTGGACACGC
PAH-tolerant female----------     CGTTGAGGACACCTCTGCAAAACACATTTTTTTTCTGTTGTTGGACACGC
PAH-tolerant male------------     CGTTGAGGACACCTCTGCAAAACACATTTTTTTTCTGTTGTTGGACACGC
CYP1Arefseq------------------     CGTTGAGGACACCTCTGCAAAACACATTTTTTTTCTGTTGTTGGACACGC
                                  ********************************* ****************

PAH-sensitive female---------     ATCTCTGGAATTAGAGTGTTCGTCTTCTTTTTTT-ATCATTCAGCTAAAG
PAH-sensitive male-----------     ATCTCTGGAATTAGAGTGTTCGTCTTCTTTTTTTTATCATTCAGCTAAAG
PAH-tolerant female----------     ATCTCTGGAATTAGAGTGTTCGTCTTCTTTTTTTTATCATTCAGCTAAAG
PAH-tolerant male------------     ATCTCTGGAATTAGAGTGTTCGTCTTCTTTTTTTTATCATTCAGCTAAAG
CYP1Arefseq------------------     ATCTCTGGAATTAGAGTGTTCGTCTTCTTTTTTT-ATCATTCAGCTAAAG
                                  ********************************** ***************

PAH-sensitive female---------     GTAAGGTGATTACCTGCATGATAAAAGTTATTTCTAAGAAAGGATTGAGA
PAH-sensitive male-----------     GTAAGGTGATTACCTGCATGATAAAAGTTATTTCTAAGAAAGGATTGAGA
PAH-tolerant female----------     GTAAGGTGATTACCTGCATGATAAAAGTTATTTCTAAGAAAGGATTGAGA
PAH-tolerant male------------     GTAAGGTGATTACCTGCATGATAAAAGTTATTTCTAAGAAAGGATTGAGA
CYP1Arefseq------------------     GTAAGGTGATTACCTGCATGATAAAAGTTATTTCTAAGAAAGGATTGAGA
                                  **************************************************
```

```
PAH-sensitive female---------       GGATTGTTTAGGACGTAGATTTACTGTTAATCAGTCATATGTA-----CA
PAH-sensitive male-----------       GGATTGTTTAGGACGTAGATTTACTGTTAATCAGTCATATGTA-----CA
PAH-tolerant female----------       GGAATATTTAAGACGTAGATTTACTGTGAATAAGTCATATGTA-----CA
PAH-tolerant male------------       GGAATATTTAAGACGTAGATTTACTGTGAATAAGTCATATGTACAGTACA
CYP1Arefseq------------------       GGATTGTTTAGGACGTAGATTTACTGTTAATCAGTCATATGTA-----CA
                                    *** * **** **************** *** ***********     **

PAH-sensitive female---------       TACTATATGCAATCTTTTAAACTGAAAGTTATAAATTATGACCTAGACAT
PAH-sensitive male-----------       TACTATATGCAATCTTTTAAACTGAAAGTTATAAATTATGACCTAGACAT
PAH-tolerant female----------       CACTATATGCAATCTTTTAAACTGAAAGTTATAAATTATGACCTAGACAT
PAH-tolerant male------------       TGATATATGCAATCTTTTAA-CTGAAAGTTATAAATTATGACCTAGACAT
CYP1Arefseq------------------       TACTATATGCAATCTTTTAAACTGAAAGTTATAAATTATGACCTAGACAT
                                       ***************** *****************************

PAH-sensitive female---------       ACACACTATTGTGTATATTCTTAAAGTGTGCTAAGATCACCAAAGTGCAG
PAH-sensitive male-----------       ACACACTATTGTGTATATTCTTAAAGTGTGCTAAGATCACCAAAGTGCAG
PAH-tolerant female----------       ACACACTATTGTGTATATTCTTAAAGTGTGCTAAGATCACCAAAGTGCAG
PAH-tolerant male------------       ACACACTATTGTGTATATTCTTAAAGTGTGCTAAGATCACCAAAGTGCAG
CYP1Arefseq------------------       ACACACTATTGTGTATATTCTTAAAGTGTGCTAAGATCACCAAAGTGCAG
                                    **************************************************

PAH-sensitive female---------       CAAGTCTAGATTAAAACGCATGGCGCAGTTCTCACTTACTTGGGACCTCG
PAH-sensitive male-----------       CAAGTCTAGATTAAAACGCATGGCGCAGTTCTCACTTACTTGGGACCTCG
PAH-tolerant female----------       CAAGTCTAGATTAAAACGCATGGCGCAGTTCTCCCTTACTTGGGACCTCG
PAH-tolerant male------------       CAAGTCTAGATTAAAACGCATGGCGCAGTTCTCCCTTACTTGGGACCTCG
CYP1Arefseq------------------       CAAGTCTAGATTAAAACGCATGGCGCAGTTCTCACTTACTTGGGACCTCG
                                    ********************************* ****************

PAH-sensitive female---------       CTGTGTAATCACTATGTTGCACAACACTTCTTTCTTCTCTCTGTCAACAC
PAH-sensitive male-----------       CTGTGTAATCACTATGTTGCACAACACTTCTTTCTTCTCTCTGTCAACAC
PAH-tolerant female----------       CTGTGTAATCACTATGTTGCACAACACTTCTTTCTTCTCTCTGTCAACAC
PAH-tolerant male------------       CTGTGTAATCACTATGTTGCACAACACTTCTTTCTTCTCTCTGTCAACAC
CYP1Arefseq------------------       CTGTGTAATCACTATGTTGCACAACACTTCTTTCTTCTCTCTGTCAACAC
                                    **************************************************

PAH-sensitive female---------       TGATCTAATTCCCTCCTATTTAATTTACAGGTTGAGCAGAGAACAGAGAA
PAH-sensitive male-----------       TGATCTAATTCCCTCCTATTTAATTTACAGGTTGAGCAGAGAACAGAGAA
PAH-tolerant female----------       TGATCTAATTCCCTCCTATTTAATTTACAGGTTGAGCAGAGAACAGAGAA
PAH-tolerant male------------       TGATCTAATTCCCTCCTATTTAATTTACAGGTTGAGCAGAGAACAGAGAA
CYP1Arefseq------------------       TGATCTAATTCCCTCCTATTTAATTTACAGGTTGAGCAGAGAACAGAGAA
                                    **************************************************
```

```
PAH-sensitive female---------        AAGTTGTCATCATGGCATTAATGATACTGCCATTCATTGGAGCACTCTCG
PAH-sensitive male-----------        AAGTTGTCATCATGGCATTAATGATACTGCCATTCATTGGAGCGCTCTCA
PAH-tolerant female----------        AAGTTGTCATCATGGCATTAATGATACTGCCATTCATTGGAGCGCTCTCA
PAH-tolerant male------------        AAGTTGTCATCATGGCATTAATGATACTGCCATTCATTGGAGCGCTCTCA
CYP1Arefseq------------------        AAGTTGTCATCATGGCATTAATGATACTGCCATTCATTGGAGCACTCTCG
                                     ******************************************** *****

PAH-sensitive female---------        GTGTCTGAGGGTTTGATAGCCTTGGTGACGGTGTGCTTGGTCTACCTGAC
PAH-sensitive male-----------        GTGTCTGAGGGTTTGATAGCCTTGGTGACGGTGTGCTTGGTCTACCTGAC
PAH-tolerant female----------        GTGTCTGAGGGTTTGATAGCCTTGGTGACGGTGTGCTTGGTCTACCTGAC
PAH-tolerant male------------        GTGTCTGAGGGTTTGATAGCCTTGGTGACGGTGTGCTTGGTCTACCTGAC
CYP1Arefseq------------------        GTGTCTGAGGGTTTGATAGCCTTGGTGACGGTGTGCTTGGTCTACCTGAC
                                     **************************************************

PAH-sensitive female---------        CCTTAAGCATTTCCGCAGAGAGATCCCAGAGGGGCTACGTCGACTCCCCG
PAH-sensitive male-----------        CCTTAAGCATTTCCGCAGAGAGATCCCAGAGGGGCTACGTCGACTCCCCG
PAH-tolerant female----------        CCTTAAGCATTTCCGCAGAGAGATCCCAGAGGGGCTACGTCGACTCCCCG
PAH-tolerant male------------        CCTTAAGCATTTCCGCAGAGAGATCCCAGAGGGGCTACGTCGACTCCCCG
CYP1Arefseq------------------        CCTTAAGCATTTCCGCAGAGAGATCCCAGAGGGGCTACGTCGACTCCCCG
                                     **************************************************

PAH-sensitive female---------        GCCCGACGCCACTTCCTATCATTGGGAATTTTCTGGAGTTGGGAAGCAAG
PAH-sensitive male-----------        GCCCGATGCCACTTCCTATCATTGGGAATTTTCTGGAGTTGGGAAGCAAG
PAH-tolerant female----------        GCCCGATGCCACTTCCTATCATTGGGAATTTTCTGGAGTTGGGAAGCAAG
PAH-tolerant male------------        GCCCGATGCCACTTCCTATCATTGGGAATTTTCTGGAGTTGGGAAGCAAG
CYP1Arefseq------------------        GCCCGATGCCACTTCCTATCATTGGGAATTTTCTGGAGTTGGGAAGCAAG
                                     ****** *******************************************

PAH-sensitive female---------        CCTTATCTAAGTCTTACTGAAATGAGCAAGCGCTTCGGCGATGTGTTCCA
PAH-sensitive male-----------        CCTTATCTAAGTCTTACTGAAATGAGCAAGCGCTTCGGAGACGTGTTCCA
PAH-tolerant female----------        CCTTATCTAAGTCTTACTGAAATGAGCAAGCGCTTCGGAGACGTGTTCCA
PAH-tolerant male------------        CCTTATCTAAGTCTTACTGAAATGAGCAAGCGCTTCGGAGACGTGTTCCA
CYP1Arefseq------------------        CCTTATCTAAGTCTTACTGAAATGAGCAAGCGCTTCGGAGACGTGTTCCA
                                     ************************************** ** ********

PAH-sensitive female---------        GATTCAGCTCGGCATGCGCCCGTGGTCATTCTGAGTGGTTATGAAACCG
PAH-sensitive male-----------        GATTCAGCTCGGCATGCGTCCCGTGGTCATTCTGAGTGGTTATGAAACAG
PAH-tolerant female----------        GATTCAGCTCGGCATGCGTCCCGTGGTCATTCTGAGTGGTTATGAAACAG
PAH-tolerant male------------        GATTCAGCTCGGCATGCGTCCCGTGGTCATTCTGAGTGGTTATGAAACAG
CYP1Arefseq------------------        GATTCAGCTCGGCATGCGTCCCGTGGTCATTCTGAGTGGTTATGAAACAG
                                     ****************** ***************************** *
```

```
PAH-sensitive female---------       TTAAACAAGCTCTCACCAAACAGGGGGACGACTTTGCCGGCAGGCCAGAC
PAH-sensitive male-----------       TTAAACAAGCTCTCACCAAACAGGGGGACGACTTTGCCGGCAGGCCAGAC
PAH-tolerant female----------       TTAAACAAGCTCTCACCAAACAGGGGGACGACTTTGCCGGCAGGCCAGAC
PAH-tolerant male------------       TTAAACAAGCTCTCACCAAACAGGGGGACGACTTTGCCGGCAGGCCAGAC
CYP1Arefseq------------------       TTAAACAAGCTCTCACCAAACAGGGGGACGACTTTGCCGGCAGGCCAGAC
                                    **************************************************

PAH-sensitive female---------       CTGTACAGCTTCCGCTTCATCAATGATGGAAAGAGTCTGGCCTTCAGCAC
PAH-sensitive male-----------       CTGTACAGCTTCCGCTTCATCAATGATGGAAAGAGTCTGGCCTTCAGCAC
PAH-tolerant female----------       CTGTACAGCTTCCGCTTCATCAATGATGGAAAGAGTCTGGCCTTCAGCAC
PAH-tolerant male------------       CTGTACAGCTTCCGCTTCATCAATGATGGAAAGAGTCTGGCCTTCAGCAC
CYP1Arefseq------------------       CTGTACAGCTTCCGCTTCATCAATGATGGAAAGAGTCTGGCCTTCAGCAC
                                    **************************************************

PAH-sensitive female---------       GGACAAAGCTGGCGTTTGGAGGGCTCGCAGAAAGCTTGCCTACAGTGCAC
PAH-sensitive male-----------       GGACAAAGCTGGCGTTTGGCGGGCTCGCAGAAAGCTTGCCTACAGTGCAC
PAH-tolerant female----------       GGACAAAGCTGGCGTTTGGCGGGCTCGCAGAAAGCTTGCCTACAGTGCAC
PAH-tolerant male------------       GGACAAAGCTGGCGTTTGGCGGGCTCGCAGAAAGCTTGCCTACAGTGCAC
CYP1Arefseq------------------       GGACAAAGCTGGCGTTTGGAGGGCTCGCAGAAAGCTTGCCTACAGTGCAC
                                    ******************* ******************************

PAH-sensitive female---------       TGCGTTCTTTTTCCTCCCTGGAGGGAAAGCTCCCAGAGTACTCCTGTGTG
PAH-sensitive male-----------       TGCGTTCTTTTTCCTCCCTGGAGGGAAAGCTCCCAGAGTACTCCTGTGTG
PAH-tolerant female----------       TGCGTTCTTTTTCCTCCCTGGAGGGAAAGCTCCCAGAGTACTCCTGTGTG
PAH-tolerant male------------       TGCGTTCTTTTTCCTCCCTGGAGGGAAAGCTCCCAGAGTACTCCTGTGTG
CYP1Arefseq------------------       TGCGTTCTTTTTCCTCCCTGGAGGGAAAGCTCCCAGAGTACTCCTGTGTG
                                    **************************************************

PAH-sensitive female---------       CTGGAGGAGCACATCTGCAAAGAGACTGAGCATCTGATCAAGGAACTCCA
PAH-sensitive male-----------       CTGGAGGAGCACATCTGCAAAGAGACTGAGCATCTGATCAAGGAACTCCA
PAH-tolerant female----------       CTGGAGGAGCACATCTGCAAAGAGACTGAGCATCTGATCAAGGAACTCCA
PAH-tolerant male------------       CTGGAGGAGCACATCTGCAAAGAGACTGAGCATCTGATCAAGGAACTCCA
CYP1Arefseq------------------       CTGGAGGAGCACATCTGCAAAGAGACTGAGCATCTGATCAAGGAACTCCA
                                    **************************************************

PAH-sensitive female---------       TAATGTCATGACAGCGGAAGGCAAATTTGACCCTTTCCGCTACATAGTTG
PAH-sensitive male-----------       TAATGTCATGACAGCGGAAGGCAAATTTGACCCTTTCCGCTACATAGTTG
PAH-tolerant female----------       TAATGTCATGACAGCGGAAGGCAAATTTGACCCTTTCCGCTACATAGTTG
PAH-tolerant male------------       TAATGTCATGACAGCGGAAGGCAAATTTGACCCTTTCCGCTACATAGGTG
CYP1Arefseq------------------       TAATGTCATGACAGCGGAAGGCAAATTTGACCCTTTCCGCTACATAGTTG
                                    *********************************************** **
```

```
PAH-sensitive female---------        TTTCTGTTGCCAATGTGATCTGTGGCATGTGCTTTGGCCGACGTTATGAC
PAH-sensitive male-----------        TTTCTGTTGCCAATGTGATCTGTGGCATGTGCTTTGGCCGACGTTATGAC
PAH-tolerant female----------        TTTCTGTTGCCAATGTGATCTGTGGCATGTGCTTTGGCCGACGTTATGAC
PAH-tolerant male------------        TTTCTGTTGCCAATGGGATCTGTGGCATGTGCTTTGGCCGACGTTAGGAC
CYP1Arefseq------------------        TTTCTGTTGCCAATGTGATCTGTGGCATGTGCTTTGGCCGACGTTATGAC
                                     *************** ****************************** ***

PAH-sensitive female---------        CATCATAACCAGGAGCTGCTGAGCTTGGTAAACCTCGCCGAAGATTTTGT
PAH-sensitive male-----------        CATCATAACCAGGAGCTGCTGAGCTTGGTAAACCTCGCCGAAGATTTTGT
PAH-tolerant female----------        CATCATAACCAGGAGCTGCTGAGCTTGGTAAACCTCGCCGAAGATTTTGT
PAH-tolerant male------------        CATCATAACCAGGAGCTGCTGAGCTTGGCAAACCTCGCCGAAGATTTTGT
CYP1Arefseq------------------        CATCATAACCAGGAGCTGCTGAGCTTGGTAAACCTCGCCGAAGATTTTGT
                                     **************************** *********************

PAH-sensitive female---------        CCAGGTGACAGGCAGCGGCAACCCAGCAGATTTTATCCCTGCTCTGCAGT
PAH-sensitive male-----------        CCAGGTGACAGGCAGCGGCAACCCAGCAGATTTCATCCCTGCTCTGCAGT
PAH-tolerant female----------        CCAGGTGACAGGCAGCGGCAACCCAGCAGATTTCATCCCTGCTCTGCAGT
PAH-tolerant male------------        CCAGGGGACAGGCAGCGGCAACCCAGCAGATTTCATCCCTGCTCTGCAGT
CYP1Arefseq------------------        CCAGGTGACAGGCAGCGGCAACCCAGCAGATTTCATCCCTGCTCTGCAGT
                                     ***** *************************** ****************

PAH-sensitive female---------        TTCTGCCCAACAAGTCAATGAAGAAGTTT-GTCAACCTCAA-CAACCGCT
PAH-sensitive male-----------        TTCTGCCCAACAGGTCAATGAAGAAGTTTCGTCAACCTCGA-CAACCGCT
PAH-tolerant female----------        TTCTGCCCAACAAGTCAATGAAGAAGTTT-GTCAACCTCAAACAACCGCT
PAH-tolerant male------------        TTCTGCCCAACAAGCCAATGAAGAAGTTT-GTCAACCTCAA-CAACCGCT
CYP1Arefseq------------------        TTCTGCCCAACAAGTCAATGAAGAAGTTT-GTCAACCTCAA-CAACCGCT
                                     ************ * ************** ********* * ********

PAH-sensitive female---------        TCAACAACTTTGTTCAGAAGATCGTCAGTGAGCACTACTCCACCTTTGAC
PAH-sensitive male-----------        TCAACAACTTTGTTCAGAAGATCGTCAGTGAGCACTACTCCACCTTTGAC
PAH-tolerant female----------        TCAACAACTTTGTTCAGAAGATCGTCAGTGAGCACTACTCCACCTTTGAC
PAH-tolerant male------------        TCAACAACTTTGTTCAGAAGATCGTCAGTGAGCACTACTCCACCTTTGAC
CYP1Arefseq------------------        TCAACAACTTTGTTCAGAAGATCGTCAGTGAGCACTACTCCACCTTTGAC
                                     **************************************************

PAH-sensitive female---------        AAGGTACCCCACTCACTGTACAAGCTATGAAACACGATATAGTCCAATTT
PAH-sensitive male-----------        AAGGTACCCCACTCACTGTACAAGCTATGAAACACGATATAGTCCAATCT
PAH-tolerant female----------        AAGGTACCCCAGTCACTGTACAAGCTATGAAACACGATATAGGCCAATTT
PAH-tolerant male------------        AAGGTACCCCACTCACTGTACAAGCTATGAAACACGATATAGTCCAATTT
CYP1Arefseq------------------        AAGGTACCCCACTCACTGTAAAAGCTATGAAACACGATATAGTCCAATCT
                                     *********** ******** ********************* ***** *
```

```
PAH-sensitive female---------     TTTTTTTCT----CTTTTAAGACGTAGTTTATTTAACCAAAATGTCAGAA
PAH-sensitive male-----------     TTTTTTTTTT-TNNNNNNNNNNNNNAGTTTATTTAACCAAAATGGCAAAA
PAH-tolerant female----------     TTTTTTTTTT-CACTTTAAACACGAAGTTTATTTAACCAAAAGGTCAAAA
PAH-tolerant male------------     TTTTTTTTTT-CACTTTTAAGACAAATTTAATTTAACCCAATGGTCAAAA
CYP1Arefseq------------------     TTTTTTTTTTTCACTTTCAAGACGTAGTTTATTTAACCAAAATGTCAAAA
                                  ******* *               * ** ******** **  * ** **

PAH-sensitive female---------     TGTAATTACTTCCAGGACAACATCCGCGACATCACAGACTCCTTA-ATAG
PAH-sensitive male-----------     TTTAATTACTTCCAGGACAACATCCGAGACATCACAGACTCCTTA-ATAG
PAH-tolerant female----------     TTTAATCTTTTCCAGGATAACATCCGCTACATCACGGACTCCTTATATAG
PAH-tolerant male------------     TTTAATCTTTTCCAGGACAATAACCCTTACATCCCAGATCCCTTA-ATAG
CYP1Arefseq------------------     TTTAATTACTTCCAGGACAACATCCGCGACATCACAGACTCCTTA-ATAG
                                  * ****   ******** ** * **    ***** * **  ***** ****

PAH-sensitive female---------     ATCACTGCGAGGACCGGAAGCTCGATGAGAACTCCAACATCCAGATGTCA
PAH-sensitive male-----------     ATCACTGCGAGGACCGGAAGCTCGATGAGAACTCCAACATCCAGATGTCA
PAH-tolerant female----------     ATCACTGCGAGGACCGGAAGCTCGATGAGAACTCCAACATCCAGATGTCA
PAH-tolerant male------------     ATCACTGCGAGGACCGGAAGCTCGATGAGAACTCCAACATCCAGATGTCA
CYP1Arefseq------------------     ATCACTGCGAGGACCGGAAGCTCGATGAGAACTCCAACATCCAGATGTCA
                                  **************************************************

PAH-sensitive female---------     GATGAGAAAATTGTTGGCATTGTCAACGACCTCTTTGGAGCTGGTAGGTT
PAH-sensitive male-----------     GATGAGAAAATTGTTGGCATTGTCAACGACCTCTTTGGAGCTGGTAGGTT
PAH-tolerant female----------     GATGAGAAAATTGTTGGCATTGTCAACGACCTCTTTGGAGCTGGTAGGTT
PAH-tolerant male------------     GATGAGAAAATTGTTGGCATTGTCAACGACCTCTTTGGAGCTGGTAGGTT
CYP1Arefseq------------------     GATGAGAAAATTGTTGGCATTGTCAACGACCTCTTCGGAGCTGGTAGGTT
                                  *********************************** **************

PAH-sensitive female---------     TTTATGCATGCTGAGTGCTTGGATCATGTGTAATGATTATTTTACACACC
PAH-sensitive male-----------     TTTATGCGTGCTGAGTGCTTGGATCAAATGTAATCATTATTTTACACAAC
PAH-tolerant female----------     TTTATGCGTGCTGAGTGCTTGGATCAAATGTAATCATTATTTTACACACC
PAH-tolerant male------------     TTTATGCGTGCTGAGTGCTTGGATCAAATGTAATCATTATTTTACACACC
CYP1Arefseq------------------     TTTATGCGTGCTGAGTGCTTGGATCATGTGTAATGATTATTTTACACACC
                                  ******* ******************  ****** ************* *

PAH-sensitive female---------     GCAGGCTTCCTCACAATTCTGCTTTTGATCCTTTTAGGTTTCGACACCAT
PAH-sensitive male-----------     GCAGGCTTCCTCACAATTCTGCTTTTGATCCTTTTAGGTTTCGACACCAT
PAH-tolerant female----------     GCAGGCTTCCTCACAATTCTGCTTTTGATCCTCTTAGGTTTCGACACCAT
PAH-tolerant male------------     GCAGGCTTCCTCACAATTCTGCTTTTGATCCTCTTAGGTTTCGACACCAT
CYP1Arefseq------------------     GCAGGCTTCCTCACAATTCTGCTTTTGATCCTTTTAGGTTTCGACACCAT
                                  ******************************** *****************
```

```
PAH-sensitive female---------     CTCTACTGCTTTGTCATGGGCAGTGATGTACCTTGTGGCTTATCCAGAGG
PAH-sensitive male-----------     CTCTACCGCTTTGTCATGGGCAGTGATGTACCTTGTGGCTTATCCAGAG-
PAH-tolerant female----------     CTCTACCGCTTTGTCATGGGCGGTGATGTACCTTGTGGCTTATCCAGAGG
PAH-tolerant male------------     CTCTACCGCTTTGTCATGGGCGGTGATGTACCTTGTGGCTTATCCAGAGG
CYP1Arefseq------------------     CTCTACCGCTTTGTCATGGGCAGTGATGTACCTTGTGGCTTATCCAGAGG
                                  ****** ************** ***************************

PAH-sensitive female---------     TTGAGGAGAGGCTTTATGAAGAAATCAGTAAGTTAGAGCTTTTTACTGAA
PAH-sensitive male-----------     CTGAGGAGAGGCTTTATGAAGAAATCAGTAAGTTAGAGCTTTTTACTGAA
PAH-tolerant female----------     TTGAGGAAAGGCTTTATGAAGAAATCAGTAAGTTAGAGCTTTTTACTGAA
PAH-tolerant male------------     TTGAGGAAAGGCTTTATGAAGAAATCAGTAAGTTAGAGCTTTTTACTGAA
CYP1Arefseq------------------     TTGAGGAGAGGCTTTATGAAGAAATCAGTAAGTTAGAGCTTTTTACTGAA
                                   ****** ******************************************

PAH-sensitive female---------     ATAGTTCTTTGACTATTTTATCTTGCATGGAAATAAATATTCCATGAAAA
PAH-sensitive male-----------     ATAGTTCTTTGACTATTTTATCTTGCATGGGAATAAATATTCCATAAAAA
PAH-tolerant female----------     ATAGTTCTTTGACTATTTTATCTTGCATGGAAATATATATTCAATAAAAA
PAH-tolerant male------------     ATAGTTCTTTGACTATTTTATCTTGCATGGAAATATATATTCAATAAAAA
CYP1Arefseq------------------     ATAGTTCTTTGACTATTTTATCTTGCATGGAAATAAATATTCCATGAAAA
                                  ****************************** **** ****** ** ****

PAH-sensitive female---------     AATATTTTATGAATATT—GGTAATAAAAATTCTTACTTCAGACTCATAA
PAH-sensitive male-----------     CATATTTCTTGAATATT--GCCAATAAAAAGGCTTGCTTCAAACTCATAC
PAH-tolerant female----------     -ATATTTCATTAATATTTTGGTAATAAAAACACTTGCTATAGACTCATAA
PAH-tolerant male------------     -ATATTTCATTAATATTTTGGTAATAAAAACACTTGCTATAGACTCATAA
CYP1Arefseq------------------     AATATTTTATGAATATT--GGTAATAAAAATTCTTGCTTCAGACTCATAA
                                   ******  * ******  *  ********  *** **  * *******

PAH-sensitive female---------     AGTAGAATTTCTTTTCTGCTATTAAGAAAACATTATTTTCTTATTTAACC
PAH-sensitive male-----------     TGTAGAATTTCTTTTCTGCTATTAAGAAAACATTATTTTCTTATTTAACC
PAH-tolerant female----------     AGTAGAATTTATTTTCTGCCATTAAGAAAACAGGATTTTCTCATTTAACC
PAH-tolerant male------------     AGTAGAATTTATTTTCTGCCATTAAGAAAACATAATTTTCTCATTTAACC
CYP1Arefseq------------------     AGTAGAATTTCTTTTCTGCTATTAAGAAAACATTATTTTCTTATTTAACC
                                   ********* ******** ************  ******* ********

PAH-sensitive female---------     AACTCTTCTCATTTTTAGAGGAGAAAGTCGGTCTGGATCGTACTCCTGTT
PAH-sensitive male-----------     AACTCTTCTCATTTTTAGAGGAGAAAGTCGGTCTGGATCGTACTCCTGTT
PAH-tolerant female----------     AACTCTTCTCATTTTTAGAGGAGAAAGTCGGTCTGGATCGTACTCCTGTT
PAH-tolerant male------------     AACTCTTCTCATTTTTAGAGGAGAAAGTCGGTCTGGATCGTACTCCTGTT
CYP1Arefseq------------------     AACTCTTCTCATTTTTAGAGGAGAAAGTCGGTCTGGATCGTACTCCTGTT
                                  **************************************************
```

```
PAH-sensitive female---------       ATGTCTGACAGAAGCAACTTGCCTTTACTTGAGTCTTTCATTCTGGAGCT
PAH-sensitive male-----------       ATGTCTGACAGAAGCAACTTGCCTTTACTTGAGTCTTTCATTCTGGAGCT
PAH-tolerant female----------       ATGTCTGACAGAAGCAACTTGCCTTTACTTGAGTCTTTCATTCTGGAGCT
PAH-tolerant male------------       ATGTCTGACAGAAGCAACTTGCCTTTACTTGAGTCTTTCATTCTGGAGCT
CYP1Arefseq------------------       ATGTCTGACAGAAGCAACTTGCCTTTACTTGAGTCTTTCATTCTGGAGCT
                                    **************************************************

PAH-sensitive female---------       CTTTCGTCATTCTTCATACCTGCCTTTCACAATCCCACACTGGTAAGATT
PAH-sensitive male-----------       CTTTCGTCATTCTTCATACCTGCCTTTCACAATCCCACACTGGTAAGATT
PAH-tolerant female----------       CTTTCGTCACTCTTCATACCTGCCTTTCACAATCCCACACTGGTAAGATT
PAH-tolerant male------------       CTTTCGTCACTCTTCATACCTGCCTTTCACAATCCCACACTGGTAAGATT
CYP1Arefseq------------------       CTTTCGTCATTCTTCATACCTGCCTTTCACAATCCCACACTGGTAAGATT
                                    ********* ****************************************

PAH-sensitive female---------       GGTCTCTTCTACTCTTTGCTTACAATAGTTTGTTGATAATGACATGAAGA
PAH-sensitive male-----------       GGTCTCTTCTACTCTTTGCTTACAATAGTTTGTTGACAATGACATGCAGA
PAH-tolerant female----------       GGTCTCTTCTACTCTTTGCTTACAAAAGTTTGTTGACAATGACATGCAGA
PAH-tolerant male------------       GGTCTCTTCTACTCTTTGCTTACAAAAGTTTGTTGACAATGACATGCAGA
CYP1Arefseq------------------       GGTCTCTTCTACTCTTTGCTTACGAAAGTTTGTTGATAATGACATGAAGA
                                    *********************** * ********** ********* ***

PAH-sensitive female---------       GTATGTCACAAAGGCTAAAAAAGATAAACAACGAGAAGGGATGCTCAATG
PAH-sensitive male-----------       GTATGTCACAAAGGCTAAAAAAGATAAACAATGAGAAGGGATGGCCAATG
PAH-tolerant female----------       GTATGTCACAAAGGCTAAAAAAGATAAACAACGAGAAGGGATGCCCAATG
PAH-tolerant male------------       GTATGTCACAAAGGCTAAAAAAGATAAACAACGAGAAGGGATGCCCAATG
CYP1Arefseq------------------       GTATGTCACAAAGGCTAAAAAAGATAAACAACGAGAAGGGATGGCCAATG
                                    ******************************* ***********  *****

PAH-sensitive female---------       GGCTGACATTTATTTTTTGCTTTTTTTTT-AGCTCTACAAAAGATACATC
PAH-sensitive male-----------       TGCTTACATTTAATTTT-GCTTTTTT--C-AGCTCTACAAAAGATACATC
PAH-tolerant female----------       GGCTGACATTTATTTTT-GCTTTTTTTTT-AGCTCTACAAAAGATACATC
PAH-tolerant male------------       GGCTGACATTTATTTTTTGCTTTTTTATT-AGCTCTACAAAAGATACATC
CYP1Arefseq------------------       GGCTGACATTTATTTTTTGCTTTTTTTTTTAGCTCTACAAAAGATACATC
                                     *** ******* **** ********    ********************

PAH-sensitive female---------       TCTGAACGGCTACTTCATTCTCCAAGACACCTGCGTGTTTGTCAACCAGT
PAH-sensitive male-----------       TCTGAACGGCTACTTCATTCCCAAAGACACCTGCGTGTTTGTCAACCAGT
PAH-tolerant female----------       TCTGAACGGCTACTTCATTCCCAAAGACACCTGCGTGTTTGTCAACCAGT
PAH-tolerant male------------       TCTGAACAGCTACTTCATTCCCAAAGACACCTGCGTGTTTGTCAACCAGT
CYP1Arefseq------------------       TCTGAACGGCTACTTCATTCCCAAAGACACCTGCGTGTTTGTCAACCAGT
                                    ******* ************ * ***************************
```

```
PAH-sensitive female---------     GGCAGATAAACCACGACCCGTAGGTTTCTTTATAGCGTTTAATGTTTAGT
PAH-sensitive male-----------     GGCAGATAAACCACGACCCGTAAGTTTCTTTATAGCATTTAATGTTTAGT
PAH-tolerant female----------     GGCAGATAAACCACGACCCGTAAGTTTCTTTATAGCGTTCAATGTTTAGT
PAH-tolerant male------------     GGCAGATAAACCACGACCCGTAAGTTTCTTTATAGCGTTTAATGTTTAGT
CYP1Arefseq------------------     GGCAGATAAACCACGACCCGTAAGTTTCTTTATAGCTTTCAATGTTTAGT
                                  ********************** ************* ** **********

PAH-sensitive female---------     TCTCAACTTTGTGCCAATACAGTTGCCGCC--TCACACGACAGACACTCT
PAH-sensitive male-----------     TCTCAATTTTGTGCCAATACAGTTGCCGCC--TCACATGACAGACACTCT
PAH-tolerant female----------     TCTCAACTTTGTGCCAATACAGTTGCCGCC--TCACACGACAGACACTCT
PAH-tolerant male------------     TCTCAACTTTGTGCCAATACAGTTTTCGTGAACGAAACGACAGACACTCT
CYP1Arefseq------------------     TCTCAATTTTGTGCCAATACAGTTGCCGCC--TCACACGACAGACACTCT
                                  ****** *****************  **      * * ************

PAH-sensitive female---------     TTTGCAAGTTGTAAACAAAGAAAACCACAATCACTTAAATTAGGAAACAA
PAH-sensitive male-----------     TTTGCAAGTTGTAAACAAAGAAAACCACAATCATTTAAATTAGGAAAGAA
PAH-tolerant female----------     TTTGCAAGTTGTAAACAAAGAAAACCACAATCACTTAAATTAGGAAACAA
PAH-tolerant male------------     TTTGCAAGTTGTAAACAAAGAAAACCACAATCACTTAAATTAGGAAACAA
CYP1Arefseq------------------     TTTGCAAGTTGTAAACAAAGAAAACCACAATCATTTAAATTAGGAAGCAA
                                  ********************************* ************  **

PAH-sensitive female---------     TAAGGATCGAGCCTGGTTTATTTATTTATTCGTTTTCATTTCTT-GCTGC
PAH-sensitive male-----------     TAAGGATCGAGCCTGATTTATTTATTTATGCGTTTTAATTTTTTTGCTGC
PAH-tolerant female----------     TAAGGATCGAGCCTGGTTTATTTATTTATTCGCTTTCATTTTTT-GCTGC
PAH-tolerant male------------     TAAGAATCGAGCCTGGTTTATTTATTTATTCGCTTTCATTTCTT-GCTGC
CYP1Arefseq------------------     TAAGGATCGAGCCTGGTTTATTTATTTATTCGTTTTCATTTTTTTGCTGC
                                  **** ********** ************* ** *** **** ** *****

PAH-sensitive female---------     CTTTCAGAGAGCTCTGGAAAGACCCGTCTATGTTCATCCCAGACCGCTTC
PAH-sensitive male-----------     CTTTCAGAGAGCTCTGGAAAGACCCGTCTATGTTCATCCCAGACCGCTTC
PAH-tolerant female----------     CTTTCAGAGAGCTCTGGAAAGACCCGTCTATGTTCATCCCAGACCGCTTC
PAH-tolerant male------------     CTTTCAGAGAGCTCTGGAAAGACCCGTCTATGTTCATCCCAGACCGCTTC
CYP1Arefseq------------------     CTTTCAGAGAGCTCTGGAAAGACCCGTCTATGTTCATCCCAGACCGCTTC
                                  **************************************************

PAH-sensitive female---------     CTCAGCGCTGACGGCACAGAGGTAAACAAGCAAGAGGGAGAGAAGGTGCT
PAH-sensitive male-----------     CTCAGCGCTGACGGCACAGAGGTAAACAAGCAAGAGGGAGAGAAGGTGCT
PAH-tolerant female----------     CTCAGCGCTGACGGCACAGAGGTAAACAAGCAAGAGGGAGAGAAGGTGCT
PAH-tolerant male------------     CTCAGCGCTGACGGCACAGAGGTAAACAAGCAAGAGGGAGAGAAGGTGCT
CYP1Arefseq------------------     CTCAGCGCTGACGGCACAGAGGTAAACAAGCAAGAGGGAGAGAAGGTGCT
                                  **************************************************
```

```
PAH-sensitive female---------       AATTTTCGGCTTGGGAAGACGGCGGTGCATCGGTGAGGTTATCGCACGAA
PAH-sensitive male-----------       TATTTTCGGCTTGGGAAGACGGCGGTGCATCGGTGAGGTTATCGCACGAA
PAH-tolerant female----------       TATTTTCGGCTTGGGAAGACGGCGGTGCATCGGTGAGGTTATCGCACGAA
PAH-tolerant male------------       TATTTTCGGCTTGGGAAGACGGCGGTGCATCGGTGAGGTTATCGCACGAA
CYP1Arefseq------------------       TATTTTCGGCTTGGGAAGACGGCGGTGCATCGGTGAGGTTATCGCACGAA
                                     *************************************************

PAH-sensitive female---------       ACGAAGTCTTCCTCTTCCTGGCAATCATCATCCAGAAACTGCACTTTTAC
PAH-sensitive male-----------       ACGAAGTCTTCCTCTTCCTGGCAATCATCATCCAGAAACTGCACTTTTAC
PAH-tolerant female----------       ACGAAGTCTTCCTCTTCCTGGCAATCATCATCCAGAAACTGCACTTTTAC
PAH-tolerant male------------       ACGAAGTCTTCCTCTTCCTGGCAATCATCATCCAGAAACTGCACTTTTAC
CYP1Arefseq------------------       ACGAAGTCTTCCTCTTCCTGGCAATCATCATCCAGAAACTGCACTTTTAC
                                    **************************************************

PAH-sensitive female---------       AAGTTGCCCGGAGAGCCCGTGGACATGACCCCGGAGTATGGCCTCACG-A
PAH-sensitive male-----------       AAGTTGCCCGGAGAGCCCGTGGACATGACCCCGGAGTATGGCCTCACG-A
PAH-tolerant female----------       AAGTTGCCCGGAGAGCCCGTGGACATGACCCCGGAGTATGGCCTCACG-A
PAH-tolerant male------------       AAGTTGCCCGGAGAGCCCGTGGACATGACCCCGGAGTATGGCCTCACGTA
CYP1Arefseq------------------       AAGTTGCCCCGAGAGCCCGTGGACATGACCCCGGAGTATGGCCTCACG-A
                                    ********* ************************************** *

PAH-sensitive female---------       TGAAGCACAAACGCTGTTACCTGGGAGTCGCAATGAGAGCTAAGGACGTG
PAH-sensitive male-----------       TGAAGCACAAACGCTGTTACCTGGGAGTCGCAATGAGAGCTAAGGACGTG
PAH-tolerant female----------       TGAAGCACAAACGCTGTTACCTGGGAGTCGCAATGAGAGCTAAGGACGTG
PAH-tolerant male------------       TGAAGCACAAACGCTGTTACCTGGGAGTCGCAATGAGAGCTAAGGACGTG
CYP1Arefseq------------------       TGAAGCACAAACGCTGTTACCTGGGAGTCGCAATGAGAGCTAAGGACGTG
                                    **************************************************

PAH-sensitive female---------       CAGTGAAGCTCTGCGTTATTTAGAATGTAAGACTTTGAAGGTGGCCTATG
PAH-sensitive male-----------       CAGTGAAGCTCTGCATTATTTAGAATGTAAGACTTTGAAGGTGGCCTGTG
PAH-tolerant female----------       CAGTGAAGCTCTGCATTATTTAGAATGTAAGACTTTGAAGGTGGCCTATG
PAH-tolerant male------------       CAGTGAAGCTCTGCATTATTTAGAATGTAAGACTTTGAAGGTGGCCTATG
CYP1Arefseq------------------       CAGTGAAGCTCTGCGTTATTTAGAATGTAAGACTTTGAAGGTGGCCTATG
                                    ************** ******************************** **

PAH-sensitive female---------       TTGACTGTGACACTGTAGATTAAGAGTTACATTAAGTTTGAGTGAAATAA
PAH-sensitive male-----------       TTGACTGTGACACTGTAGATTAAGAGTTACATTAAGTTTGAGTGAAATAA
PAH-tolerant female----------       TTGACTGTGACACTGTAGATTAAGAGTTACATTAAGTTTGAGTGAAATAA
PAH-tolerant male------------       TTGACTGTGACACTGTAGATTAAGAGTTACATTAAGTTTGAGTGAAATAA
CYP1Arefseq------------------       TTGACTGTGACACTGTAAATTAAGAGTTACATTAAGTTTGAGTGAAATAA
                                    ***************** ********************************
```

```
PAH-sensitive female---------       GTATATTTCTGAGAATGGAGGACACTGGATGACCTTAATTTATAGAGCTA
PAH-sensitive male-----------       GTATATTTCTGAGAATGGAGGACACTGGATGACCTTAATTTATAGAGCTA
PAH-tolerant female----------       GTATATTTCTAAGAATGGAGGACACTGGATGACCTTAATTTATAGAGCTA
PAH-tolerant male------------       GTATATTTCTAAGAATGGAGGACACTGGATGACCTTAATTTATAGAGCTA
CYP1Arefseq------------------       GTATATTTCTGAGAATGGAGGACACTGGATGACCTTAATTTATAGAGCTA
                                    ********** ***************************************

PAH-sensitive female---------       ACGACATTGAGGCAAATCCAATGAAATTTGCTTGATGCAGACTGCAAGAC
PAH-sensitive male-----------       ACGGCATTGAGGCAAATCCAA-GAAATTTGCTTGATGCAGACTGCAAGAC
PAH-tolerant female----------       ACAGCATTGAGGCAAATCCAA-GAAATTTGCTTGATGCAGACTGCAAGAC
PAH-tolerant male------------       ACAGCATTGAGGCAAATCCAA-GAAATTTGCTTGATGCAGACTGCAAGAC
CYP1Arefseq------------------       ACGGCATTGAGGCAAATCCAA-GAAATTTGCTTGATGCAGACTGCAAGAC
                                    **  ***************** ****************************

PAH-sensitive female---------       ATGTCTACATTTAGGTTC-ATTGCTTTGTAGTTTGAGGCTCTGCAAAGTA
PAH-sensitive male-----------       ATGTCTACATTTAGCCCCCATTGCTTTGTAGTTTGAGGCTCTGCAAAGTA
PAH-tolerant female----------       ATGTCTACATTTAGGTTC-ATTGCTTTGTAGTTTGAGGCTCTGCAAAGTA
PAH-tolerant male------------       ATGTCTACATTTAGGTTC-ATTGCTTTGTAGTTTGAGGCTCTGCAAAGTA
CYP1Arefseq------------------       ATGTCTACATTTAGGTTC-ATTGCTTTGTAGTTTGAGGCTCTGCAAAGTA
                                    **************   * *******************************

PAH-sensitive female---------       ACATTCCTACACGTTGTCTCTGCTTCTAAATGAAGAACAAAAAGGAGGAT
PAH-sensitive male-----------       ACATTCCTACACGTTGTCTCTGCTTCTAAATGAAGAACAAAAAGGAGGAT
PAH-tolerant female----------       ACATTCCTACACGTTGTCTCTGCTTCTAAATGAAGAACAAAAAGGTGGAT
PAH-tolerant male------------       ACATTCCTACACGTTGTCTCTGCTTCTAAATGAAGAACAAAAAGGTGGAT
CYP1Arefseq------------------       ACATTCCTACACGTTGTCTCTGCTTCTAAATGAAGAACAAAAAGGTGGAT
                                    ********************************************* ****

PAH-sensitive female---------       GCTTCGCTGTCGTGTTGTCCTGTAATCTAAAAGCAATCCATGCAGAAGTG
PAH-sensitive male-----------       GCTTCGCTGTCGTGTTGTCCTGTAATCTAAAAGCAATCCATGCAGAAGTG
PAH-tolerant female----------       GCTTCGCTGTCGTGTTGTCCTGTAATCTAAAAGCAATCCATGCAGAAGTG
PAH-tolerant male------------       GCTTCGCTGTCGTGTTGTCCTGTAATCTAAAAGCAATCCATGCAGAAGTG
CYP1Arefseq------------------       GCTTCGCTGTCGTGCTGTCCTGTAATCTAAAAGCAATCCATGCAGAAGTG
                                    ************** ***********************************

PAH-sensitive female---------       ACTAAGCAGACAAGAGTTTTTGGATTGAGAGCAAATTTTATAGGCATTAC
PAH-sensitive male-----------       ACTAAGCAGACAAGAGTTTTTGGATTGAGAGTAAATTTTATAGGCATTAC
PAH-tolerant female----------       TCTAAGCAGACAAGAGTTTTTGGATTGAGAGTAAATTTTATAGGCATTAC
PAH-tolerant male------------       TCTAAGCAGACAAGAGTTTTTGGATTGAGAGTAAATTTTATAGGCATTAC
CYP1Arefseq------------------       ACTAAGCAGACAAGAGTTTTTGGATTGAGAGTAAATTTTATAGGCATTAC
                                     ****************************** ******************
```

```
PAH-sensitive female---------        CATGAAGGGCACTGCAATGTGTGCATTGTAACAGAAAACTCT-GGAGCAA
PAH-sensitive male-----------        CATGAAGGGCACTGCAATGTGTGCATTGTAACAGATTACTCT-GGAGCAA
PAH-tolerant female----------        CATGAAGGGCACTGCAATGTGTGCATTGTAACAGAAAACTCTTGGATCAA
PAH-tolerant male------------        CATGAAGGGCACTGCAATGTGTGCATTGTAACAGAAAACTCT-GGAGCAA
PAH-sensitive female---------        CATGAAGGGCACTGCAATGTGTGCATTGTAACAGAAAACTCT-GGAGCAA
CYP1Arefseq------------------        CATGAAGGGCACTGCAATGTGTGCATTGTAACAGAAAACTCT-GGAGCAA
                                     ***********************************  ***** *** ***

PAH-sensitive female---------        ACAGATATTACAGTATGACACCATTCACG-TTTGTAAGCTAATTATTTTT
PAH-sensitive male-----------        ACAGATATTACAGTATGACACCATTCACG-TTTGTAAGCTAATTATTTGT
PAH-tolerant female----------        ACAGATATTACAGTATGACACCATTCACGGTTTGGAAGCTAATTATTTTT
PAH-tolerant male------------        ACAGATATTACAGTATGACACCATTCACG-TTTGTAAGCTAATTATTTTT
CYP1Arefseq------------------        ACAGATATTACAGTATGACACCATTCACG-TTTGTAAGCTAATTATTTTT
                                     ***************************** **** ************* *

PAH-sensitive female---------        CATTTCATTATACTGTAAATGCCAATCCTGAAGCTATATTTGTATCCCTA
PAH-sensitive male-----------        CATTTCATTATACTGTAAATGCCAATCCTGAAGCTATATTTGTATCCCTA
PAH-tolerant female----------        CATTTCATTATACTGTAAATGCCAATCCTGAAGCTATATTTGTATCCCTA
PAH-tolerant male------------        CATTTCATTATACTGTAAATGCCAATCCTGAAGCTATATTTGTATCCCTA
CYP1Arefseq------------------        CATTTCATTATACTGTAAATGCCAATCCTGAAGCTATATTTGTATCCCTA
                                     **************************************************

PAH-sensitive female---------        AATGTGATTATTTGTGTATGTGTGTCTGTATTTTCTTATAGGTCATTTTT
PAH-sensitive male-----------        AATGTGATTATTTGTGTATGTGTGTCTGTATTTTCTTATAGGTCATTTTT
PAH-tolerant female----------        AATGTGATTATTTGTGTATGTGTGTCTGTATTTTCTTATAGGTCTTTTTT
PAH-tolerant male------------        AATGTGATTATTTGTGTATGTGTGTCTGTATTTTCTTATAGGTCTTTTTT
CYP1Arefseq------------------        AATGTGATTATTTGTGTATGTGTGTCTGTATTTTCTTATAGGTCATTTTT
                                     ******************************************** *****

PAH-sensitive female---------        TTT-ATGAACCAAGGCTTTTGGTTATGTTTGAAGGGAACCACTTATCTGT
PAH-sensitive male-----------        TTTTATGATCCAAGCATTTTGTTTATGTTTGAGGTGAACTACTTATCTGT
PAH-tolerant female----------        TTTTATGATCCAAGCATTTTGTTTATGTTTGAGGTGAACTACTTATCTGT
PAH-tolerant male------------        TTTTATGATCCAAGCATTTTGTTTATGTTTGAGGTGAACTACTTATCTGT
CYP1Arefseq------------------        TTTTATGATCCAAGCATTTTGTTTATGTTTGAGGTGAACTACTTATCTGT
                                     *** **** *****  ***** ********** * **** **********

PAH-sensitive female---------        GCCCAAAAATGGTTTTTTCGGGCAGAACCAATTATTTCGGAAAAATTAAC
PAH-sensitive male-----------        GCCAAAAAAATGATTTATCAGGGCAGATCAAATGATTCTGGAAAAATTATC
PAH-tolerant female----------        GCCAAAAAAATGATTTATCAGGGCAGATCAAATGATTCTGGAAAAATTATC
PAH-tolerant male------------        GCCAAAAAAATGATTTATCAGGGCAGATCAAATGATTCTGGAAAAATTATC
CYP1Arefseq------------------        GCCAAAAAAATGATTTATCAGGGCAGATCAAATGATTCTGGAAAAAATATC
                                     *** ******* *** *  ******* * *** ***  ******* ** *
```

```
PAH-sensitive female---------       ACCATGCTGTACCAATTTAAATAACCTAAGGAATAATTCTTGTAAAAGGA
PAH-sensitive male-----------       ACAATGCTGTACAATTTTATATATCTTATGTATCATTCATTGTAAAAG-A
PAH-tolerant female----------       ACAATGCTGTACAATTTTATATATCTTATGTATCATTCATTGTAAAAG-A
PAH-tolerant male------------       ACAATGCTGTACAATTTTATATATCTTATGTATCATTCATTGTAAAAG-A
CYP1Arefseq------------------       ACAATGCTGTACAATTTTATATATCTTATGTATCATTCATTGTAAAAGAA
                                    ** ********* * **** *** * ** * *  * *  ********* *

PAH-sensitive female---------       AAAAAATGTCAGGGTTTGAGTTATTTTATGAACAGAAATAAAATTGAATG|
PAH-sensitive male-----------       AAAAAATGTCAGGGTTTGAGTTTTTTTATGAACCGAAATAAAATTGAATG|
PAH-tolerant female----------       AAAAAATGTCAGGGTTTGAGTTATTTTATGAACAGAAATAAAATTGAATG|
PAH-tolerant male------------       AAAAAATGTCAGGGTTTGAGTTATTTTATGAACAGAAATAAAATTGAATG|
CYP1Arefseq------------------       AAAAAATGTCAGGGTTTGAGTTATTTTATGAACAGAAATAAAATTGAATG|
                                    ********************** *********** ***************
```

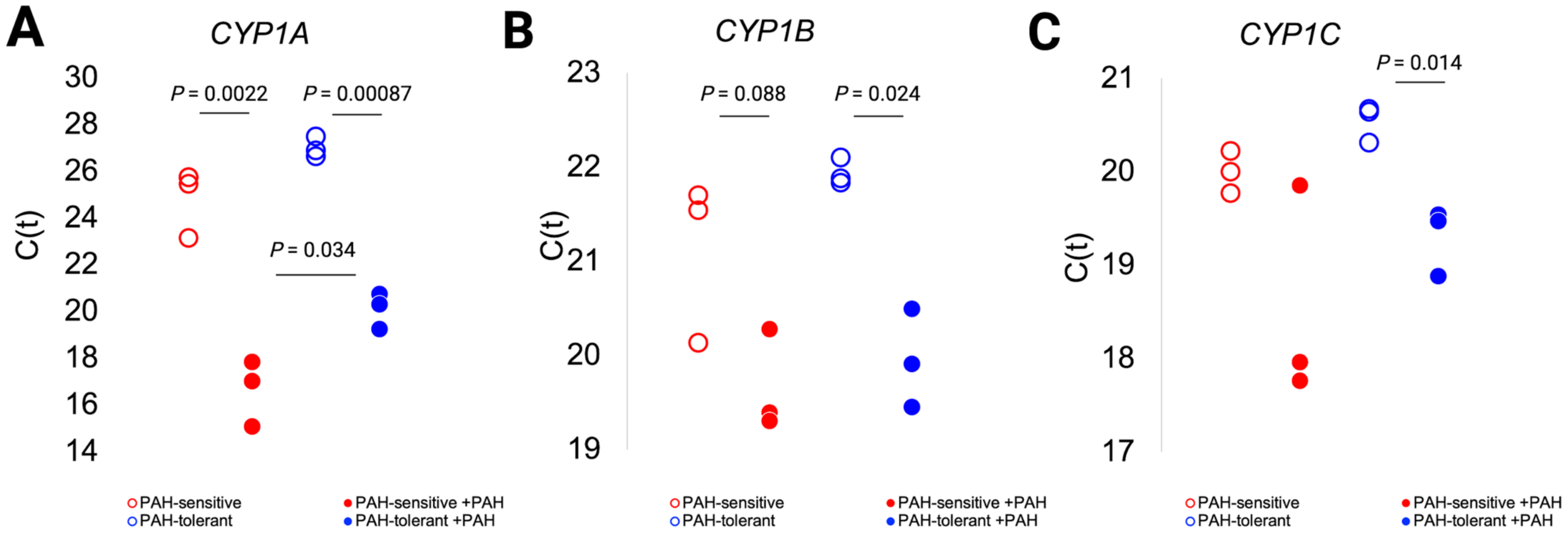


**Supplemental Figure 2. Expression of AhR target genes *CYP1A*, *CYP1B*, and *CYP1C* in PAH-sensitive and PAH-tolerant *F. heteroclitus* embryos, with and without PAH challenge. (a)** Cycles to threshold (C(t)) values for *CYP1A* by population and exposure status. **(b)** C(t) values for *CYP1B* by population and exposure status. **(c)** C(t) values for *CYP1C* by population and exposure status. *P*-values are for independent samples t-tests comparing C(t) values between populations.

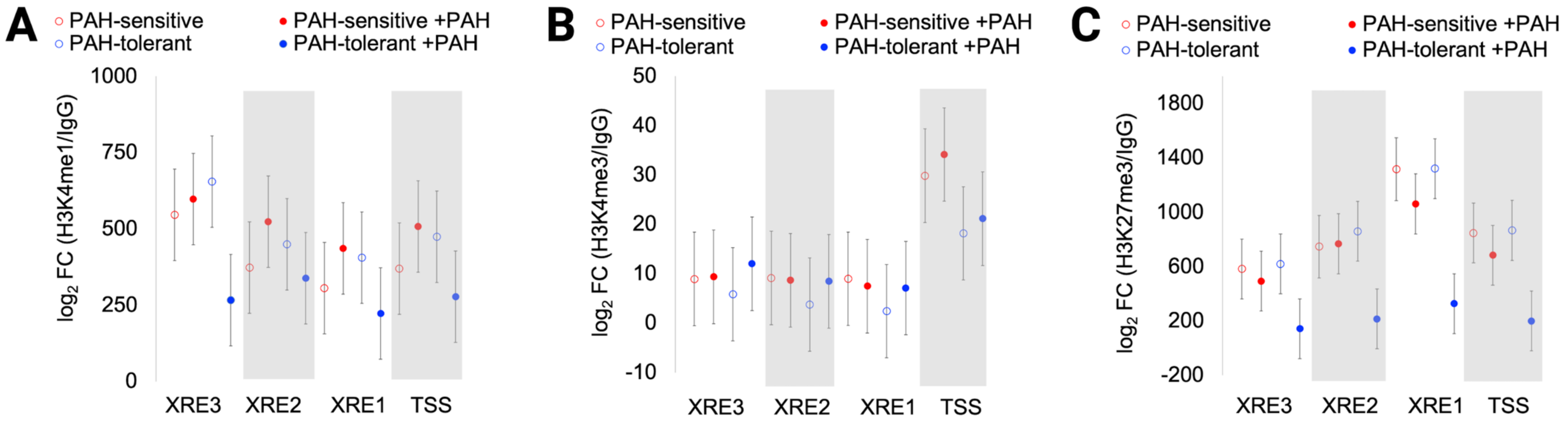


**Supplemental Figure 3. Histone post-translational modifications at the *CYP1A* proximal promoter-enhancer in PAH-sensitive and PAH-tolerant *F. heteroclitus* embryos, with and without PAH challenge. (a)** Least squares means and SEM for histone post-translational modification (PTM) H3K4me1 enrichment in embryos challenged with PAH or control embryos from PAH-tolerant and PAH-sensitive populations. **(b)** Least squares means and SEM for histone PTM H3K4me3 enrichment in embryos challenged with PAH or control embryos from PAH-tolerant and PAH-sensitive populations. **(c)** Least squares means and SEM for histone PTM H3K27me3 enrichment in embryos challenged with PAH or control embryos from PAH-tolerant and PAH-sensitive populations. Data are adjusted for experimental batch and correlation among gene regions within each sample (see Methods for more details). XRE – xenobiotic response element, TSS – transcriptional start site.

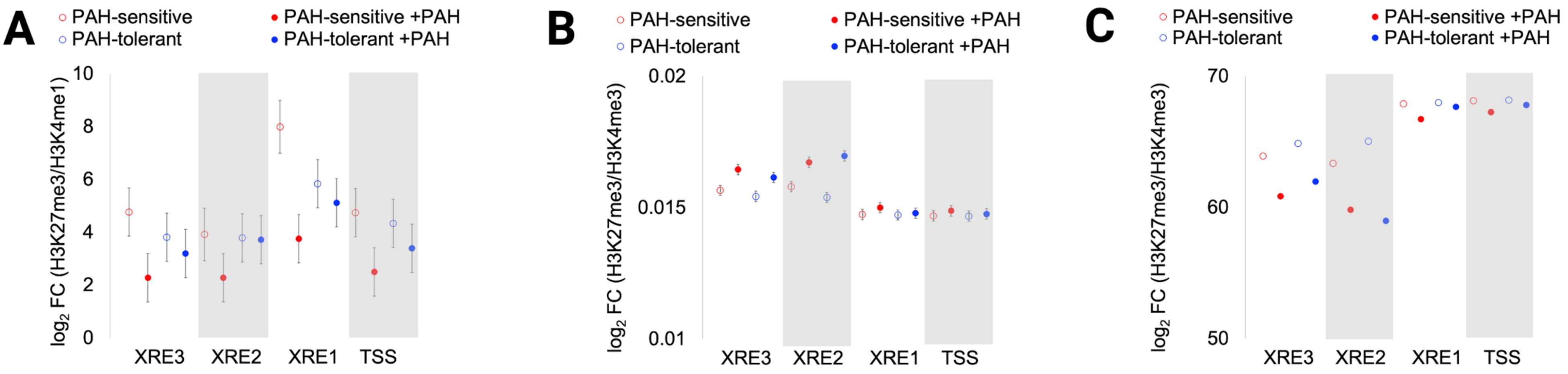


**Supplemental Figure 4. Repressive: activating ratios of histone post-translational modifications at the *CYP1A* proximal promoter-enhancer in PAH-sensitive and PAH-tolerant *F. heteroclitus* embryos, with and without PAH challenge. (a)** Least squares means and SEM for enrichment of histone PTM ratio H3K27me3: H3K4me1 in embryos challenged with PAH or control embryos from PAH-tolerant and PAH-sensitive populations. **(b)** Least squares means and SD (inverse-transformed) for enrichment of histone PTM ratio H3K27me3: H3K4me3 in embryos challenged with PAH or control embryos from PAH-tolerant and PAH-sensitive populations. **(c)** Least squares means (back-transformed) for enrichment of histone PTM ratio H3K27me3: H3K4me3 in embryos challenged with PAH or control embryos from PAH-tolerant and PAH-sensitive populations. Original models required inverse transformation of skewed data. Data are adjusted for experimental batch and correlation among gene regions within each sample (see Methods for more details). XRE – xenobiotic response element, TSS – transcriptional start site.

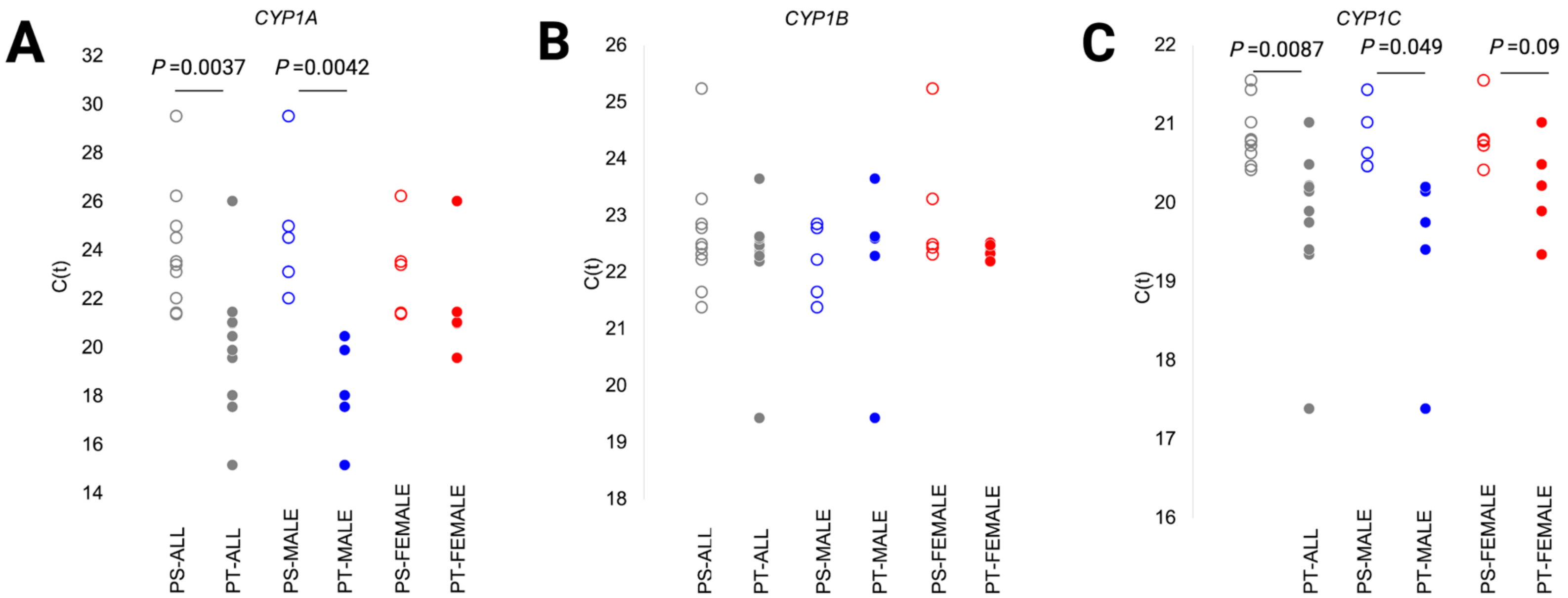


**Supplemental Figure 5. Expression of AhR target genes *CYP1A*, *CYP1B*, and *CYP1C* in adult PAH-sensitive and PAH-tolerant *F. heteroclitus*. (a)** Cycles to threshold (C(t)) values for *CYP1A* by population and sex. **(b)** C(t) values for *CYP1B* by population and exposure status. **(c)** C(t) values for *CYP1C* by population and exposure status. *P*-values are for independent samples t-tests comparing C(t) values between populations. PS- PAH-sensitive, PT- PAH-tolerant.

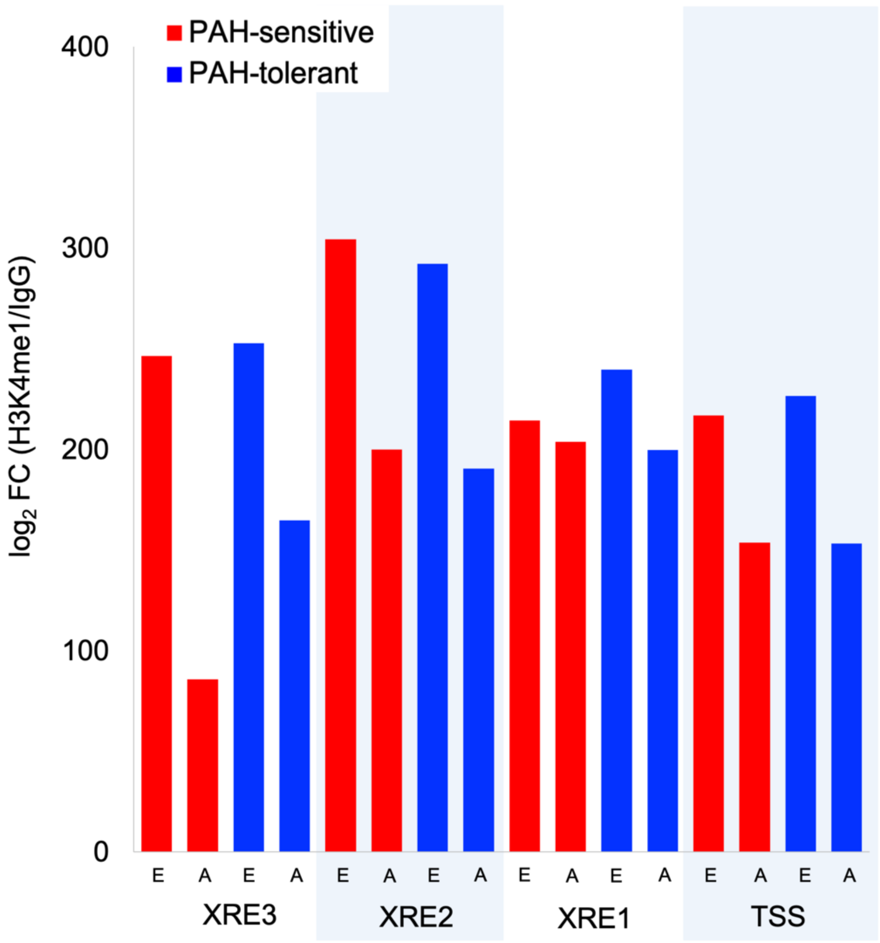


**Supplemental Figure 6. H3K4me1 enrichment at the *CYP1A* proximal promoter-enhancer in PAH-sensitive and PAH-tolerant *F. heteroclitus* embryos and adults with no PAH exposure.** Least squares means (back-transformed from inverse-transformed model output) histone post-translational modification (PTM) H3K4me1 enrichment in adult liver and embryos from PAH-tolerant and PAH-sensitive populations. Data shown for PAH-sensitive adults and embryos that never exposed to PAH. Data shown for adult PAH-tolerant fish with no current PAH exposure but with a history of extreme PAH exposure. Data are adjusted for experimental batch and correlation among gene regions within each sample (see Methods for more details). XRE - xenobiotic response element, TSS - transcriptional start site, E - embryos, A - adults.

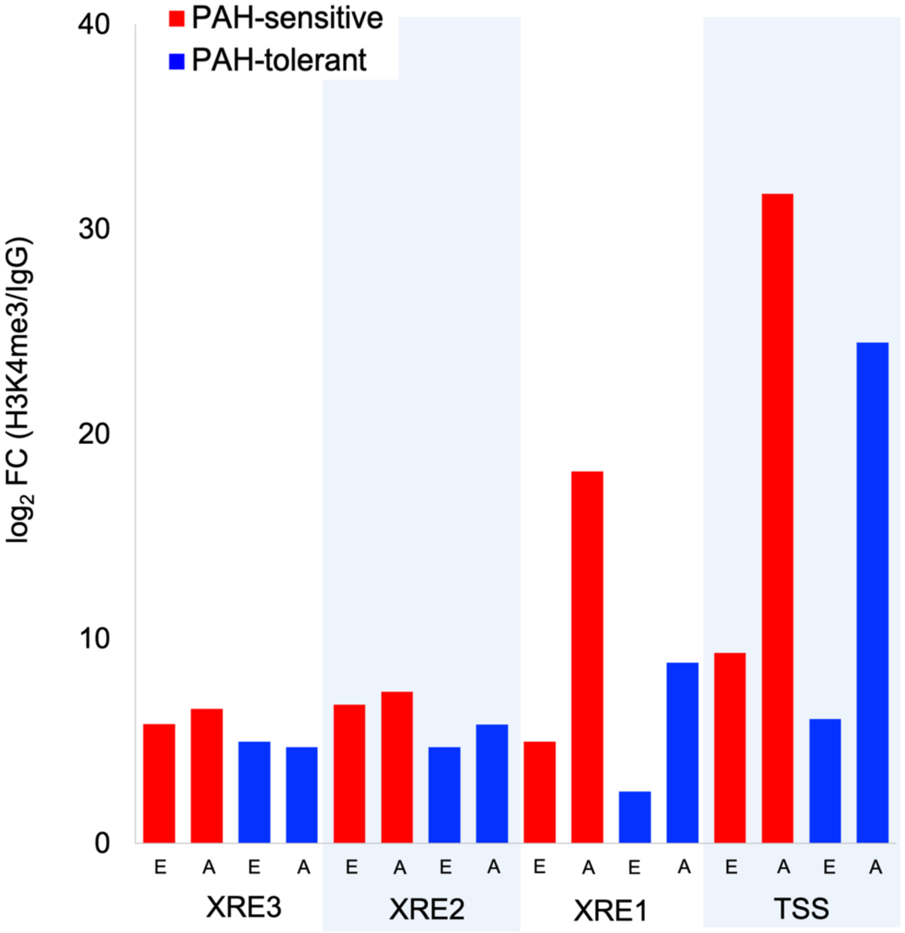


**Supplemental Figure 7. H3K4me3 enrichment at the *CYP1A* proximal promoter-enhancer in PAH-sensitive and PAH-tolerant *F. heteroclitus* embryos and adults with no PAH exposure.** Least squares means (back-transformed from inverse-transformed model output) histone post-translational modification (PTM) H3K4me3 enrichment in adult liver and embryos from PAH-tolerant and PAH-sensitive populations. Data shown for PAH-sensitive adults and embryos that never exposed to PAH. Data shown for adult PAH-tolerant fish with no current PAH exposure but with a history of extreme PAH exposure. Data are adjusted for experimental batch and correlation among gene regions within each sample (see Methods for more details). XRE - xenobiotic response element, TSS - transcriptional start site, E - embryos, A - adults.

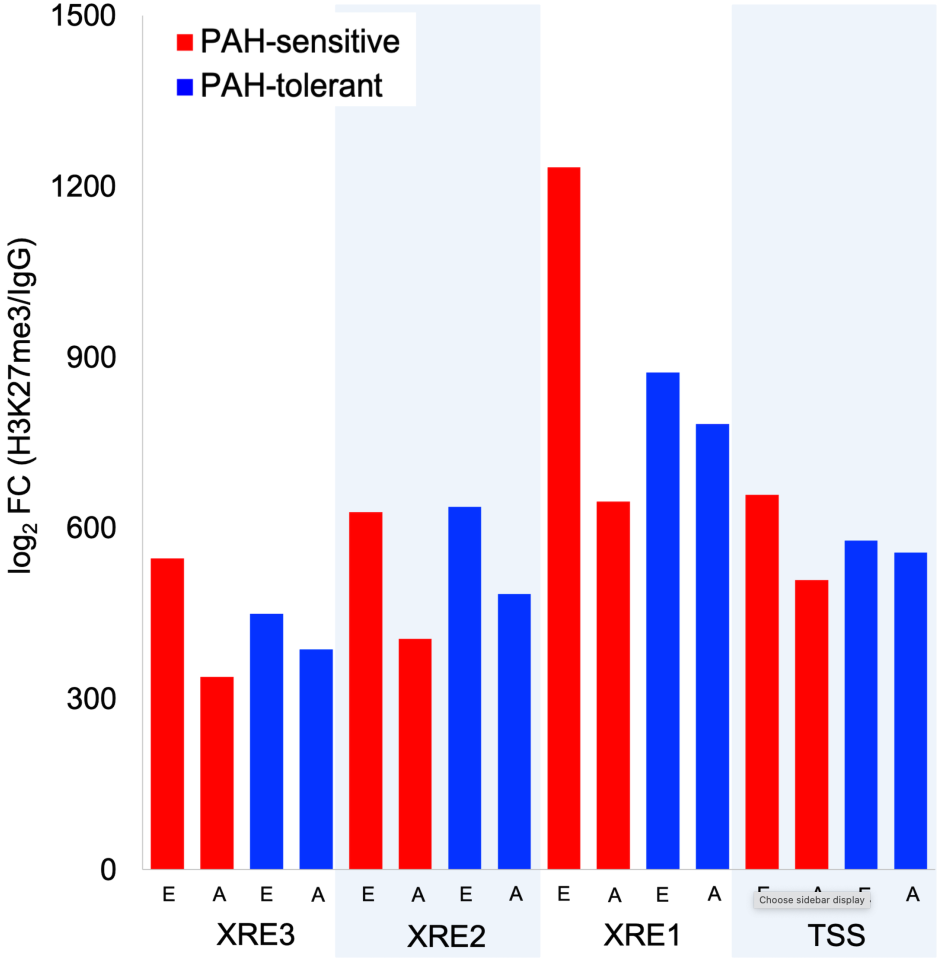


**Supplemental Figure 8. H3K27me3 enrichment at the *CYP1A* proximal promoter-enhancer in PAH-sensitive and PAH-tolerant *F. heteroclitus* embryos and adults with no PAH exposure.** Least squares means (back-transformed from inverse-transformed model output) histone post-translational modification (PTM) H3K27me3 enrichment in adult liver and embryos from PAH-tolerant and PAH-sensitive populations. Data shown for PAH-sensitive adults and embryos that never exposed to PAH. Data shown for adult PAH-tolerant fish with no current PAH exposure but with a history of extreme PAH exposure. Data are adjusted for experimental batch and correlation among gene regions within each sample (see Methods for more details). XRE - xenobiotic response element, TSS - transcriptional start site, E - embryos, A - adults.

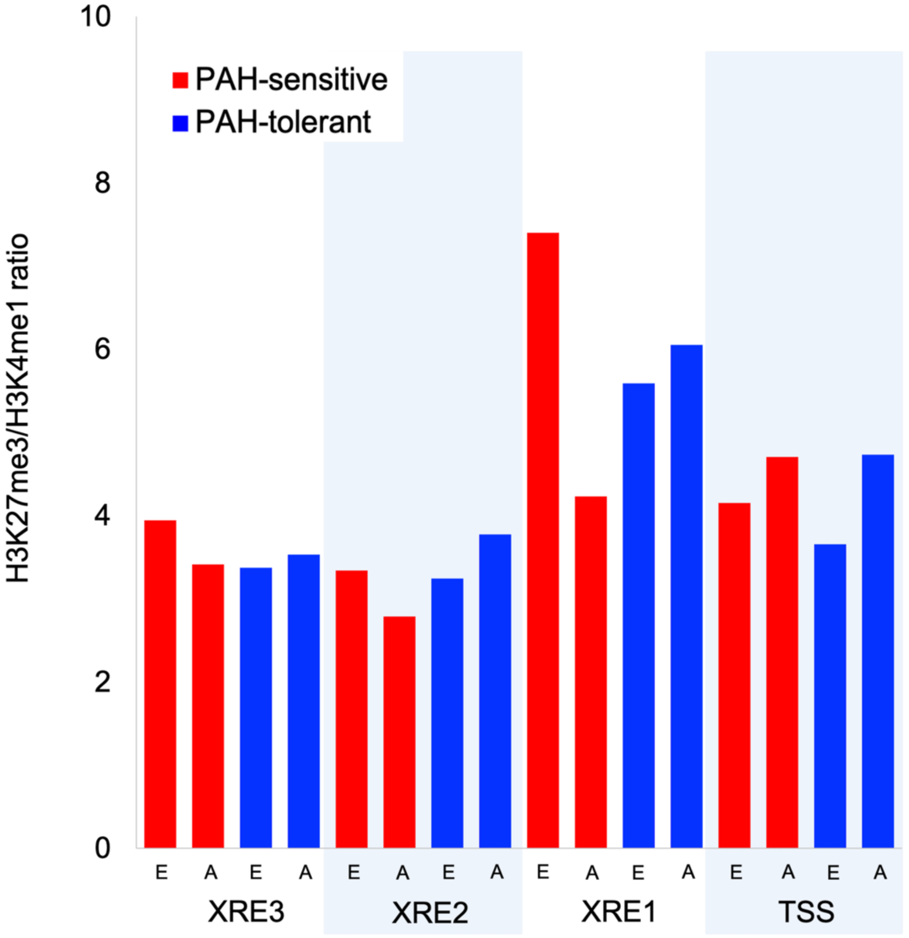


**Supplemental Figure 9. H3K27me3: H3K4me1 ratio at the *CYP1A* proximal promoter-enhancer in PAH-sensitive and PAH-tolerant *F. heteroclitus* embryos and adults with no PAH exposure.** Least squares means (back-transformed from inverse-transformed model output) histone post-translational modification (PTM) H3K27me3: H3K4me1 ratios in adult liver and embryos from PAH-tolerant and PAH-sensitive populations. Data shown for PAH-sensitive adults and embryos that never exposed to PAH. Data shown for adult PAH-tolerant fish with no current PAH exposure but with a history of extreme PAH exposure. Data are adjusted for experimental batch and correlation among gene regions within each sample (see Methods for more details). XRE - xenobiotic response element, TSS - transcriptional start site, E - embryos, A - adults.

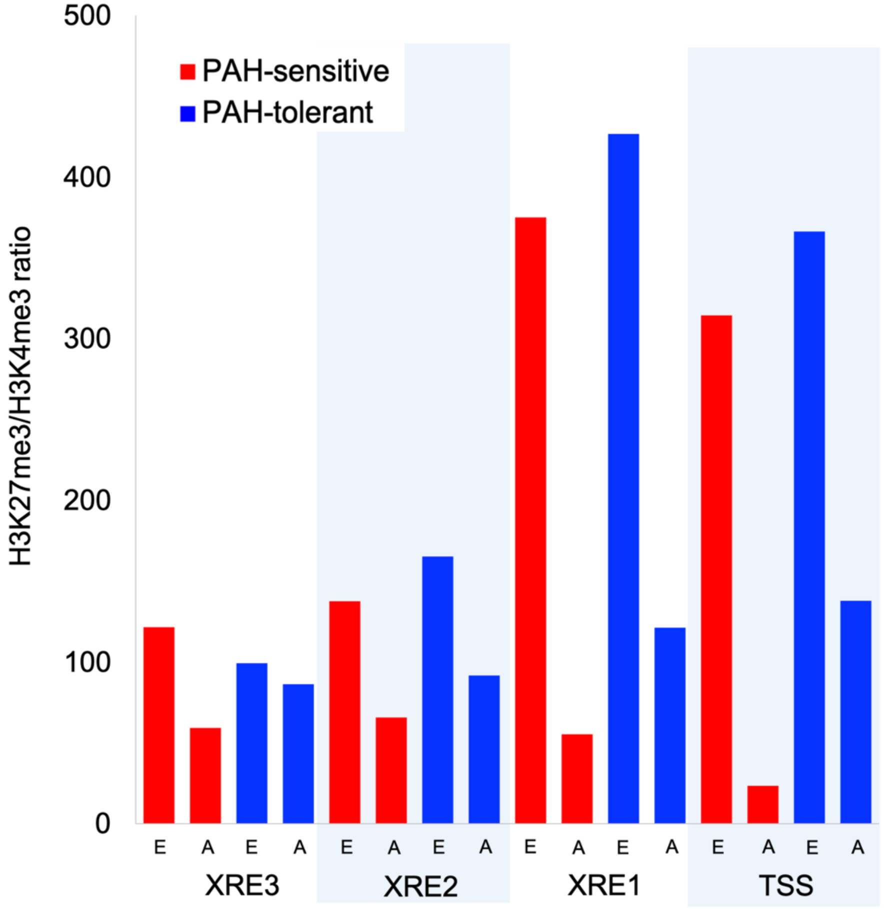


**Supplemental Figure 10. H3K27me3: H3K4me3 ratio at the *CYP1A* proximal promoter-enhancer in PAH-sensitive and PAH-tolerant *F. heteroclitus* embryos and adults with no PAH exposure.** Least squares means of (back-transformed from inverse-transformed model output) histone post-translational modification (PTM) H3K27me3: H3K4me3 ratios in adult liver and embryos from PAH-tolerant and PAH-sensitive populations. Data shown for PAH-sensitive adults and embryos that never exposed to PAH. Data shown for adult PAH-tolerant fish with no current PAH exposure but with a history of extreme PAH exposure. Data are adjusted for experimental batch and correlation among gene regions within each sample (see Methods for more details). XRE - xenobiotic response element, TSS - transcriptional start site, E - embryos, A - adults.

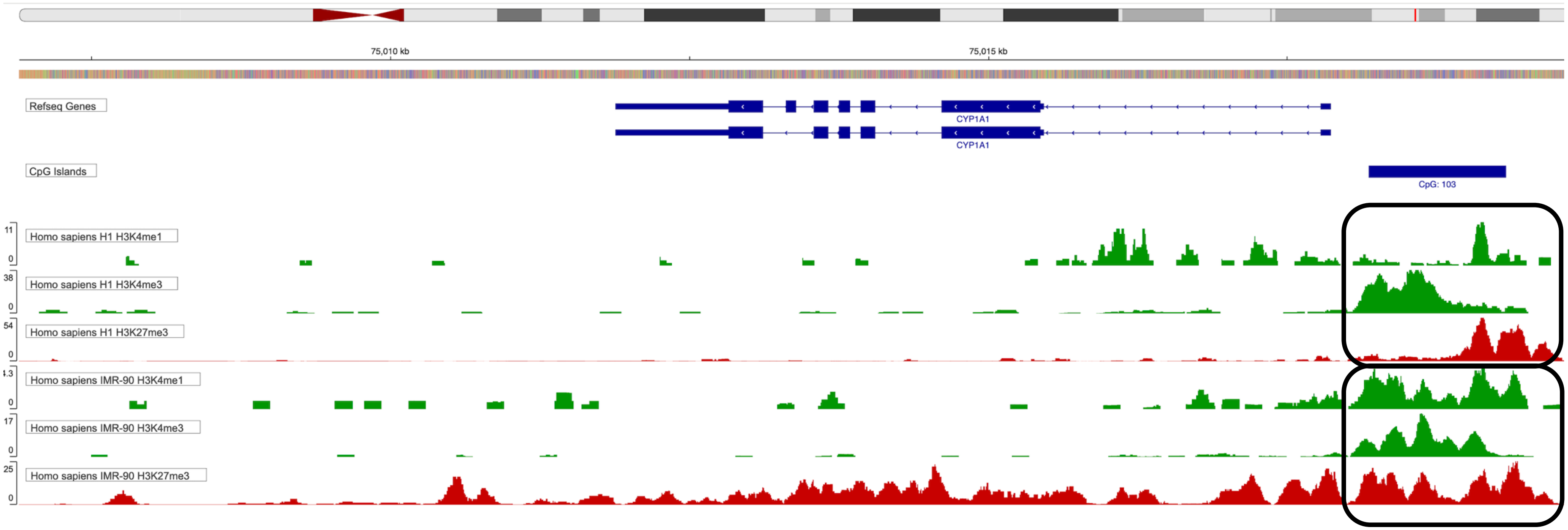


**Supplemental Figure 11. The human *CYP1A1* promoter contains a bivalent domain.** Human *CYP1A1* contains a bivalent domain, characterized by the presence of both repressive H3K27me3 and activating H3K4me1/3 (black boxes) in its proximal promoter-enhancer in both H1 embryonic stem cells and differentiated, PAH-target tissue IMR90 lung fibroblasts in the publicly available NIH ENCODE datasets.

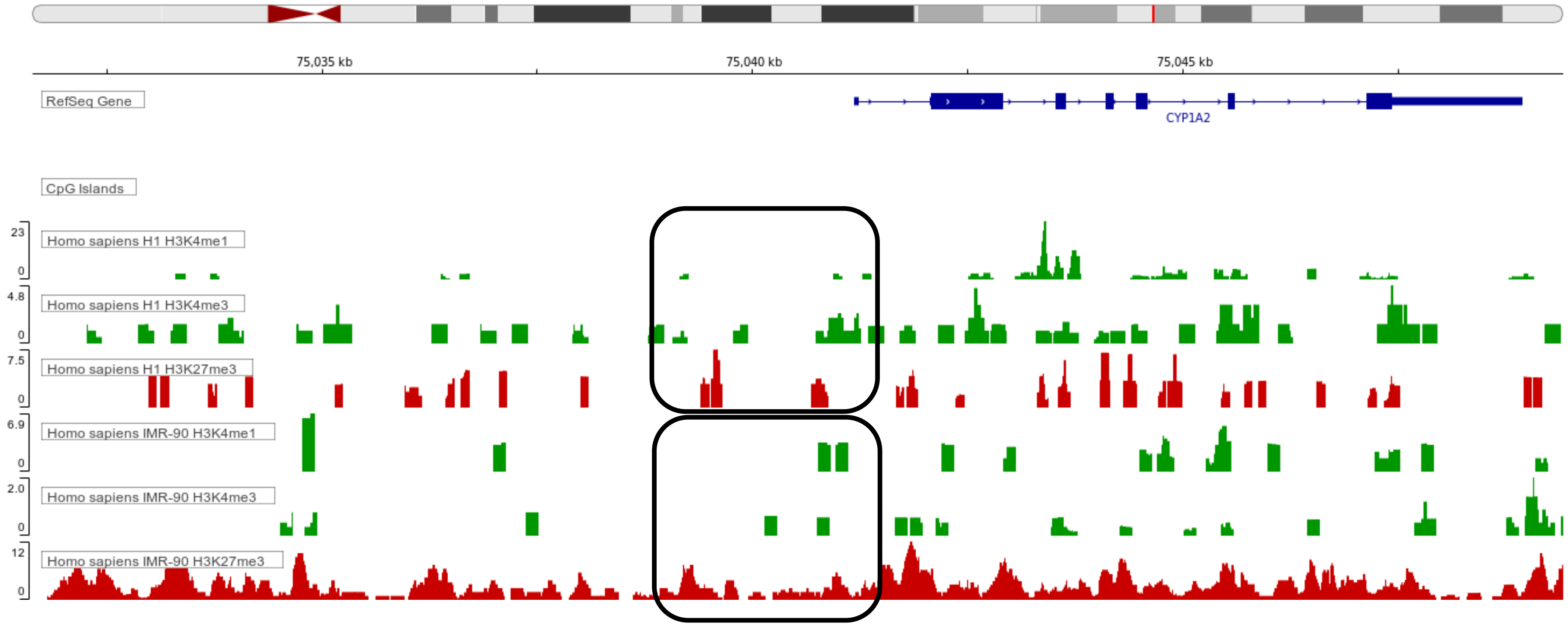


**Supplemental Figure 12. The human *CYP1A2* promoter does not contain a bivalent domain.** Human *CYP1A2* does not contain a bivalent domain in its proximal promoter-enhancer in either H1 embryonic stem cells or differentiated, PAH-target tissue IMR90 lung fibroblasts in the publicly available NIH ENCODE datasets.

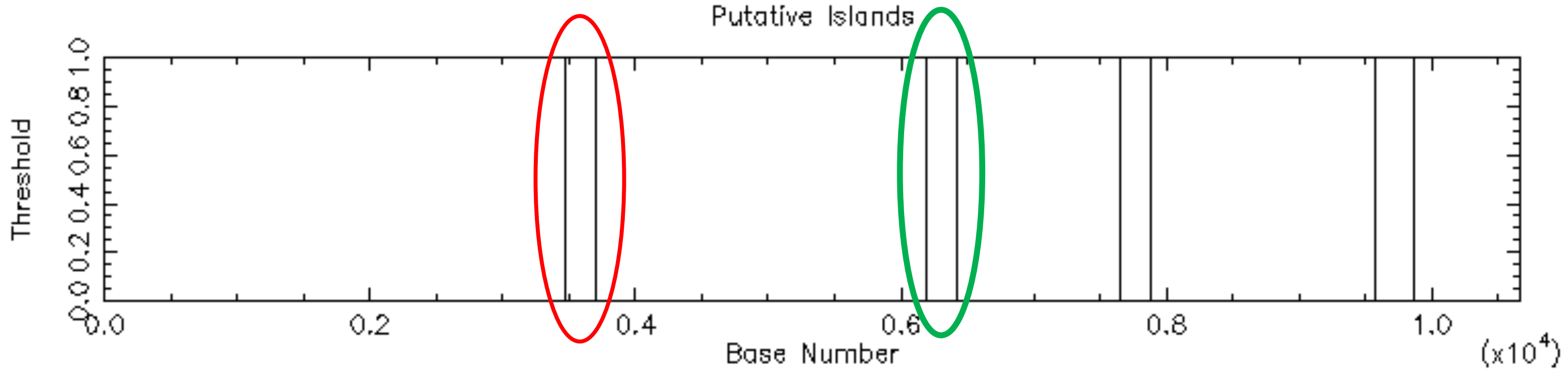


**Supplemental Figure 13. *CYP1A* in *Fundulus heteroclitus* contains two CpG islands in its promoter-enhancer region.** The CYP1A gene in Fundulus heteroclitus contains two predicted CpG islands, including one (shown in green) that contains two XREs (XRE3 and XRE2) that bind AhR to activate transcription in the presence of PAH. CpG islands were predicted using the EMBOSS CpGplot tool.
